# The developing mouse dopaminergic system: Cortical-subcortical shift in D1/D2 receptor balance and increasing regional differentiation

**DOI:** 10.1101/2024.03.05.583309

**Authors:** Ingvild E. Bjerke, Harry Carey, Jan G. Bjaalie, Trygve B. Leergaard, Jee Hyun Kim

## Abstract

The dopaminergic system of the brain is involved in complex cognitive functioning and undergoes extensive reorganization during development. Yet, these changes are poorly characterized. We have quantified the density of dopamine 1- and 2-receptor (D1 and D2) positive cells across the forebrain of male and female mice at five developmental stages. Our findings show a cortico-subcortical shift in D1/D2 balance, with increasing D1 dominance in cortical regions as a maturational pattern that occurs earlier in females. We describe postnatal trajectories of D1 and D2 cell densities across major brain regions and observe increasing regional differentiation of D1 densities through development. Our results provide the most comprehensive overview of the developing dopaminergic system to date, and an empirical foundation for further experimental and computational investigations of dopaminergic signaling.

## Introduction

Dopamine is a monoamine neurotransmitter modulating complex behaviors such as attention, cognitive flexibility, and decision making via its receptors^1–3^. Amongst five receptors, dopamine receptors 1 and 2 (D1 and D2) are the most abundant in the mammalian central nervous system^4^. D1 and D2 have distinct molecular structures and opposing intracellular processes^4^, which explains how dopamine can have such a varied role in cognition and affect^5^. These receptors also represent the main therapeutic targets for many psychiatric disorders, many of which are the most prevalent in adolescence^6^. Therefore, insight into the development of D1 and D2 distribution in the brain is essential for precision medicine.

Popular neurocognitive theories to explain adolescence as the dominant period for neuropsychiatric disorders posit that the function of subcortical socioemotional systems of the brain may develop faster compared to cortical cognitive control systems to result in reduced cognitive control during adolescence^7,8^. In addition, immature functional connectivity between cortical and subcortical regions during adolescence may further dampen cognitive control at this age^9,10^. The dopaminergic system is critical for neural connectivity^11^, and D1 and D2 signaling is particularly important for neural network function^12,13^. However, the postnatal development of D1 and D2 neurons has only been characterized for a few brain regions, and a comprehensive overview allowing for analysis of differences across the brain has been lacking.

To address this gap, we performed region-wise quantitative analysis of D1- and D2-expressing cells (hereafter referred to as D1 and D2 cells, respectively) in the entire forebrain of developing mice, using a comprehensive collection of immunohistochemically stained sections (DOPAMAP)^14^ that are spatially registered to age-specific Allen Mouse Brain atlases^15^. The brains available from the DOPAMAP collection (n = 153) span five age groups (postnatal day 17, 25, 35, 49 and 70) together covering development from the juvenile stage through adolescence, and both sexes. We created a custom scheme of anatomical regions largely following the hierarchical organization specified for the adult Allen Mouse Brain atlas^16^, but with cortical layers and very fine subregions merged. Immunolabelled cells were identified in section images using the image segmentation tool ilastik^17^ (Figure 1A-C), an open-source software for machine-learning based segmentation of features in biomedical images. The segmentation images were combined with the atlas maps using Nutil Quantifier^18^ (v0.8.0; Figure 1D), outputting the number of segmented objects (i.e. labelled cells) and coordinates representing them, sorted according to the atlas regions.

**Fig. 1.**
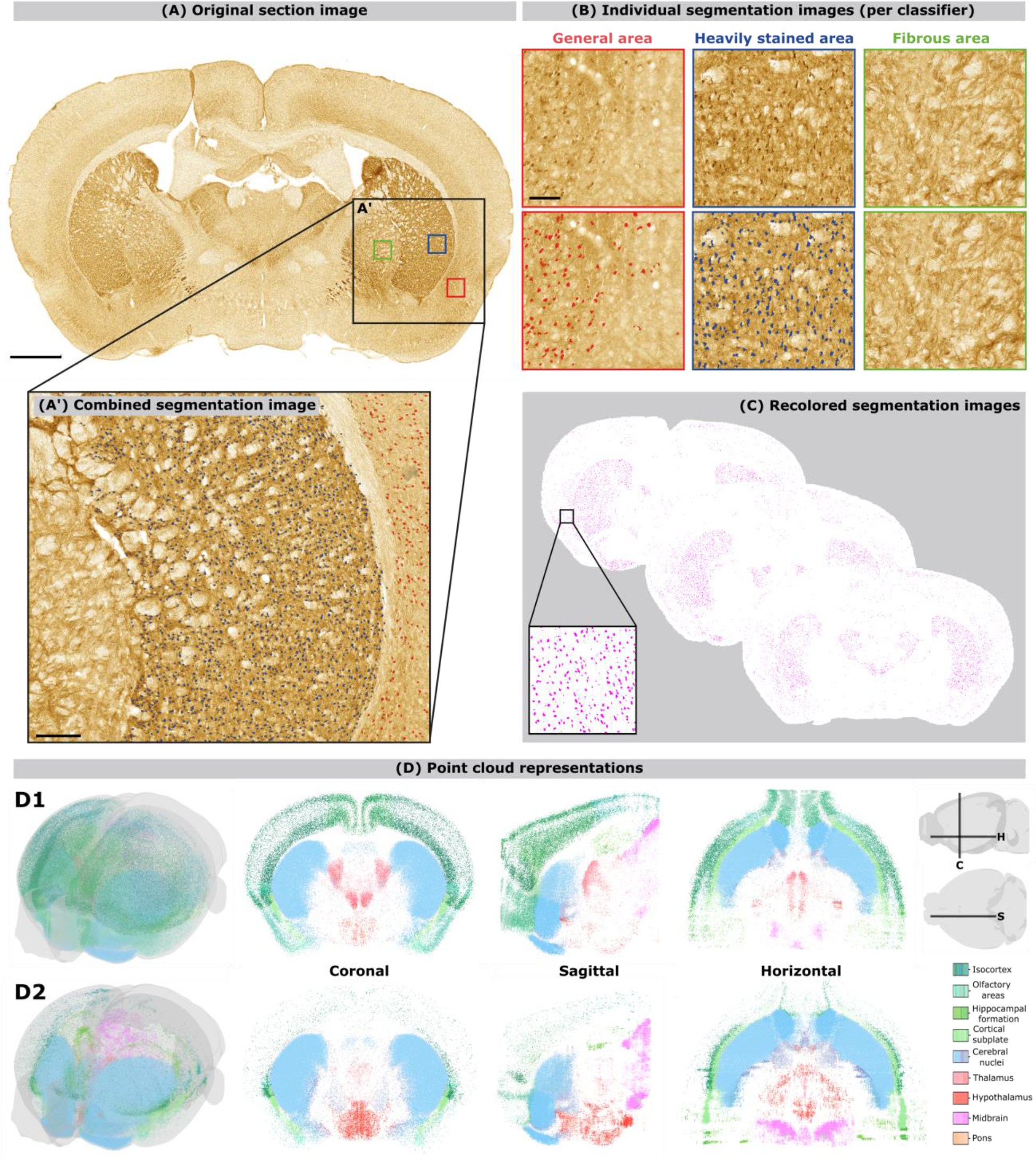
Forebrain-wide mapping of cells containing dopamine receptor 1 (D1) or 2 (D2). **(A)** Image of coronal section showing immunolabelled D1 cells (from subject C60 of the adult female group) that were segmented using a machine-learning based method. Because the level of staining appeared very different across areas, three separate algorithms were trained for different major regions defined using atlas delineations: one general, one for heavily stained areas, and one for fibrous areas (**B**). The resulting segmentations (**A’)** were combined based on the atlas regions (**C-D**). All objects in the segmentation images were extracted and quantified per region in the Allen mouse brain Common Coordinate Framework. (**E**) Point cloud representations showing the average expression of D1 (top) and D2 (bottom) cells based on all P70 and P49 subjects. The panels from left to right show a 3D view, as well as coronal, sagittal and horizontal slices through this volume. The individual points are color coded by the atlas regions. Scale bars are 2 mm, 200 µm and 100 µm in A, A’ and B, respectively.

The quantitative data were used for statistical analysis to investigate regional changes in D1- and D2 densities across development of female and male mice. Considering the importance of balance between D1- and D2-expressing cells in psychiatry^19–21^, we first asked whether D1 density normalized to D2 density (i.e., D1:D2 ratio^22^) show cortical-subcortical maturational differences. Raw D1 and D2 density values were then analyzed. With the cortical-subcortical maturation theory in mind, we investigated the maturation of D1 and D2 expression across cortical and subcortical regions and explored the correlation of D1 and D2 expression between regions across the age groups. We provide the quantitative data as an openly shared resource, which can be used to explore the data presented here, to answer questions beyond those asked in the current study or to build computational models.

## Materials and Methods

We used a comprehensive collection of openly shared datasets showing dopamine 1- and 2- receptor positive cells in the developing mouse forebrain^23–40^. These datasets consist of high-resolution microscopic images that are publicly available through the EBRAINS Knowledge Graph (https://search.kg.ebrains.eu/). The acquisition and sharing of these data are described in detail in previous publications^14,22^. We used the semi-automated QUINT workflow^41,42^ to extract labelled cells from the public images and perform quantitative analysis across the forebrain and through postnatal development.

### Overview of data

The DOPAMAP collection includes data from DR1a-EGFP and DR2-EGFP mice, with male and female subject from five age groups (P17, P25, P35, P49 and P70). Animals were sacrificed by perfusion fixation, and 40 µm thick coronal sections were cut and immunohistochemically stained using 3,3-diaminobenzidine (DAB). Series of sections were then scanned and shared as high-resolution microscopic images through the EBRAINS Knowledge Graph, with the full collection comprising 153 image series. Technical details on antibodies and protocols are given in Cullity et al.^22^

### Selection of data for analysis

Histological sections are vulnerable to damage during processing, such as tears or holes in sections; also, section images may display suboptimal staining. To select series for analysis, we evaluated whether the staining quality of each series was suitable for automatic extraction of cells. We also evaluated the completeness of each series, with regards to how much damage the sections displayed and how much of the brain a series covered; if a series displayed a lot of damage and covered only a small part of the brain, we did not include it in our analysis. This was largely a judgement call based on the cost of including a series with a lot of damage in the analysis (as such series requires significant additional time for non-linear registration and damage correction) and the added value the series would give to the analysis (which is higher the more of the brain is covered). An example series excluded based on tissue damage is series C36 from the D1 late adolescent male group ^29^. In this series, sections between section numbers 20 and 28 were missing, and sections posterior to s028 displayed considerable tissue damage. Thus, the posterior part of the series would likely have been excluded entirely. Given that several of the remaining sections also displayed damage and only covered the most rostral end of the forebrain, the series was excluded. Based on such considerations, we selected 111 out of 153 DOPAMAP series for semi-automatic, quantitative analysis of labelled cells. Table S13 summarizes the number of subjects in each group. Note that the series cover variable extents of the forebrain (for a full overview, see our previous publication), and that the number of animals in the D2 group with coverage through the posterior part of the forebrain was relatively low. However, the expression patterns in our individual and averaged datasets resembled those seen in other public datasets of D2 receptor expression from the Allen Institute (Figure S1).

### Spatial registration to atlas

The images in the DOPAMAP data collection were previously spatially registered to the Allen mouse brain Common Coordinate Framework (v3, 2017 versions of the delineations)^16^ using the QuickNII tool to perform linear transformation of the atlas to fit each section image^14^. For the youngest age groups (P17, P25 and P35), we used versions of the atlas that have been tailored to fit serial two-photon tomography templates of developing brains^15^. The overlays achieved by use of linear registration with QuickNII provide a good starting point for interpreting neuroanatomy, but do not fit perfectly due to the normal variation between animals and deformities introduced by the histological processing. To facilitate accurate automatic quantification of cells in different atlas regions, we therefore further refined the existing atlas overlay images by use of VisuAlign (RRID:SCR_017978), which enables in-plane non-linear transformations. When using VisuAlign, we first focused on fitting the atlas overlay to the outer edges of the section images, at the same time altering the placement of borders between cortical regions as little as possible. Secondly, we adjusted the placement of borders for clearly distinguishable landmarks deep in the brain, such as the hippocampus, striatum, and globus pallidus. We avoided adjusting the atlas overlay in regions where a neuroanatomical substrate for its borders could not be seen in the section.

### Image segmentation

Image segmentation was performed using ilastik, an open-source software for machine-learning based segmentation of features in biomedical images^17^. Prior to segmenting images, we downscaled the images to 20% of their original resolution using Nutil (v.0.6.0). Twenty of these downscaled images were selected (one image from each of the 20 subject groups) to use during training. Both in D1 and D2 stained material, there were large variations in the general appearance and intensity of staining in different brain areas. For example, striatal regions showed heavy staining with dense cell populations, while other areas showed almost no cellular staining but a lot of fibers. Because of this variability, it was not possible to train a single classifier to reliably identify cells across the whole brain while simultaneously ignoring fiber staining in areas with few cells. We therefore trained one general classifier, designed to segment cells across most brain areas, and two specific ones to segment cells in 1) brain regions with very dense cell populations (e.g. the caudoputamen, nucleus accumbens and olfactory tubercle) and 2) areas with a lot of fiber staining (e.g. the globus pallidus and all white matter tracts). The resulting three algorithms are referred to as classifiers for “general”, “heavily stained” and “fibrous” regions, respectively. Examples of the different types of regions and resulting segmentations are shown in Figure 1.

We used the pixel classification workflow to train each of the three classifiers, creating two classes (“cell” and “background”). Our training images were cropped to mainly show regions of interest for the classifier being trained, and using cropped images also served to make the ilastik software more stable. Example segmentations were made in all training images and across different brain areas. The resulting segmentation was continuously evaluated using the “Live update” function in ilastik. Training was stopped when a satisfactory result was obtained, and the classification did not change significantly with more training. Segmentation results were exported as Simple Segmentation (.png) images using the Batch Processing function, running each of the three classifiers on each of the 111 image series.

The result of the above procedure were three sets of segmentation images for each brain, one for each classifier. However, from each segmentation image series, only a subset of the image was of interest for the final quantification. That is, for each region in the brain, either the ‘general’, ‘heavily’ or ‘fibrous’ segmentation image would be used depending on the staining pattern. The assignation of classifier for each brain region was the same across all subjects and age groups but was different for D1 and D2 stained material as different regions show heavy staining in these data. A custom python script was used to combine the three segmentation images based on the region definitions from the non-linear atlas maps. For each segmentation image resulting from each of the three classifiers, the script masks every region except those for which the given classifier should be used, based on information contained in the .FLAT files exported from VisuAlign. It then combines the three masked images to create one image consisting of segmentations from the three different classifiers (Figure 1C). The segmentations resulting from each of the classifiers were exported using different colors for the cells (Figure 1B) to verify that the automatic combination was working correctly. However, prior to quantification with Nutil, all objects were recolored (Figure 1D) to allow extraction of all objects in a single run.

### Quantification of segmented objects

To quantify the segmented objects across the brain, segmentation images were combined with the atlas maps using Nutil Quantifier (v0.8.0)^18^.

Object splitting was turned off and the pixel cut-off was set to 4 pixels. We used a custom region scheme largely following the Allen Mouse Brain atlas hierarchy, but with cortical layers and very fine subregions merged. The custom regions excel file, compatible with Nutil Quantifier, is shared with the public datasets. Remaining Nutil parameters followed the default settings.

### Identification of damaged and missing areas

Although we excluded some series due to the overall staining quality or amount of damage (see “Selection of data for analysis”), many images within the selected image series still have some artifacts caused by the processing of histological tissue such as damage or suboptimal staining in parts of sections. To account for such artifacts, we manually inspected all analyzed sections. Whenever a section displayed damage or suboptimal staining, we created a custom “mask” in Adobe Photoshop that covered the affected area. The mask images were then automatically superimposed on the segmentation images using a custom python script. The result of this procedure is a segmentation image where the parts corresponding to region(s) with damage are covered by a black mask. Any object present in damaged areas are thus masked and effectively excluded from the analysis. To account for excluded areas in the quantification, we used numbers from the from neighboring sections or the contralateral side to interpolate data (see “post-processing of results” below).

To identify and quantify areas that were masked for each image, we ran the mask images through Nutil Quantifier (v0.8.0). We used the same parameters as for quantifying segmented objects except that object splitting was turned on (since a mask can overlap more than one custom region). The mask load (i.e. the size of the mask relative to the size of the custom region) was used during post-processing of results to account for damaged areas in the final quantitative data (see below).

In some images, part of the section was missing (e.g. a cortical hemisphere missing in the caudal part of the brain) with the consequence that the image was cropped to fit the dimensions of the remaining section parts. When the image is cropped, the corresponding atlas plate is also cropped, and will consequently be missing parts of or whole regions. Such missing regions will not be represented in the mask load. To amend this, we created a python script to generate, for each atlas plate, a corresponding uncropped atlas plate. By comparison of the number of pixels for a region in each cropped and uncropped atlas map pair, we estimated the percentage of each region that were missing in each section image. This “hidden mask load” was added to the mask load to give a total mask load accounting for missing and damaged parts of sections.

### Post-processing of results

Nutil Quantifier generates reports containing the counts of objects in custom brain regions, sorted per coronal section image. However, several post-processing steps are necessary in order to generate realistic neuron numbers, accounting for 1) the section sampling interval; 2) the tendency to over-estimate when extrapolating numbers to whole regions^43,44^; 3) any damage or suboptimal staining intensity in section images. We generated a python script to streamline the post-processing of Nutil Quantifier results into realistic cell numbers and densities. The steps included in this post-processing pipeline are outlined below.

First, we corrected for damaged areas in all the section-wise object counts by use of the mask load information. The total mask load was the sum of the damage mask load (extracted from the “Load” column of the section-wise Nutil reports on the mask quantification) and the hidden mask load (see above). Regions could be either partly or fully masked (> 90% of the region covered by a mask were considered fully masked, as a few pixels of the region could have been missed when creating the mask). In fully masked regions, numbers were interpolated by use of data from neighboring sections. Numbers for missing sections (indicated by a section number in the serial order missing) were also interpolated.

Partly masked regions were corrected using the data for the intact part within the section, by use of the following formula:

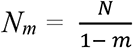

Where N is the raw number of objects, m is the mask load, and Nm is the mask-corrected number of objects.

Secondly, to account for the tendency to over-estimate object counts when extrapolating numbers observed in sections, we corrected the section-wise object counts using Abercrombie’s formula^43^:

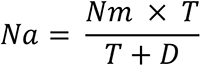

Where Nm is the mask-corrected object count, T is the thickness of the sections, and D is the mean diameter of the profiles, and Na is the resulting estimated number of cells (Abercrombie-corrected). The mean diameter of the profiles was calculated based on the size estimates in the Object reports from Nutil Quantifier. This report gives the estimated area in pixels for every object. We calculated the mean object size per custom region in µm by multiplying this by the pixel size (1.21), and converted this into the object diameter by the following formula:

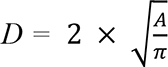

To obtain region areas in µm^2^, the section-wise region areas were multiplied by the pixel size of the images (which was 1.21 µm in all series). Lastly, to arrive at section-wise densities per region, the corrected object counts were divided by the region area. This gave the neuron count per µm^2^. To convert this into neuron counts per mm^3^, we divided it by the section thickness (40 µm; giving the count per µm^3^) and multiplied by 10^9^ (giving the count per mm^3^).

### Statistical analysis

For statistical analysis, we grouped the custom regions used in the QUINT analysis into 17 major brain regions. This was done to reduce the number of comparisons made and to increase the sample size for each area, making the analyses more robust. The sorting was guided by the hierarchy levels of the Allen Mouse brain CCF ontology. A full overview of the mapping of regions to this level of the hierarchy is given in Table S14, and an overview of the sorting of the fine-grained regions of the CCF into custom regions is shared with the public dataset^45^.

All statistics used IBM SPSS Statistics 29 (IBM Corp., NY, USA). Density data for each group for each brain region were first examined for normality (skewness and kurtosis of the distribution of individual datapoints). Most groups for each brain region fell within acceptable range (skewness range = -1.960 to 1.976; kurtosis range = -3.270 to 3.448)^46^, and these groups contained no statistical outliers (Grubb’s test p’s < 0.05). Groups with data that fell outside this range for skewness and/or kurtosis are shown in Table S15. Grubb’s test revealed an outlier from each of these groups, which were removed for subsequent analyses.

Group differences were then analyzed with two- or three-way analyses of variances (ANOVA) with Tukey’s post hoc multiple comparisons for age main effects and Bonferroni-corrected post hoc tests for any significant interactions (Table S1-2). For inter-regional correlations, two-tailed Pearson’s correlation (r) tests were used (Table S3-12).

### Visualization of data

We used the python matplotlib library to calculate descriptive statistics, summary data and D1:D2 ratios, as well as to create graphs used in the figures. All the code used for this is available through the DOPAMAP github repository and can be re-run on the data sheets provided in the public dataset^45^, to reproduce the figures presented in this paper or create similar figures for other datasets. The color-coding of D1:D2 ratio across major brain regions (Figure 2) and correlations coefficients (Figure 3) were created using conditional formatting in Microsoft Excel.

**Figure 2.**
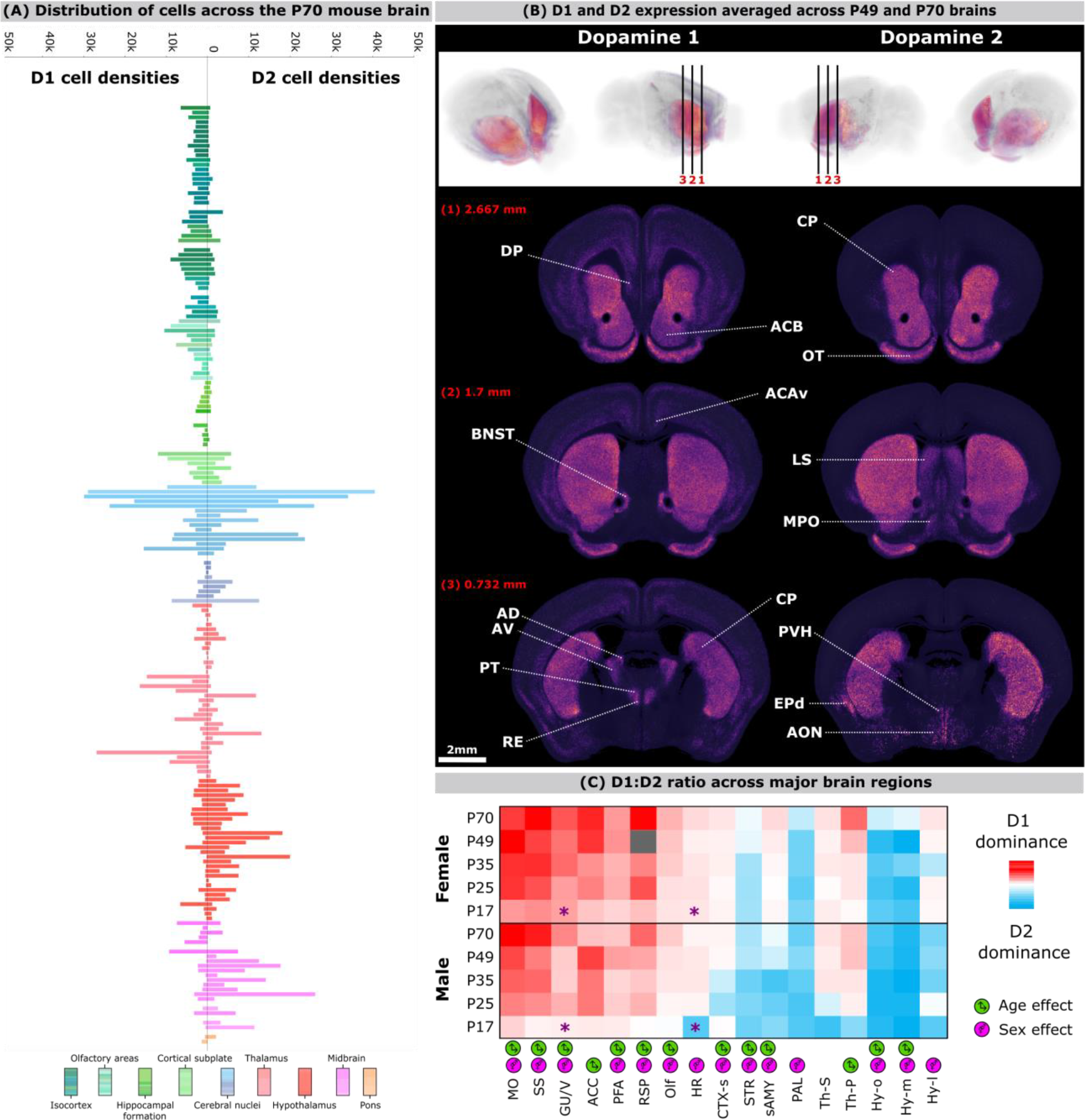
Distribution and ratio of dopamine receptor 1 or 2 expressing cells across the mouse brain and through development. (A) Bar plots of dopamine 1 (D1) and 2 (D2) cell densities across the P70 mouse brain, with D1 shown in the left panel and D2 in the right. Each bar represents a brain region and is color coded according to the Allen Mouse brain Common Coordinate Framework (bottom row) and sorted from top to bottom following the hierarchy. (B) 3D rendering (top) and coronal slice images (bottom) showing the intensity of D1 (left) and D2 (right) cell staining, averaged across P70 and P49 data to improve the visualization. Differences in the distributions can be seen corresponding to the data shown in panel A and are indicated by white lines. For example, D1 is more abundant in the cortex (e.g. in the anterior cingulate area (ACAv) and in deep layers across the cortex) and thalamus (e.g. anterodorsal (AD), anteroventral (AV), and reuniens (RE) nuclei), while D2 is more abundant in the lateral septum (LS), dorsal entopeduncular nucleus (EPd), and regions of the hypothalamus (e.g. anterior olfactory (AON) and paraventricular hypothalamic (PVH) nuclei). These intensity volumes can be explored interactively in the EBRAINS interactive atlas viewer; physical coordinates from the viewer of the levels shown here are indicated with red text ^47,48^. (C) Ratios of D1 and D2 cell densities (D1 divided by D2) in 17 major brain regions across development, with male and female ratios shown separately. Red and blue colors indicate strong D1 and D2 dominance respectively, while white indicates a balanced ratio. Grey indicates missing data. Significant *post-hoc* tests for age-specific sex effects following a significant age × sex interaction are indicated by an asterisk for the relevant age group. **Abbreviations:** MO, motor areas; SS, somatosensory areas; GU/V, gustatory and visceral areas; ACC, anterior cingulate areas; PFA, prefrontal areas; RSP, retrosplenial areas; Olf, olfactory areas; HR, hippocampal region; CTX-s, cortical subplate; STR, striatum; sAMY, striatum-like amygdalar nuclei; PAL, pallidum; Th-s, thalamus, sensory-motor cortex related; Th-P, thalamus, polymodal association cortex related; Hy-o, hypothalamus, other; Hy-m, hypothalamic medial zone; Hy-l, hypothalamic lateral zone.

**Figure 3.**
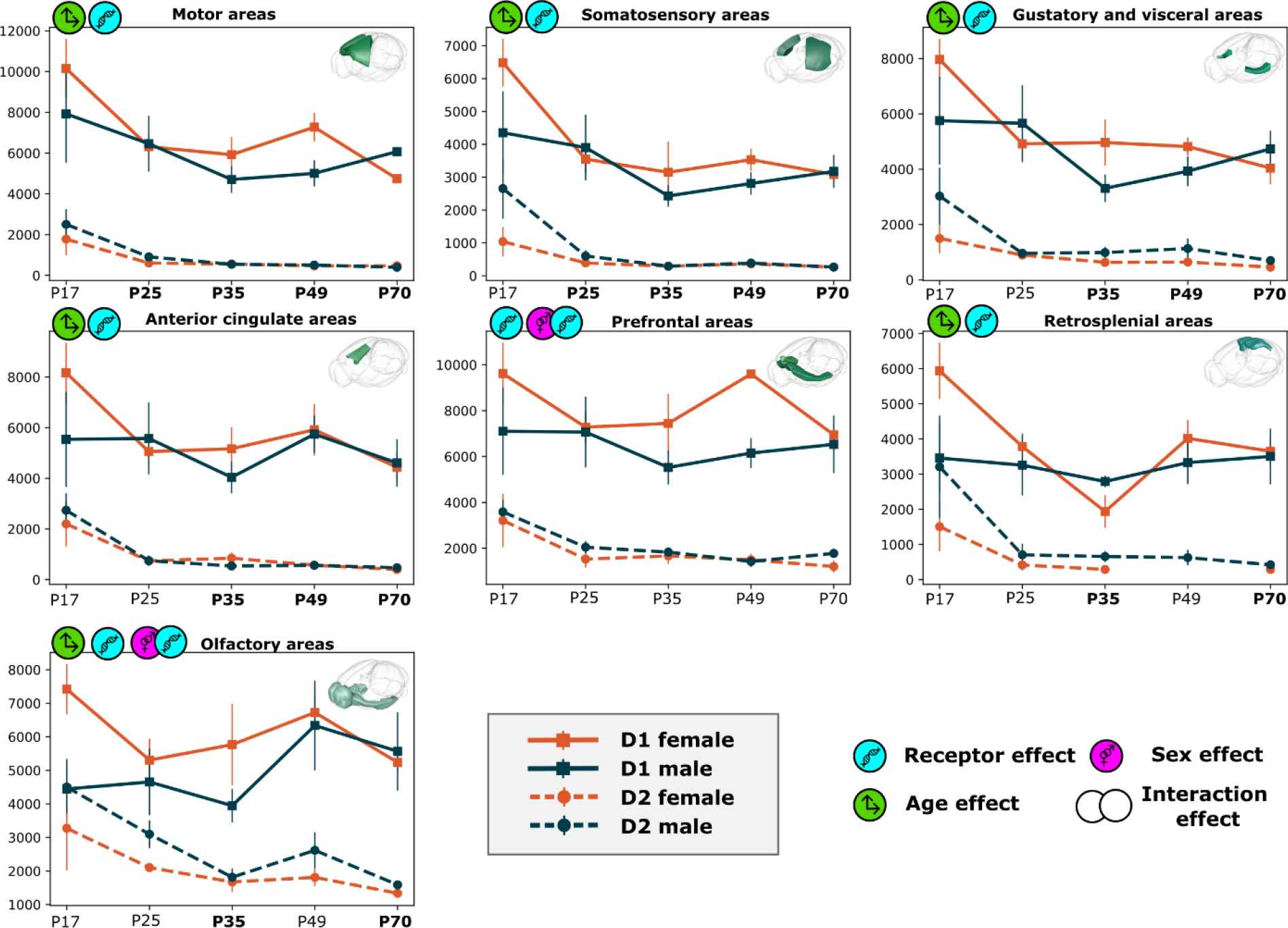
Postnatal trajectories of D1 and D2 cell densities in cortical regions. The line graphs show the development of D1 (solid lines) and D2 (dashed lines) for female (orange lines) and male (blue lines) subjects. Significant effects are indicated on the graphs, and age groups that were significantly different from P17 are bolded on the x axis. Error bars indicate SEM.

To create an average representation of the D1 and D2 neurons across the brain we used the .json files with coordinates from Nutil (3D_combined.json file, available for each subject in the shared dataset^45^). For each subject, this file contains the 3D atlas coordinates for all segmented pixels. Since prior analysis had shown there were no substantial differences between the two oldest age groups (P49 and P70), subjects from both these groups were combined for the average visualization. For each brain we created an ‘individual intensity volume’. This was achieved by first generating a volume corresponding to the Allen Mouse brain Common Coordinate Framework, with 25-micron voxel resolution. In this volume, each voxel was given a value corresponding to the number of segmented pixels with coordinates falling inside that voxel, as extracted from the Nutil .json file. To fill the space between sections, each of these volumes were linearly interpolated. To give the most accurate average representation of the individual intensity volumes, we wanted to weigh values by their distance to observed data (i.e. values close to observed data-points should contribute more to the average representation than values which were interpolated and far from any observed data-point). To achieve this, we created a second volume for each brain which contained the distance of each voxel from the nearest section, giving large values to voxels which were far from any section. Weighting values for all the voxels were then calculated so that they would exponentially decrease in relation to their magnitude, creating a ‘weights volume’ for each brain. The individual intensity volumes were then averaged together, with values in each volume weighted by the corresponding voxel in the weights volume. This process was done separately for D1 and D2 animals, creating average D1 and D2 volumes that were saved in both nifti and neuroglancer file formats. The NIfTI files were shared with the public dataset^45^, while the Neuroglancer format volumes were made available through Siibra, enabling users to interact with the data alongside atlas delineations. We used the interactive viewer when creating Figure 2, with the parameters set as follows:

- Opacity: 0.9
- Lower threshold: 0
- Higher threshold: 0.51 for D1 and 0.34 for D2. The higher threshold was set differently in the two volumes in order to normalize them to the highest intensity overall, which was found in the D2 volume, thus facilitating side-by-side comparison in the figure.
- Brightness: 0.08
- Contrast: 0
- Color map: magma

The average expression volumes were used to create an average point cloud representation to represent the D1 and D2 data (shown in Figure 1). In these volumes, points were placed according to the expression distribution represented in the average volumes, until the number of points was equal to the total number of cells estimated in our calculations from the QUINT data.

### Data sharing

All the data from the current study was curated and shared as a dataset via the EBRAINS Knowledge Graph^45^. The dataset contains all the necessary files to re-run any part of the analysis performed here, or to re-use parts of the dataset (e.g. the nonlinear registration information, segmentation images) in new analyses. The content and structure of the shared data is fully described in the data descriptor accompanying the dataset on the EBRAINS Knowledge Graph.

## Results

### Cortical-subcortical shift in dopamine receptor 1 dominance to dopamine receptor 2 dominance

Bar charts of D1 and D2 densities averaged across male and female mice of five age groups showed a marked shift from D1 dominance in cortical regions to D2 dominance in subcortical regions (Figure 2A). The cortical-subcortical shift in D1 vs D2 balance was also clearly seen when inspecting 3D representations of the average expressions (Figure 2B) and the D1:D2 ratio across regions (Figure 2C). For statistical analyses we grouped regions into 17 major brain regions, of which names and abbreviations are listed in the legend of Figure 2.

Two-way (age and sex factor) analyses of variances (ANOVA) with Tukey’s post hoc multiple comparisons for age main effects and Bonferroni-corrected post hoc tests for any significant interactions (Table S1, Figure 2C) showed that there were main effects of age in all cortical regions (MO, SS, GU/V, ACC, PFA, RSP, Olf; see Figure 2 legend for abbreviations), with P17 showing the least and P70 generally showing the most pronounced D1 dominance (except for motor and prefrontal areas, where D1 was most dominant at P49). The D1 dominance also varied with sex. In all cortical regions except for anterior cingulate areas, there were significant main effects of sex with females showing a stronger D1 dominance than males. Given that the strengthening of D1 dominance in the cortical regions is an overall maturational pattern, this indicates an earlier maturation of cortical regions in females. Significant age × sex interactions were also observed for D1 dominance in motor and gustatory and visceral areas. While follow-up *post hoc* tests indicated no sex effects at any age for motor areas, for gustatory and visceral areas there was a significant sex effect at P17 with females showing higher D1 dominance than males (t=2.504, p=0.020).

D2 dominance was seen in most subcortical regions (HR, CTX-s, STR, sAMY, PAL, Th-S, Th-P, Hy-o, Hy-m, Hy-l; see Figure 2 legend for abbreviations) except for the hippocampal region, pallidum, and hypothalamic lateral zone. In these region significant main effects of age were seen, with P17, P25, P35, and/or P49 being more D2 dominant than P70 (Table S1, Figure 2C). Significant main effects of sex were also observed in every subcortical region with males having stronger D2 dominance, except for the thalamus (Th-s, Th-p). Significant age × sex interactions were observed in the hippocampal region and hypothalamic medial zone. Follow-up *post hoc* tests indicated a significant sex effect at P17 with males showing stronger D2 dominance in the hippocampal region (t=3.657, p<0.001), whereas no sex effects were observed at any age for hypothalamic medial zone. Consistent with cortical regions, the results indicate an earlier maturation of subcortical regions in females with the weakening of D2 dominance an overall maturational pattern in the subcortical regions.

### Postnatal trajectories of dopamine receptor cells across the forebrain

To understand the independent changes in D1 and D2 density across age and sex, these values were analyzed with three-way (receptor, age, and sex factors) ANOVAs with *post hoc* tests as described above (Figure 3-4). Main effects of receptor were significant in all cortical regions, with higher D1 than D2 density (Table S2). All cortical areas (except the prefrontal areas) also showed a main effect of age, with the P17 group showing the highest overall receptor densities. Specifically, P17 showed higher receptor densities compared to all other ages in motor and somatosensory areas, compared to P35-P70 in gustatory and visceral, anterior cingulate, and retrosplenial areas, and compared to P35 in olfactory areas (Figure 3). In prefrontal and olfactory areas, significant sex × receptor interactions were observed. Follow up *post hoc* tests revealed a significant sex effect for D1 (t=2.655, p=0.010) but not D2 in olfactory areas; no sex differences were detected for any receptor for the prefrontal areas.

**Figure 4.**
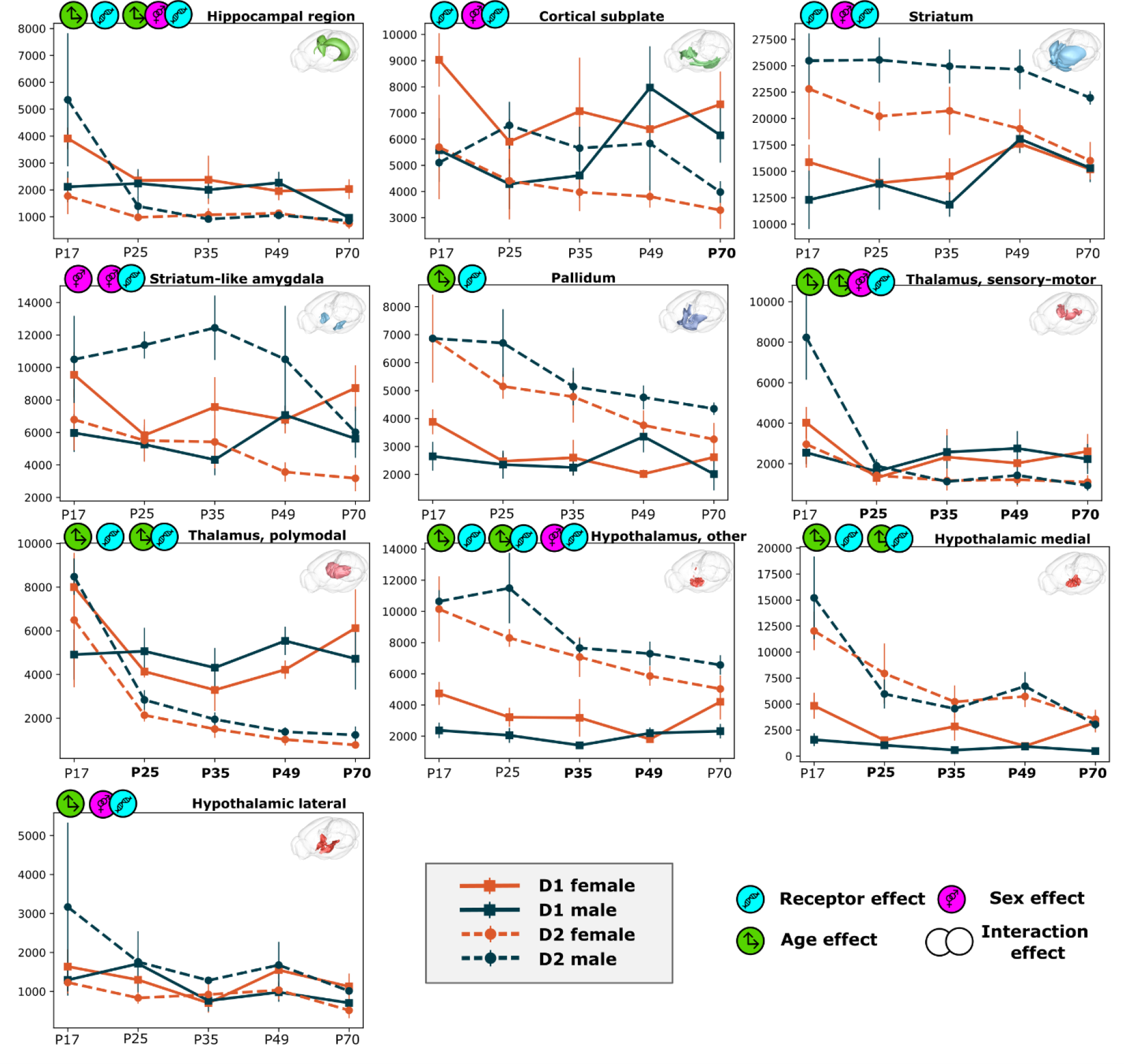
Postnatal trajectories of D1 and D2 cell densities in subcortical regions. The line graphs show the development of D1 (solid lines) and D2 (dashed lines) for female (orange lines) and male (blue lines) subjects. Significant effects are indicated on the graphs, and age groups that were significantly different from P17 are bolded on the x axis. Error bars indicate SEM.

Most subcortical regions also showed significant differences between receptor types (except for striatum-like amygdalar areas, the sensory-motor cortex related thalamus, and hypothalamic lateral zone; Table S2). These effects of receptor type were more complex than in the cortical regions, interacting with one or more factors (age and/or sex) in all regions, except for the pallidum (where the D2 density was higher than D1 density regardless of age and sex (Figure 4)). Age effects were observed in all subcortical regions except for cortical subplate, striatum, and striatum-like amygdalar nuclei. All the age effects interacted with receptor and/or sex, except in the pallidum (where post hoc test indicated higher overall receptor levels in P17 than P70) and hypothalamic lateral zone (post hoc tests were not significant).

Follow up post hoc tests to sex × receptor interactions showed that females had a higher D1 density than males in cortical subplate, striatum-like amygdalar nuclei, and hypothalamus other areas (biggest p = 0.040), while there were no sex effects on D1 density in striatum and hypothalamus lateral areas. Males had a higher D2 density than females in striatum, striatum-like amygdalar nuclei, and hypothalamic lateral zone (biggest p = 0.049), while there were no sex effects on D2 density in cortical subplate and hypothalamus other areas.

Follow up post hoc tests to age × receptor interactions showed that in thalamus polymodal area D2 density is significantly higher in P17 compared to all the other ages (biggest p<0.001), while D1 density was not affected by age. Similar analyses in hypothalamus other area showed that D2 density in P17 is higher than in P35-P70 and P25 is higher than P70 (biggest p=0.036), but no age effects in D1 density. In contrast, hypothalamic medial zone D1 density was significantly higher in P17 compared to P49 while D2 density was higher in P17 compared to all the other ages.

Thus, while cortical D1 and D2 densities (Figure 3) were generally similar in male and female mice with age effects not dependent on receptor or sex, several subcortical areas showed receptor-specific sex and age differences (Figure 4). Interestingly, we observed a three-way interaction effect (age × sex × receptor) in the hippocampal region and sensory-motor cortex related thalamus. For both areas, the interaction was driven by D2 density that showed age x sex interactions (biggest p = 0.023), with Figure 4 indicating particularly high levels of D2 in P17 males compared to all other groups.

### Age-specific correlation of dopamine receptor 1 or 2 density between regions

For each age group, we asked whether the cell densities correlated across different areas within the same receptor type (Table S3-S12). We found a striking reduction in correlation of D1 cell densities across regions with age. In P17, almost all region pairs (116 out of 136, or 85 %) showed a significant positive correlation (Figure 5), and the correlation coefficients were generally higher within broader hierarchical groups. For example, the density in cortical motor areas was highly correlated with that of the somatosensory areas (r(14) = 0.91, p < 0.001), and slightly less correlated with that in the hippocampal region (r(14) = 0.70, p = 0.003). The correlation coefficients gradually decreased with age (Figure 5), so that at P70, there were no significant correlation between the cortical motor areas and these regions (somatosensory areas, r(7) = 0.36, p = 0.344, hippocampal region, r(7) = -0.44, p = 0.241); in general, only 31% of region pairs showed a significant positive correlation at this age. The D1 density in the hypothalamic lateral zone was in fact significantly negatively correlated with that in the striatum-like amygdalar nuclei at P35 (r(10) = -0.60, p = 0.04) and P49 (r(9) = -0.70, p = 0.016), and the sensory-motor cortex related thalamus at P49 (r(8) = -0.63, p = 0.049). Although the D1 and D2 density values for some other region pairs were negatively correlated across the dataset (Figure 5), only those mentioned above were significant. In contrast, D2 densities were generally less correlated between regions than D1 densities, with adolescent periods of P35 and P49 showing the most significant positive correlations (42% and 38% of region pairs, respectively). Although some negative correlations were seen in almost all groups, very few were significant (D1 densities for the hypothalamic lateral zone with sensory-motor cortex related thalamus at P35 and P49, and with striatum-like amygdala nuclei at P49).

**Figure 5.**
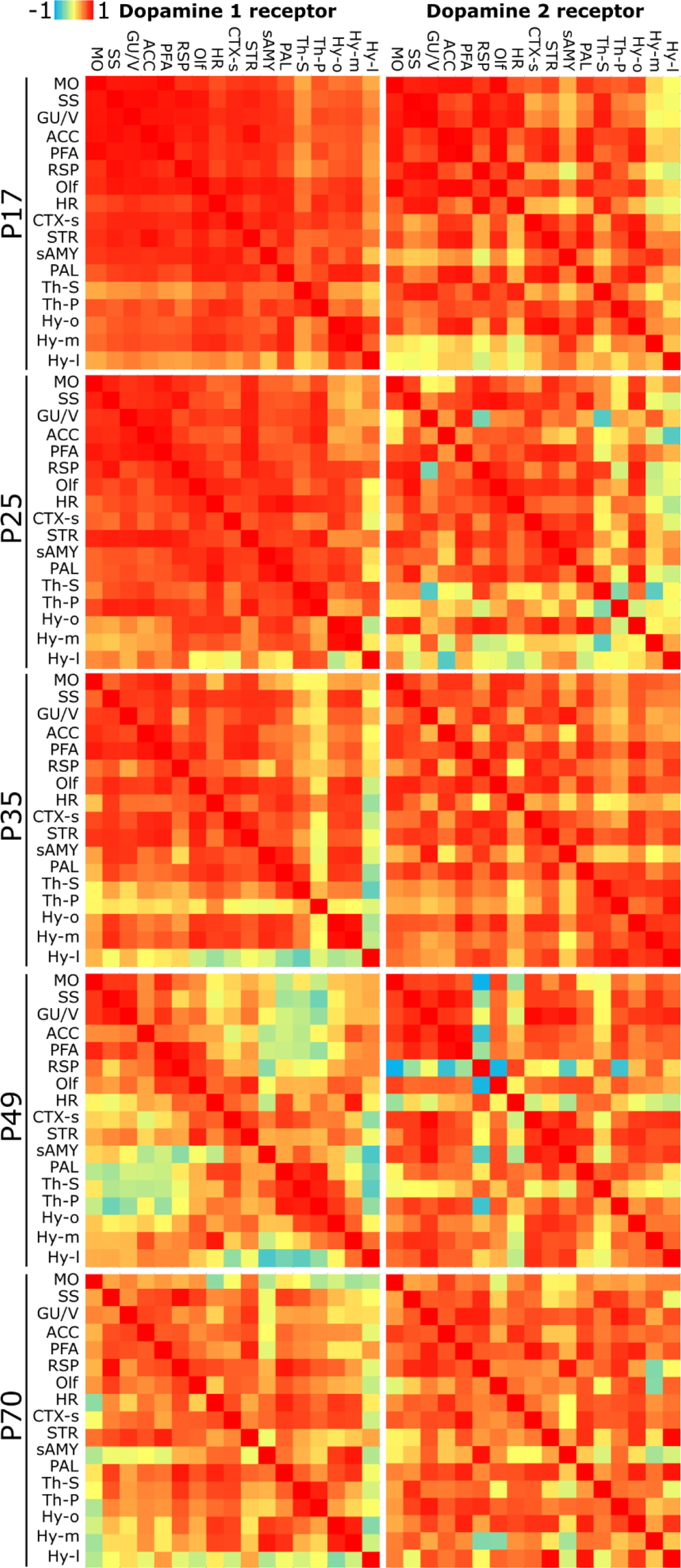
Correlation matrices showing the correlation of D1 (left) and D2 cell densities (right) across 17 major forebrain regions. D1 densities are highly correlated across regions within animals at P17, with correlation coefficient gradually decreasing through development. At P17, 85% region pairs show a significant positive correlation, compared to 77%, 54%, 24% and 31% at P25, P35, P49 and in the adult, respectively. D2 densities are generally less correlated than D1 densities, with 35% of region pairs showing a significant positive correlation at P17, 18% at P25, 42% at P35, 38% at P49 and 22% at P70.

## Discussion

We quantified D1 and D2 cell densities in atlas-defined regions in the forebrain of male and female mice across five age groups from the juvenile stage to young adulthood. While previous studies have investigated D1 and D2 expression in one or a few areas at a time, either through cell counting^22,49,50^ or other methods^51,52^, the current study is the first to cover the entire forebrain. The present results show considerable differences in D1 and D2 densities across age groups, sex, and regions. While the cortical-subcortical maturation theory might suggest that D1 and D2 expression should stabilize in subcortical regions before cortical regions, we did not find such a pattern in our dataset. Most of the complex age-related interaction effects were found in subcortical areas, while cortical areas showed simpler age main effects with P17 expressing the most D1 and D2. In addition, D1 expression between regions correlated less with age, a surprising finding that may indicate increasing differentiation with age. D1 may be upregulated with relatively low selectivity early in life, but with maturation its role appears to be highly specialized and dissociated across regions. The highest regional correlations in D2 were observed during adolescence. This may reflect alterations in the relative amounts of D2 across the brain in this period, a period where D2 expression levels are prone to dynamic changes in response to stressful events^53–55^.

D1 and D2 cell densities typically declined from P17 onwards in most regions, indicating pruning of dopamine receptors in this period. Such pruning has been shown in several brain areas before, but the extent and timing might differ amongst areas^56,57^. The difference of the P17 group from the rest of the ages is highly interesting. One of the best characterized behaviors in P17 rodents is extinction of conditioned fear, which refers to when a cue that used to elicit a threat response no longer elicits a response because it has been repeatedly presented without any consequences. It represents a new learning that confers a neutral meaning to the cue that competes with the cue-threat association. Extinction is generally reversible from three weeks of age and the threat response to the cue can relapse. However, extinction is irreversible in P17 male rodents, which is attributed to extinction erasing conditioned fear^58,59^. Importantly, extinction is reversible in P17 female rodents^60,61^, which is consistent with the present study showing a more mature D1 and D2 expression than males at this age. As D1 and D2 cells are crucial players in mechanisms of learnt fear^62,63^, one might speculate that high levels of D1 and D2-expressing cells in P17 mice play a role in the unique extinction capabilities at this age.

D2 expression was especially irregular across sex and age in subcortical areas, where D2 cells were generally more abundant than D1 cells. Alterations in the D2 system are associated with several psychiatric disorders^64,65^, and many medications in psychiatry exert their effect through modulation of D2 activity^66,67^. However, vulnerability to disorders and effect of drugs is often sex- and age-specific^6,68,69^. In this light, our finding of a higher D2 density in males than females early in life is particularly interesting, and may indicate important new targets for intervention studies for sexually dimorphic psychiatric disorders^70^.

Our study provides a comprehensive overview of the number of D1 or D2 positive cells in the developing mouse forebrain. Both the raw and derived data are made available through the EBRAINS Knowledge Graph and provide a starting point for new analyses or generation of hypotheses. The derived data include point clouds that can be integrated in computational models, where the current data should be of high interest for modelling the basal ganglia. Thus, we believe our study provides an important advance to our knowledge of the developing dopaminergic system and that the comprehensive data provided with it will serve as an important resource of broad interest for neuroscientists.

## Acknowledgments

We thank Gergely Csucs, Nicolaas Groeneboom, Maja Puchades and Sharon Yates for expert technical assistance and help with use of the tools of the QUINT workflow, and Xiao Gui for technical expertise and customization of the Siibra viewer to visualize the presented data.

## Author contributions

Conceptualization: IEB, TBL, JHK Methodology: IEB, HC, JHK Investigation: IEB, JHK Visualization: IEB, HC

Funding acquisition: JGB, TBL, JHK Project administration: IEB, JHK Supervision: TBL, JHK

Writing – original draft: IEB, JHK

Writing – review & editing: IEB, HC, JGB, TBL, JHK

## Data and materials availability

All data underlying the analyses presented in this paper is available through the EBRAINS Knowledge Graph (https://doi.org/10.25493/KB7F-4VW). All the code used to process and analyze data is available in the DOPAMAP Github repository (https://github.com/ingvildeb/DOPAMAP). All the data from statistical analyses are available in the main text or the supplementary materials.

