## Supplementary file 1 for "The developing mouse dopaminergic system: Cortical-subcortical shift in D1/D2 receptor balance and increasing regional differentiation"

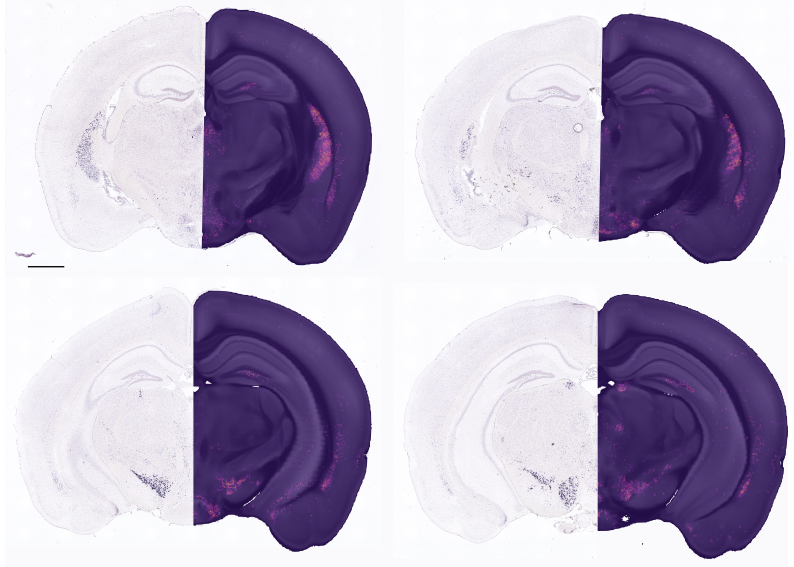

**Fig. S1. Comparison of posterior forebrain D2 data to Allen Institute data.** The figure shows sections from a case of Drd2 *in situ* hybridization data from the Allen Institute (case 357; left hemispheres) compared to our averaged D2 data^26^). The Allen Institute data was cut at an angle deviating ~11 degrees from the coronal plane; the average expression volume from our study was cut to match this angle in the EBRAINS interactive atlas viewer. Our data shows largely similar patterns to the Allen Institute data, with high expression in the caudate-putamen, substantia nigra, and ventral tegmental area, as well as clear staining in the endopiriform nucleus. Our data show somewhat higher expression in the mammillary nuclei of the hypothalamus; unfortunately, only one case was available from the Allen Institute, preventing further comparison. We observed clear D2 staining in this area in all cases covering this region (n = 5 across age groups).

|  |  | **D1:D2 Ratio** | | |
| --- | --- | --- | --- | --- |
|  |  | **Age** | **Sex** | **Age x Sex** |
| **Motor areas** | **F** | 18.040 | 4.995 | 4.998 |
|  | **p** | <.001 | .029 | .002 |
| **Somatosensory areas** | **F** | 10.035 | 7.993 | .899 |
|  | **p** | <.001 | .007 | .471 |
| **Gustatory and visceral areas** | **F** | 6.322 | 25.709 | 2.816 |
|  | **p** | <.001 | <.001 | .033 |
| **Anterior cingulate areas** | **F** | 12.836 | .091 | .553 |
|  | **p** | <.001 | .764 | .698 |
| **Prefrontal areas** | **F** | 8.587 | 22.097 | .408 |
|  | **p** | <.001 | <.001 | .802 |
| **Retrosplenial areas** | **F** | 23.498 | 23.385 | .498 |
|  | **p** | <.001 | <.001 | .686 |
| **Olfactory areas** | **F** | 9.407 | 18.553 | .425 |
|  | **p** | <.001 | <.001 | .790 |
| **Hippocampal region** | **F** | 1.810 | 9.548 | 3.150 |
|  | **p** | .140 | .003 | .021 |
| **Cortical subplate** | **F** | 3.093 | 15.041 | .445 |
|  | **p** | .023 | <.001 | .776 |
| **Striatum** | **F** | 6.299 | 21.968 | .160 |
|  | **p** | <.001 | <.001 | .958 |
| **Striatum-like amygdalar areas** | **F** | 8.560 | 73.804 | 2.476 |
|  | **p** | <.001 | <.001 | .055 |
| **Pallidum** | **F** | 2.312 | 4.560 | 2.000 |
|  | **p** | .069 | .037 | .107 |
| **Thalamus, sensory-motor related** | **F** | 3.391 | .087 | .533 |
|  | **p** | .016 | .769 | .712 |
| **Thalamus, polymodal association cortex related** | **F** | 11.170 | 3.496 | 2.000 |
|  | **p** | <.001 | .067 | .108 |
| **Hypothalamus, other** | **F** | 3.329 | 16.466 | 1.438 |
|  | **p** | .016 | <.001 | .233 |
| **Hypothalamus, medial zone** | **F** | 2.837 | 16.238 | 2.945 |
|  | **p** | .033 | <.001 | .029 |
| **Hypothalamus, lateral zone** | **F** | 1.770 | 16.410 | 1.097 |
|  | **p** | .148 | <.001 | .367 |

**Table S1. Results from statistical analysis of D1:D2 ratios across 17 major brain regions.** Two-way (age and sex factor) analyses of variances (ANOVA) with Tukey’s post hoc multiple comparisons was performed. The F and p values are shown.

|  |  | **Density** | | | | | | |
| --- | --- | --- | --- | --- | --- | --- | --- | --- |
|  |  | **Age** | **Sex** | **Receptor** | **Age x Sex** | **Age x Receptor** | **Sex x Receptor** | **Age x Sex x Receptor** |
| **Motor areas** | **F** | 3.940 | .411 | 121.320 | .396 | .560 | 1.059 | .570 |
|  | **p** | .005 | .523 | <.001 | .811 | .692 | .306 | .685 |
| **Somatosensory areas** | **F** | 7.021 | .191 | 105.509 | .181 | .421 | 2.922 | 1.485 |
|  | **p** | <.001 | .663 | <.001 | .947 | .793 | .091 | .213 |
| **Gustatory and visceral areas** | **F** | 4.618 | .030 | 109.975 | .336 | .443 | 2.580 | 1.165 |
|  | **p** | .002 | .863 | <.001 | .853 | .778 | .112 | .332 |
| **Anterior cingulate areas** | **F** | 2.911 | .415 | 95.399 | .304 | .283 | .605 | .509 |
|  | **p** | .026 | .521 | <.001 | .874 | .888 | .439 | .729 |
| **Prefrontal areas** | **F** | 1.770 | 1.972 | 116.830 | .520 | .347 | 4.145 | .264 |
|  | **p** | .142 | .164 | <.001 | .721 | .846 | .045 | .900 |
| **Retrosplenial areas** | **F** | 4.098 | .003 | 37.853 | .271 | .485 | 2.007 | 1.740 |
|  | **p** | .005 | .956 | <.001 | .846 | .694 | .162 | .169 |
| **Olfactory areas** | **F** | 2.503 | .289 | 66.809 | .481 | 1.283 | 5.331 | .797 |
|  | **p** | .048 | .592 | <.001 | .750 | .283 | .023 | .530 |
| **Hippocampal region** | **F** | 9.513 | .121 | 8.488 | .959 | 1.862 | 8.405 | 4.801 |
|  | **p** | <.001 | .729 | .005 | .434 | .124 | .005 | .001 |
| **Cortical subplate** | **F** | .674 | .045 | 8.130 | 1.232 | 1.069 | 5.389 | .365 |
|  | **p** | .612 | .832 | .005 | .303 | .376 | .022 | .833 |
| **Striatum** | **F** | 1.044 | 3.853 | 63.045 | .590 | 2.374 | 10.360 | .030 |
|  | **p** | .389 | .053 | <.001 | .671 | .058 | .002 | .998 |
| **Striatum-like amygdalar areas** | **F** | 1.240 | 5.891 | 1.675 | 1.197 | 2.302 | 30.184 | .351 |
|  | **p** | .300 | .017 | .199 | .318 | .065 | <.001 | .842 |
| **Pallidum** | **F** | 5.654 | 1.099 | 77.122 | 1.113 | 2.407 | 2.976 | .423 |
|  | **p** | <.001 | .297 | <.001 | .355 | .055 | .088 | .791 |
| **Thalamus, sensory-motor related** | **F** | 9.745 | 2.481 | .615 | 1.144 | 4.206 | 3.689 | 3.944 |
|  | **p** | <.001 | .119 | .435 | .342 | .004 | .058 | .006 |
| **Thalamus, polymodal association cortex related** | **F** | 9.030 | .280 | 19.263 | .388 | 3.284 | 1.011 | 1.182 |
|  | **p** | <.001 | .598 | <.001 | .817 | .015 | .318 | .325 |
| **Hypothalamus, other** | **F** | 6.351 | .012 | 158.704 | .887 | 3.975 | 11.301 | .434 |
|  | **p** | <.001 | .913 | <.001 | .475 | .005 | .001 | .784 |
| **Hypothalamus, medial zone** | **F** | 15.643 | 2.225 | 99.384 | .666 | 8.237 | 3.572 | 1.499 |
|  | **p** | <.001 | .140 | <.001 | .618 | <.001 | .062 | .210 |
| **Hypothalamus, lateral zone** | **F** | 2.998 | 2.881 | .653 | .610 | .708 | 6.401 | .726 |
|  | **p** | .023 | .093 | .421 | .656 | .589 | .013 | .577 |

**Table S2. Results from statistical analysis of D1 and D2 densities across 17 major brain regions.** Three-way (receptor, age and sex factor) analyses of variances (ANOVA) with Tukey’s post hoc multiple comparisons was performed. The F and p values are shown.

|  |  | **MO** | **SS** | **GU/V** | **ACC** | **PFA** | **RSP** | **Olf** | **HR** | **CTX-s** | **STR** | **sAMY** | **PAL** | **Th-S** | **Th-P** | **Hy-o** | **Hy-m** | **Hy-l** |
| --- | --- | --- | --- | --- | --- | --- | --- | --- | --- | --- | --- | --- | --- | --- | --- | --- | --- | --- |
| **MO** | **r** | 1 | **.908^**^** | **.914^**^** | **.923^**^** | **.974^**^** | **.807^**^** | **.830^**^** | **.695^**^** | **.781^**^** | **.811^**^** | **.777^**^** | **.649^**^** | 0,335 | **.669^**^** | **.539^*^** | **.586^*^** | 0,330 |
|  | **p** |  | 0,000 | 0,000 | 0,000 | 0,000 | 0,000 | 0,000 | 0,003 | 0,000 | 0,000 | 0,001 | 0,007 | 0,241 | 0,006 | 0,038 | 0,022 | 0,230 |
|  | **n** | 16 | 16 | 16 | 16 | 16 | 16 | 16 | 16 | 16 | 16 | 15 | 16 | 14 | 15 | 15 | 15 | 15 |
| **SS** | **r** | **.908^**^** | 1 | **.952^**^** | **.960^**^** | **.909^**^** | **.950^**^** | **.915^**^** | **.751^**^** | **.851^**^** | **.880^**^** | **.822^**^** | **.751^**^** | 0,417 | **.617^*^** | **.654^**^** | **.674^**^** | 0,440 |
|  | **p** | 0,000 |  | 0,000 | 0,000 | 0,000 | 0,000 | 0,000 | 0,001 | 0,000 | 0,000 | 0,000 | 0,001 | 0,138 | 0,014 | 0,008 | 0,006 | 0,101 |
|  | **n** | 16 | 16 | 16 | 16 | 16 | 16 | 16 | 16 | 16 | 16 | 15 | 16 | 14 | 15 | 15 | 15 | 15 |
| **GU/V** | **r** | **.914^**^** | **.952^**^** | 1 | **.925^**^** | **.920^**^** | **.905^**^** | **.893^**^** | **.818^**^** | **.862^**^** | **.843^**^** | **.827^**^** | **.784^**^** | 0,421 | **.724^**^** | **.687^**^** | **.715^**^** | 0,513 |
|  | **p** | 0,000 | 0,000 |  | 0,000 | 0,000 | 0,000 | 0,000 | 0,000 | 0,000 | 0,000 | 0,000 | 0,000 | 0,134 | 0,002 | 0,005 | 0,003 | 0,051 |
|  | **n** | 16 | 16 | 16 | 16 | 16 | 16 | 16 | 16 | 16 | 16 | 15 | 16 | 14 | 15 | 15 | 15 | 15 |
| **ACC** | **r** | **.923^**^** | **.960^**^** | **.925^**^** | 1 | **.942^**^** | **.867^**^** | **.907^**^** | **.767^**^** | **.856^**^** | **.953^**^** | **.804^**^** | **.796^**^** | 0,496 | **.641^*^** | **.654^**^** | **.685^**^** | 0,439 |
|  | **p** | 0,000 | 0,000 | 0,000 |  | 0,000 | 0,000 | 0,000 | 0,001 | 0,000 | 0,000 | 0,000 | 0,000 | 0,071 | 0,010 | 0,008 | 0,005 | 0,102 |
|  | **n** | 16 | 16 | 16 | 16 | 16 | 16 | 16 | 16 | 16 | 16 | 15 | 16 | 14 | 15 | 15 | 15 | 15 |
| **PFA** | **r** | **.974^**^** | **.909^**^** | **.920^**^** | **.942^**^** | 1 | **.814^**^** | **.871^**^** | **.762^**^** | **.816^**^** | **.855^**^** | **.829^**^** | **.718^**^** | 0,389 | **.723^**^** | **.625^*^** | **.674^**^** | 0,380 |
|  | **p** | 0,000 | 0,000 | 0,000 | 0,000 |  | 0,000 | 0,000 | 0,001 | 0,000 | 0,000 | 0,000 | 0,002 | 0,169 | 0,002 | 0,013 | 0,006 | 0,162 |
|  | **n** | 16 | 16 | 16 | 16 | 16 | 16 | 16 | 16 | 16 | 16 | 15 | 16 | 14 | 15 | 15 | 15 | 15 |
| **RSP** | **r** | **.807^**^** | **.950^**^** | **.905^**^** | **.867^**^** | **.814^**^** | 1 | **.897^**^** | **.721^**^** | **.829^**^** | **.759^**^** | **.828^**^** | **.678^**^** | 0,342 | **.529^*^** | **.653^**^** | **.691^**^** | 0,368 |
|  | **p** | 0,000 | 0,000 | 0,000 | 0,000 | 0,000 |  | 0,000 | 0,002 | 0,000 | 0,001 | 0,000 | 0,004 | 0,231 | 0,042 | 0,008 | 0,004 | 0,177 |
|  | **n** | 16 | 16 | 16 | 16 | 16 | 16 | 16 | 16 | 16 | 16 | 15 | 16 | 14 | 15 | 15 | 15 | 15 |
| **Olf** | **r** | **.830^**^** | **.915^**^** | **.893^**^** | **.907^**^** | **.871^**^** | **.897^**^** | 1 | **.869^**^** | **.947^**^** | **.871^**^** | **.903^**^** | **.873^**^** | **.565^*^** | **.719^**^** | **.788^**^** | **.832^**^** | 0,443 |
|  | **p** | 0,000 | 0,000 | 0,000 | 0,000 | 0,000 | 0,000 |  | 0,000 | 0,000 | 0,000 | 0,000 | 0,000 | 0,035 | 0,003 | 0,000 | 0,000 | 0,098 |
|  | **n** | 16 | 16 | 16 | 16 | 16 | 16 | 16 | 16 | 16 | 16 | 15 | 16 | 14 | 15 | 15 | 15 | 15 |
| **HR** | **r** | **.695^**^** | **.751^**^** | **.818^**^** | **.767^**^** | **.762^**^** | **.721^**^** | **.869^**^** | 1 | **.859^**^** | **.750^**^** | **.860^**^** | **.879^**^** | **.539^*^** | **.844^**^** | **.767^**^** | **.849^**^** | 0,482 |
|  | **p** | 0,003 | 0,001 | 0,000 | 0,001 | 0,001 | 0,002 | 0,000 |  | 0,000 | 0,001 | 0,000 | 0,000 | 0,047 | 0,000 | 0,001 | 0,000 | 0,069 |
|  | **n** | 16 | 16 | 16 | 16 | 16 | 16 | 16 | 16 | 16 | 16 | 15 | 16 | 14 | 15 | 15 | 15 | 15 |
| **CTX-s** | **r** | **.781^**^** | **.851^**^** | **.862^**^** | **.856^**^** | **.816^**^** | **.829^**^** | **.947^**^** | **.859^**^** | 1 | **.824^**^** | **.896^**^** | **.840^**^** | **.731^**^** | **.805^**^** | **.700^**^** | **.755^**^** | 0,369 |
|  | **p** | 0,000 | 0,000 | 0,000 | 0,000 | 0,000 | 0,000 | 0,000 | 0,000 |  | 0,000 | 0,000 | 0,000 | 0,003 | 0,000 | 0,004 | 0,001 | 0,176 |
|  | **n** | 16 | 16 | 16 | 16 | 16 | 16 | 16 | 16 | 16 | 16 | 15 | 16 | 14 | 15 | 15 | 15 | 15 |
| **STR** | **r** | **.811^**^** | **.880^**^** | **.843^**^** | **.953^**^** | **.855^**^** | **.759^**^** | **.871^**^** | **.750^**^** | **.824^**^** | 1 | **.721^**^** | **.859^**^** | **.571^*^** | **.585^*^** | **.677^**^** | **.668^**^** | **.532^*^** |
|  | **p** | 0,000 | 0,000 | 0,000 | 0,000 | 0,000 | 0,001 | 0,000 | 0,001 | 0,000 |  | 0,002 | 0,000 | 0,033 | 0,022 | 0,006 | 0,007 | 0,041 |
|  | **n** | 16 | 16 | 16 | 16 | 16 | 16 | 16 | 16 | 16 | 16 | 15 | 16 | 14 | 15 | 15 | 15 | 15 |
| **sAMY** | **r** | **.777^**^** | **.822^**^** | **.827^**^** | **.804^**^** | **.829^**^** | **.828^**^** | **.903^**^** | **.860^**^** | **.896^**^** | **.721^**^** | 1 | **.726^**^** | **.554^*^** | **.793^**^** | **.606^*^** | **.761^**^** | 0,222 |
|  | **p** | 0,001 | 0,000 | 0,000 | 0,000 | 0,000 | 0,000 | 0,000 | 0,000 | 0,000 | 0,002 |  | 0,002 | 0,040 | 0,000 | 0,017 | 0,001 | 0,426 |
|  | **n** | 15 | 15 | 15 | 15 | 15 | 15 | 15 | 15 | 15 | 15 | 15 | 15 | 14 | 15 | 15 | 15 | 15 |
| **PAL** | **r** | **.649^**^** | **.751^**^** | **.784^**^** | **.796^**^** | **.718^**^** | **.678^**^** | **.873^**^** | **.879^**^** | **.840^**^** | **.859^**^** | **.726^**^** | 1 | **.587^*^** | **.710^**^** | **.879^**^** | **.887^**^** | **.702^**^** |
|  | **p** | 0,007 | 0,001 | 0,000 | 0,000 | 0,002 | 0,004 | 0,000 | 0,000 | 0,000 | 0,000 | 0,002 |  | 0,027 | 0,003 | 0,000 | 0,000 | 0,004 |
|  | **n** | 16 | 16 | 16 | 16 | 16 | 16 | 16 | 16 | 16 | 16 | 15 | 16 | 14 | 15 | 15 | 15 | 15 |
| **Th-S** | **r** | 0,335 | 0,417 | 0,421 | 0,496 | 0,389 | 0,342 | **.565^*^** | **.539^*^** | **.731^**^** | **.571^*^** | **.554^*^** | **.587^*^** | 1 | **.805^**^** | 0,373 | 0,370 | 0,163 |
|  | **p** | 0,241 | 0,138 | 0,134 | 0,071 | 0,169 | 0,231 | 0,035 | 0,047 | 0,003 | 0,033 | 0,040 | 0,027 |  | 0,001 | 0,189 | 0,193 | 0,578 |
|  | **n** | 14 | 14 | 14 | 14 | 14 | 14 | 14 | 14 | 14 | 14 | 14 | 14 | 14 | 14 | 14 | 14 | 14 |
| **Th-P** | **r** | **.669^**^** | **.617^*^** | **.724^**^** | **.641^*^** | **.723^**^** | **.529^*^** | **.719^**^** | **.844^**^** | **.805^**^** | **.585^*^** | **.793^**^** | **.710^**^** | **.805^**^** | 1 | **.541^*^** | **.636^*^** | 0,290 |
|  | **p** | 0,006 | 0,014 | 0,002 | 0,010 | 0,002 | 0,042 | 0,003 | 0,000 | 0,000 | 0,022 | 0,000 | 0,003 | 0,001 |  | 0,037 | 0,011 | 0,294 |
|  | **n** | 15 | 15 | 15 | 15 | 15 | 15 | 15 | 15 | 15 | 15 | 15 | 15 | 14 | 15 | 15 | 15 | 15 |
| **Hy-o** | **r** | **.539^*^** | **.654^**^** | **.687^**^** | **.654^**^** | **.625^*^** | **.653^**^** | **.788^**^** | **.767^**^** | **.700^**^** | **.677^**^** | **.606^*^** | **.879^**^** | 0,373 | **.541^*^** | 1 | **.940^**^** | **.786^**^** |
|  | **p** | 0,038 | 0,008 | 0,005 | 0,008 | 0,013 | 0,008 | 0,000 | 0,001 | 0,004 | 0,006 | 0,017 | 0,000 | 0,189 | 0,037 |  | 0,000 | 0,001 |
|  | **n** | 15 | 15 | 15 | 15 | 15 | 15 | 15 | 15 | 15 | 15 | 15 | 15 | 14 | 15 | 15 | 15 | 15 |
| **Hy-m** | **r** | **.586^*^** | **.674^**^** | **.715^**^** | **.685^**^** | **.674^**^** | **.691^**^** | **.832^**^** | **.849^**^** | **.755^**^** | **.668^**^** | **.761^**^** | **.887^**^** | 0,370 | **.636^*^** | **.940^**^** | 1 | **.626^*^** |
|  | **p** | 0,022 | 0,006 | 0,003 | 0,005 | 0,006 | 0,004 | 0,000 | 0,000 | 0,001 | 0,007 | 0,001 | 0,000 | 0,193 | 0,011 | 0,000 |  | 0,013 |
|  | **n** | 15 | 15 | 15 | 15 | 15 | 15 | 15 | 15 | 15 | 15 | 15 | 15 | 14 | 15 | 15 | 15 | 15 |
| **Hy-l** | **r** | 0,330 | 0,440 | 0,513 | 0,439 | 0,380 | 0,368 | 0,443 | 0,482 | 0,369 | **.532^*^** | 0,222 | **.702^**^** | 0,163 | 0,290 | **.786^**^** | **.626^*^** | 1 |
|  | **p** | 0,230 | 0,101 | 0,051 | 0,102 | 0,162 | 0,177 | 0,098 | 0,069 | 0,176 | 0,041 | 0,426 | 0,004 | 0,578 | 0,294 | 0,001 | 0,013 |  |
|  | **n** | 15 | 15 | 15 | 15 | 15 | 15 | 15 | 15 | 15 | 15 | 15 | 15 | 14 | 15 | 15 | 15 | 15 |

**Table S3. Results from correlation analysis of cell densities across regions for the D1 P17 group.** Two-tailed Pearson’s correlation (r) tests were used; pearson’s correlation coefficient (r), p value (p) and the number of subjects (n) is shown for each region pair.

|  |  | **MO** | **SS** | **GU/V** | **ACC** | **PFA** | **RSP** | **Olf** | **HR** | **CTX-s** | **STR** | **sAMY** | **PAL** | **Th-S** | **Th-P** | **Hy-o** | **Hy-m** | **Hy-l** |
| --- | --- | --- | --- | --- | --- | --- | --- | --- | --- | --- | --- | --- | --- | --- | --- | --- | --- | --- |
| **MO** | **r** | 1 | **.871**** | **.799**** | **.836**** | **.913**** | **.809**** | **.773**** | **.581*** | **.612*** | **.858**** | **.602*** | **.602*** | 0,465 | **.721**** | **0,37** | **0,27** | 0,454 |
|  | **p** |  | 0,000 | 0,001 | 0,000 | 0,000 | 0,001 | 0,001 | 0,029 | 0,020 | 0,000 | 0,023 | 0,023 | 0,128 | 0,005 | 0,189 | 0,380 | 0,103 |
|  | **n** | 14 | 14 | 14 | 14 | 14 | 12 | 14 | 14 | 14 | 14 | 14 | 14 | 12 | 13 | 14 | 13 | 14 |
| **SS** | **r** | **.871**** | 1 | **.815**** | **.824**** | **.855**** | **.930**** | **.701**** | **.687**** | **.560*** | **.887**** | **.668**** | **.648*** | .689* | **.874**** | **0,33** | **0,21** | 0,512 |
|  | **p** | 0,000 |  | 0,000 | 0,000 | 0,000 | 0,000 | 0,005 | 0,007 | 0,037 | 0,000 | 0,009 | 0,012 | 0,013 | 0,000 | 0,251 | 0,486 | 0,061 |
|  | **n** | 14 | 14 | 14 | 14 | 14 | 12 | 14 | 14 | 14 | 14 | 14 | 14 | 12 | 13 | 14 | 13 | 14 |
| **GU/V** | **r** | **.799**** | **.815**** | 1 | **.898**** | **.914**** | **.688*** | **.789**** | **.741**** | **.765**** | **.872**** | **.677**** | **.707**** | .741** | **.856**** | **0,47** | **0,31** | 0,295 |
|  | **p** | 0,001 | 0,000 |  | 0,000 | 0,000 | 0,013 | 0,001 | 0,002 | 0,001 | 0,000 | 0,008 | 0,005 | 0,006 | 0,000 | 0,091 | 0,308 | 0,305 |
|  | **n** | 14 | 14 | 14 | 14 | 14 | 12 | 14 | 14 | 14 | 14 | 14 | 14 | 12 | 13 | 14 | 13 | 14 |
| **ACC** | **r** | **.836**** | **.824**** | **.898**** | 1 | **.934**** | **.762**** | **.649*** | **.660*** | **.588*** | **.902**** | **.609*** | **.637*** | .773** | **.885**** | **0,32** | **0,36** | .599* |
|  | **p** | 0,000 | 0,000 | 0,000 |  | 0,000 | 0,004 | 0,012 | 0,010 | 0,027 | 0,000 | 0,021 | 0,014 | 0,003 | 0,000 | 0,262 | 0,228 | 0,024 |
|  | **n** | 14 | 14 | 14 | 14 | 14 | 12 | 14 | 14 | 14 | 14 | 14 | 14 | 12 | 13 | 14 | 13 | 14 |
| **PFA** | **r** | **.913**** | **.855**** | **.914**** | **.934**** | 1 | **.842**** | **.824**** | **.698**** | **.740**** | **.913**** | **.686**** | **.665**** | .629* | **.792**** | **0,47** | **0,39** | 0,454 |
|  | **p** | 0,000 | 0,000 | 0,000 | 0,000 |  | 0,001 | 0,000 | 0,005 | 0,003 | 0,000 | 0,007 | 0,010 | 0,028 | 0,001 | 0,088 | 0,183 | 0,103 |
|  | **n** | 14 | 14 | 14 | 14 | 14 | 12 | 14 | 14 | 14 | 14 | 14 | 14 | 12 | 13 | 14 | 13 | 14 |
| **RSP** | **r** | **.809**** | **.930**** | **.688*** | **.762**** | **.842**** | 1 | **.842**** | **.804**** | **.640*** | **.868**** | **.821**** | **.742**** | .605* | **.777**** | **.767**** | **.743**** | .651* |
|  | **p** | 0,001 | 0,000 | 0,013 | 0,004 | 0,001 |  | 0,001 | 0,002 | 0,025 | 0,000 | 0,001 | 0,006 | 0,037 | 0,003 | 0,004 | 0,006 | 0,022 |
|  | **n** | 12 | 12 | 12 | 12 | 12 | 12 | 12 | 12 | 12 | 12 | 12 | 12 | 12 | 12 | 12 | 12 | 12 |
| **Olf** | **r** | **.773**** | **.701**** | **.789**** | **.649*** | **.824**** | **.842**** | 1 | **.852**** | **.836**** | **.760**** | **.808**** | **.809**** | **0,54** | **.683*** | **.800**** | **.636*** | 0,006 |
|  | **p** | 0,001 | 0,005 | 0,001 | 0,012 | 0,000 | 0,001 |  | 0,000 | 0,000 | 0,002 | 0,000 | 0,000 | 0,070 | 0,010 | 0,001 | 0,019 | 0,985 |
|  | **n** | 14 | 14 | 14 | 14 | 14 | 12 | 14 | 14 | 14 | 14 | 14 | 14 | 12 | 13 | 14 | 13 | 14 |
| **HR** | **r** | **.581*** | **.687**** | **.741**** | **.660*** | **.698**** | **.804**** | **.852**** | 1 | **.717**** | **.751**** | **.901**** | **.950**** | **.832**** | **.813**** | **.785**** | **.698**** | 0,061 |
|  | **p** | 0,029 | 0,007 | 0,002 | 0,010 | 0,005 | 0,002 | 0,000 |  | 0,004 | 0,002 | 0,000 | 0,000 | 0,001 | 0,001 | 0,001 | 0,008 | 0,835 |
|  | **n** | 14 | 14 | 14 | 14 | 14 | 12 | 14 | 14 | 14 | 14 | 14 | 14 | 12 | 13 | 14 | 13 | 14 |
| **CTX-s** | **r** | **.612*** | **.560*** | **.765**** | **.588*** | **.740**** | **.640*** | **.836**** | **.717**** | 1 | **.661*** | **.773**** | **.762**** | **0,47** | **0,51** | **.809**** | **.614*** | -0,070 |
|  | **p** | 0,020 | 0,037 | 0,001 | 0,027 | 0,003 | 0,025 | 0,000 | 0,004 |  | 0,010 | 0,001 | 0,002 | 0,120 | 0,073 | 0,000 | 0,026 | 0,813 |
|  | **n** | 14 | 14 | 14 | 14 | 14 | 12 | 14 | 14 | 14 | 14 | 14 | 14 | 12 | 13 | 14 | 13 | 14 |
| **STR** | **r** | **.858**** | **.887**** | **.872**** | **.902**** | **.913**** | **.868**** | **.760**** | **.751**** | **.661*** | 1 | **.758**** | **.726**** | **.771**** | **.910**** | **0,44** | **0,44** | **0,52** |
|  | **p** | 0,000 | 0,000 | 0,000 | 0,000 | 0,000 | 0,000 | 0,002 | 0,002 | 0,010 |  | 0,002 | 0,003 | 0,003 | 0,000 | 0,117 | 0,128 | 0,057 |
|  | **n** | 14 | 14 | 14 | 14 | 14 | 12 | 14 | 14 | 14 | 14 | 14 | 14 | 12 | 13 | 14 | 13 | 14 |
| **sAMY** | **r** | **.602*** | **.668**** | **.677**** | **.609*** | **.686**** | **.821**** | **.808**** | **.901**** | **.773**** | **.758**** | 1 | **.919**** | **.725**** | **.694**** | **.777**** | **.690**** | 0,135 |
|  | **p** | 0,023 | 0,009 | 0,008 | 0,021 | 0,007 | 0,001 | 0,000 | 0,000 | 0,001 | 0,002 |  | 0,000 | 0,008 | 0,009 | 0,001 | 0,009 | 0,646 |
|  | **n** | 14 | 14 | 14 | 14 | 14 | 12 | 14 | 14 | 14 | 14 | 14 | 14 | 12 | 13 | 14 | 13 | 14 |
| **PAL** | **r** | **.602*** | **.648*** | **.707**** | **.637*** | **.665**** | **.742**** | **.809**** | **.950**** | **.762**** | **.726**** | **.919**** | 1 | **.762**** | **.711**** | **.829**** | **.731**** | **0,05** |
|  | **p** | 0,023 | 0,012 | 0,005 | 0,014 | 0,010 | 0,006 | 0,000 | 0,000 | 0,002 | 0,003 | 0,000 |  | 0,004 | 0,006 | 0,000 | 0,004 | 0,865 |
|  | **n** | 14 | 14 | 14 | 14 | 14 | 12 | 14 | 14 | 14 | 14 | 14 | 14 | 12 | 13 | 14 | 13 | 14 |
| **Th-S** | **r** | 0,465 | .689* | .741** | .773** | .629* | .605* | **0,54** | **.832**** | **0,47** | **.771**** | **.725**** | **.762**** | 1 | **.917**** | 0,534 | .666* | .738** |
|  | **p** | 0,128 | 0,013 | 0,006 | 0,003 | 0,028 | 0,037 | 0,070 | 0,001 | 0,120 | 0,003 | 0,008 | 0,004 |  | 0,000 | 0,074 | 0,018 | 0,006 |
|  | **n** | 12 | 12 | 12 | 12 | 12 | 12 | 12 | 12 | 12 | 12 | 12 | 12 | 12 | 12 | 12 | 12 | 12 |
| **Th-P** | **r** | **.721**** | **.874**** | **.856**** | **.885**** | **.792**** | **.777**** | **.683*** | **.813**** | **0,51** | **.910**** | **.694**** | **.711**** | **.917**** | 1 | **0,31** | **0,35** | .667* |
|  | **p** | 0,005 | 0,000 | 0,000 | 0,000 | 0,001 | 0,003 | 0,010 | 0,001 | 0,073 | 0,000 | 0,009 | 0,006 | 0,000 |  | 0,302 | 0,242 | 0,013 |
|  | **n** | 13 | 13 | 13 | 13 | 13 | 12 | 13 | 13 | 13 | 13 | 13 | 13 | 12 | 13 | 13 | 13 | 13 |
| **Hy-o** | **r** | **0,37** | **0,33** | **0,47** | **0,32** | **0,47** | **.767**** | **.800**** | **.785**** | **.809**** | **0,44** | **.777**** | **.829**** | 0,534 | **0,31** | 1 | **.948**** | **-0,28** |
|  | **p** | 0,189 | 0,251 | 0,091 | 0,262 | 0,088 | 0,004 | 0,001 | 0,001 | 0,000 | 0,117 | 0,001 | 0,000 | 0,074 | 0,302 |  | 0,000 | 0,339 |
|  | **n** | 14 | 14 | 14 | 14 | 14 | 12 | 14 | 14 | 14 | 14 | 14 | 14 | 12 | 13 | 14 | 13 | 14 |
| **Hy-m** | **r** | **0,27** | **0,21** | **0,31** | **0,36** | **0,39** | **.743**** | **.636*** | **.698**** | **.614*** | **0,44** | **.690**** | **.731**** | .666* | **0,35** | **.948**** | 1 | **0,08** |
|  | **p** | 0,380 | 0,486 | 0,308 | 0,228 | 0,183 | 0,006 | 0,019 | 0,008 | 0,026 | 0,128 | 0,009 | 0,004 | 0,018 | 0,242 | 0,000 |  | 0,800 |
|  | **n** | 13 | 13 | 13 | 13 | 13 | 12 | 13 | 13 | 13 | 13 | 13 | 13 | 12 | 13 | 13 | 13 | 13 |
| **Hy-l** | **r** | 0,454 | 0,512 | 0,295 | .599* | 0,454 | .651* | 0,006 | 0,061 | -0,070 | **0,52** | 0,135 | **0,05** | .738** | .667* | **-0,28** | **0,08** | 1 |
|  | **p** | 0,103 | 0,061 | 0,305 | 0,024 | 0,103 | 0,022 | 0,985 | 0,835 | 0,813 | 0,057 | 0,646 | 0,865 | 0,006 | 0,013 | 0,339 | 0,800 |  |
|  | **n** | 14 | 14 | 14 | 14 | 14 | 12 | 14 | 14 | 14 | 14 | 14 | 14 | 12 | 13 | 14 | 13 | 14 |

**Table S4. Results from correlation analysis of cell densities across regions for the D1 P25 group.** Two-tailed Pearson’s correlation (r) tests were used; pearson’s correlation coefficient (r), p value (p) and the number of subjects (n) is shown for each region pair.

|  |  | **MO** | **SS** | **GU/V** | **ACC** | **PFA** | **RSP** | **Olf** | **HR** | **CTX-s** | **STR** | **sAMY** | **PAL** | **Th-S** | **Th-P** | **Hy-o** | **Hy-m** | **Hy-l** |
| --- | --- | --- | --- | --- | --- | --- | --- | --- | --- | --- | --- | --- | --- | --- | --- | --- | --- | --- |
| **MO** | **r** | 1 | **.607*** | **.764**** | **.828**** | **.898**** | **0,55** | **.682**** | **0,23** | **.717**** | **.800**** | **0,36** | **0,26** | 0,025 | **0,03** | **0,35** | **0,4** | 0,288 |
|  | **p** |  | 0,021 | 0,001 | 0,000 | 0,000 | 0,123 | 0,007 | 0,420 | 0,004 | 0,001 | 0,225 | 0,362 | 0,938 | 0,912 | 0,216 | 0,181 | 0,317 |
|  | **n** | 14 | 14 | 14 | 14 | 14 | 9 | 14 | 14 | 14 | 14 | 13 | 14 | 12 | 14 | 14 | 13 | 14 |
| **SS** | **r** | **.607*** | 1 | **.779**** | **.761**** | **.829**** | **.669*** | **.847**** | **.783**** | **.790**** | **.818**** | **.811**** | **.747**** | 0,354 | **0,12** | **.796**** | **.834**** | -0,028 |
|  | **p** | 0,021 |  | 0,001 | 0,002 | 0,000 | 0,049 | 0,000 | 0,001 | 0,001 | 0,000 | 0,001 | 0,002 | 0,259 | 0,679 | 0,001 | 0,000 | 0,923 |
|  | **n** | 14 | 14 | 14 | 14 | 14 | 9 | 14 | 14 | 14 | 14 | 13 | 14 | 12 | 14 | 14 | 13 | 14 |
| **GU/V** | **r** | **.764**** | **.779**** | 1 | **.712**** | **.832**** | **0,35** | **.767**** | **.562*** | **.686**** | **.785**** | **.794**** | **.609*** | 0,143 | **0,07** | **.630*** | **.647*** | 0,276 |
|  | **p** | 0,001 | 0,001 |  | 0,004 | 0,000 | 0,359 | 0,001 | 0,037 | 0,007 | 0,001 | 0,001 | 0,021 | 0,657 | 0,801 | 0,016 | 0,017 | 0,340 |
|  | **n** | 14 | 14 | 14 | 14 | 14 | 9 | 14 | 14 | 14 | 14 | 13 | 14 | 12 | 14 | 14 | 13 | 14 |
| **ACC** | **r** | **.828**** | **.761**** | **.712**** | 1 | **.921**** | **0,62** | **.751**** | **.565*** | **.839**** | **.848**** | **0,53** | **0,51** | 0,419 | **0,16** | **0,49** | **.559*** | 0,012 |
|  | **p** | 0,000 | 0,002 | 0,004 |  | 0,000 | 0,075 | 0,002 | 0,035 | 0,000 | 0,000 | 0,062 | 0,061 | 0,175 | 0,593 | 0,072 | 0,047 | 0,967 |
|  | **n** | 14 | 14 | 14 | 14 | 14 | 9 | 14 | 14 | 14 | 14 | 13 | 14 | 12 | 14 | 14 | 13 | 14 |
| **PFA** | **r** | **.898**** | **.829**** | **.832**** | **.921**** | 1 | **.770*** | **.875**** | **.547*** | **.858**** | **.883**** | **.640*** | **.560*** | 0,355 | **0,1** | **.571*** | **.670*** | 0,090 |
|  | **p** | 0,000 | 0,000 | 0,000 | 0,000 |  | 0,015 | 0,000 | 0,043 | 0,000 | 0,000 | 0,019 | 0,037 | 0,257 | 0,736 | 0,033 | 0,012 | 0,761 |
|  | **n** | 14 | 14 | 14 | 14 | 14 | 9 | 14 | 14 | 14 | 14 | 13 | 14 | 12 | 14 | 14 | 13 | 14 |
| **RSP** | **r** | **0,55** | **.669*** | **0,35** | **0,62** | **.770*** | 1 | **0,66** | **0,64** | **0,56** | **0,44** | **0,15** | **0,64** | 0,275 | **0,22** | **0,39** | **0,57** | 0,164 |
|  | **p** | 0,123 | 0,049 | 0,359 | 0,075 | 0,015 |  | 0,051 | 0,065 | 0,114 | 0,239 | 0,731 | 0,065 | 0,509 | 0,563 | 0,296 | 0,138 | 0,674 |
|  | **n** | 9 | 9 | 9 | 9 | 9 | 9 | 9 | 9 | 9 | 9 | 8 | 9 | 8 | 9 | 9 | 8 | 9 |
| **Olf** | **r** | **.682**** | **.847**** | **.767**** | **.751**** | **.875**** | **0,66** | 1 | **.700**** | **.904**** | **.869**** | **.770**** | **.786**** | **0,52** | **0,08** | **.794**** | **.865**** | -0,199 |
|  | **p** | 0,007 | 0,000 | 0,001 | 0,002 | 0,000 | 0,051 |  | 0,005 | 0,000 | 0,000 | 0,002 | 0,001 | 0,084 | 0,794 | 0,001 | 0,000 | 0,495 |
|  | **n** | 14 | 14 | 14 | 14 | 14 | 9 | 14 | 14 | 14 | 14 | 13 | 14 | 12 | 14 | 14 | 13 | 14 |
| **HR** | **r** | **0,23** | **.783**** | **.562*** | **.565*** | **.547*** | **0,64** | **.700**** | 1 | **.686**** | **0,53** | **.745**** | **.841**** | **.661*** | **0,46** | **.767**** | **.750**** | -0,391 |
|  | **p** | 0,420 | 0,001 | 0,037 | 0,035 | 0,043 | 0,065 | 0,005 |  | 0,007 | 0,051 | 0,003 | 0,000 | 0,019 | 0,100 | 0,001 | 0,003 | 0,167 |
|  | **n** | 14 | 14 | 14 | 14 | 14 | 9 | 14 | 14 | 14 | 14 | 13 | 14 | 12 | 14 | 14 | 13 | 14 |
| **CTX-s** | **r** | **.717**** | **.790**** | **.686**** | **.839**** | **.858**** | **0,56** | **.904**** | **.686**** | 1 | **.847**** | **.637*** | **.710**** | **0,56** | **0,02** | **.734**** | **.779**** | -0,214 |
|  | **p** | 0,004 | 0,001 | 0,007 | 0,000 | 0,000 | 0,114 | 0,000 | 0,007 |  | 0,000 | 0,019 | 0,004 | 0,061 | 0,944 | 0,003 | 0,002 | 0,461 |
|  | **n** | 14 | 14 | 14 | 14 | 14 | 9 | 14 | 14 | 14 | 14 | 13 | 14 | 12 | 14 | 14 | 13 | 14 |
| **STR** | **r** | **.800**** | **.818**** | **.785**** | **.848**** | **.883**** | **0,44** | **.869**** | **0,53** | **.847**** | 1 | **.661*** | **.677**** | **0,34** | **-0,05** | **.663**** | **.647*** | **-0,03** |
|  | **p** | 0,001 | 0,000 | 0,001 | 0,000 | 0,000 | 0,239 | 0,000 | 0,051 | 0,000 |  | 0,014 | 0,008 | 0,273 | 0,865 | 0,010 | 0,017 | 0,922 |
|  | **n** | 14 | 14 | 14 | 14 | 14 | 9 | 14 | 14 | 14 | 14 | 13 | 14 | 12 | 14 | 14 | 13 | 14 |
| **sAMY** | **r** | **0,36** | **.811**** | **.794**** | **0,53** | **.640*** | **0,15** | **.770**** | **.745**** | **.637*** | **.661*** | 1 | **.878**** | **0,5** | **-0,07** | **.826**** | **.841**** | -0,122 |
|  | **p** | 0,225 | 0,001 | 0,001 | 0,062 | 0,019 | 0,731 | 0,002 | 0,003 | 0,019 | 0,014 |  | 0,000 | 0,097 | 0,823 | 0,001 | 0,000 | 0,691 |
|  | **n** | 13 | 13 | 13 | 13 | 13 | 8 | 13 | 13 | 13 | 13 | 13 | 13 | 12 | 13 | 13 | 13 | 13 |
| **PAL** | **r** | **0,26** | **.747**** | **.609*** | **0,51** | **.560*** | **0,64** | **.786**** | **.841**** | **.710**** | **.677**** | **.878**** | 1 | **.787**** | **0,05** | **.789**** | **.760**** | **-0,4** |
|  | **p** | 0,362 | 0,002 | 0,021 | 0,061 | 0,037 | 0,065 | 0,001 | 0,000 | 0,004 | 0,008 | 0,000 |  | 0,002 | 0,853 | 0,001 | 0,003 | 0,159 |
|  | **n** | 14 | 14 | 14 | 14 | 14 | 9 | 14 | 14 | 14 | 14 | 13 | 14 | 12 | 14 | 14 | 13 | 14 |
| **Th-S** | **r** | 0,025 | 0,354 | 0,143 | 0,419 | 0,355 | 0,275 | **0,52** | **.661*** | **0,56** | **0,34** | **0,5** | **.787**** | 1 | **0,29** | 0,493 | 0,509 | -.597* |
|  | **p** | 0,938 | 0,259 | 0,657 | 0,175 | 0,257 | 0,509 | 0,084 | 0,019 | 0,061 | 0,273 | 0,097 | 0,002 |  | 0,369 | 0,104 | 0,091 | 0,040 |
|  | **n** | 12 | 12 | 12 | 12 | 12 | 8 | 12 | 12 | 12 | 12 | 12 | 12 | 12 | 12 | 12 | 12 | 12 |
| **Th-P** | **r** | **0,03** | **0,12** | **0,07** | **0,16** | **0,1** | **0,22** | **0,08** | **0,46** | **0,02** | **-0,05** | **-0,07** | **0,05** | **0,29** | 1 | **0,05** | **0,01** | -0,120 |
|  | **p** | 0,912 | 0,679 | 0,801 | 0,593 | 0,736 | 0,563 | 0,794 | 0,100 | 0,944 | 0,865 | 0,823 | 0,853 | 0,369 |  | 0,862 | 0,985 | 0,683 |
|  | **n** | 14 | 14 | 14 | 14 | 14 | 9 | 14 | 14 | 14 | 14 | 13 | 14 | 12 | 14 | 14 | 13 | 14 |
| **Hy-o** | **r** | **0,35** | **.796**** | **.630*** | **0,49** | **.571*** | **0,39** | **.794**** | **.767**** | **.734**** | **.663**** | **.826**** | **.789**** | 0,493 | **0,05** | 1 | **.942**** | **-0,32** |
|  | **p** | 0,216 | 0,001 | 0,016 | 0,072 | 0,033 | 0,296 | 0,001 | 0,001 | 0,003 | 0,010 | 0,001 | 0,001 | 0,104 | 0,862 |  | 0,000 | 0,266 |
|  | **n** | 14 | 14 | 14 | 14 | 14 | 9 | 14 | 14 | 14 | 14 | 13 | 14 | 12 | 14 | 14 | 13 | 14 |
| **Hy-m** | **r** | **0,4** | **.834**** | **.647*** | **.559*** | **.670*** | **0,57** | **.865**** | **.750**** | **.779**** | **.647*** | **.841**** | **.760**** | 0,509 | **0,01** | **.942**** | 1 | **-0,24** |
|  | **p** | 0,181 | 0,000 | 0,017 | 0,047 | 0,012 | 0,138 | 0,000 | 0,003 | 0,002 | 0,017 | 0,000 | 0,003 | 0,091 | 0,985 | 0,000 |  | 0,422 |
|  | **n** | 13 | 13 | 13 | 13 | 13 | 8 | 13 | 13 | 13 | 13 | 13 | 13 | 12 | 13 | 13 | 13 | 13 |
| **Hy-l** | **r** | 0,288 | -0,028 | 0,276 | 0,012 | 0,090 | 0,164 | -0,199 | -0,391 | -0,214 | **-0,03** | -0,122 | **-0,4** | -.597* | -0,120 | **-0,32** | **-0,24** | 1 |
|  | **p** | 0,317 | 0,923 | 0,340 | 0,967 | 0,761 | 0,674 | 0,495 | 0,167 | 0,461 | 0,922 | 0,691 | 0,159 | 0,040 | 0,683 | 0,266 | 0,422 |  |
|  | **n** | 14 | 14 | 14 | 14 | 14 | 9 | 14 | 14 | 14 | 14 | 13 | 14 | 12 | 14 | 14 | 13 | 14 |

**Table S5. Results from correlation analysis of cell densities across regions for the D1 P35 group.** Two-tailed Pearson’s correlation (r) tests were used; pearson’s correlation coefficient (r), p value (p) and the number of subjects (n) is shown for each region pair.

|  |  | **MO** | **SS** | **GU/V** | **ACC** | **PFA** | **RSP** | **Olf** | **HR** | **CTX-s** | **STR** | **sAMY** | **PAL** | **Th-S** | **Th-P** | **Hy-o** | **Hy-m** | **Hy-l** |
| --- | --- | --- | --- | --- | --- | --- | --- | --- | --- | --- | --- | --- | --- | --- | --- | --- | --- | --- |
| **MO** | **r** | 1 | **.791**** | **.795**** | **0,45** | **.861**** | **0,56** | **0,45** | **0,04** | **0,18** | **0,32** | **0,16** | **-0,28** | 0,092 | **-0,22** | **0,22** | **0,27** | 0,322 |
|  | **p** |  | 0,002 | 0,002 | 0,140 | 0,001 | 0,092 | 0,141 | 0,911 | 0,584 | 0,318 | 0,636 | 0,381 | 0,800 | 0,538 | 0,488 | 0,422 | 0,307 |
|  | **n** | 12 | 12 | 12 | 12 | 10 | 10 | 12 | 12 | 12 | 12 | 11 | 12 | 10 | 10 | 12 | 11 | 12 |
| **SS** | **r** | **.791**** | 1 | **.895**** | **0,46** | **.665*** | **0,15** | **0,38** | **-0,06** | **0,12** | **0,46** | **0,22** | **-0,35** | -0,296 | **-0,57** | **0,05** | **0,26** | 0,333 |
|  | **p** | 0,002 |  | 0,000 | 0,133 | 0,036 | 0,685 | 0,225 | 0,862 | 0,699 | 0,129 | 0,515 | 0,264 | 0,406 | 0,088 | 0,883 | 0,444 | 0,290 |
|  | **n** | 12 | 12 | 12 | 12 | 10 | 10 | 12 | 12 | 12 | 12 | 11 | 12 | 10 | 10 | 12 | 11 | 12 |
| **GU/V** | **r** | **.795**** | **.895**** | 1 | **0,5** | **.805**** | **0,48** | **.577*** | **0,11** | **0,27** | **0,54** | **0,03** | **-0,17** | -0,249 | **-0,41** | **0,09** | **0,28** | 0,274 |
|  | **p** | 0,002 | 0,000 |  | 0,097 | 0,005 | 0,160 | 0,050 | 0,725 | 0,394 | 0,069 | 0,933 | 0,599 | 0,487 | 0,245 | 0,770 | 0,413 | 0,389 |
|  | **n** | 12 | 12 | 12 | 12 | 10 | 10 | 12 | 12 | 12 | 12 | 11 | 12 | 10 | 10 | 12 | 11 | 12 |
| **ACC** | **r** | **0,45** | **0,46** | **0,5** | 1 | **0,57** | **0,54** | **0,35** | **0,35** | **-0,12** | **0,15** | **-0,3** | **-0,18** | -0,269 | **0,03** | **0,2** | **.695*** | 0,503 |
|  | **p** | 0,140 | 0,133 | 0,097 |  | 0,084 | 0,103 | 0,266 | 0,271 | 0,720 | 0,635 | 0,376 | 0,570 | 0,453 | 0,937 | 0,532 | 0,017 | 0,095 |
|  | **n** | 12 | 12 | 12 | 12 | 10 | 10 | 12 | 12 | 12 | 12 | 11 | 12 | 10 | 10 | 12 | 11 | 12 |
| **PFA** | **r** | **.861**** | **.665*** | **.805**** | **0,57** | 1 | **.906**** | **.789**** | **0,27** | **0,11** | **0,39** | **-0,19** | **-0,21** | -0,248 | **-0,4** | **0,02** | **0,41** | 0,465 |
|  | **p** | 0,001 | 0,036 | 0,005 | 0,084 |  | 0,002 | 0,007 | 0,453 | 0,765 | 0,261 | 0,628 | 0,561 | 0,553 | 0,331 | 0,948 | 0,267 | 0,176 |
|  | **n** | 10 | 10 | 10 | 10 | 10 | 8 | 10 | 10 | 10 | 10 | 9 | 10 | 8 | 8 | 10 | 9 | 10 |
| **RSP** | **r** | **0,56** | **0,15** | **0,48** | **0,54** | **.906**** | 1 | **.756*** | **0,63** | **0,31** | **0,35** | **-0,44** | **0,08** | -0,062 | **-0** | **0,31** | **0,47** | 0,465 |
|  | **p** | 0,092 | 0,685 | 0,160 | 0,103 | 0,002 |  | 0,011 | 0,052 | 0,384 | 0,328 | 0,199 | 0,818 | 0,866 | 0,996 | 0,382 | 0,168 | 0,175 |
|  | **n** | 10 | 10 | 10 | 10 | 8 | 10 | 10 | 10 | 10 | 10 | 10 | 10 | 10 | 10 | 10 | 10 | 10 |
| **Olf** | **r** | **0,45** | **0,38** | **.577*** | **0,35** | **.789**** | **.756*** | 1 | **.586*** | **.681*** | **.869**** | **-0,03** | **0,3** | **0,29** | **0,3** | **.650*** | **.609*** | 0,211 |
|  | **p** | 0,141 | 0,225 | 0,050 | 0,266 | 0,007 | 0,011 |  | 0,045 | 0,015 | 0,000 | 0,941 | 0,341 | 0,423 | 0,394 | 0,022 | 0,047 | 0,511 |
|  | **n** | 12 | 12 | 12 | 12 | 10 | 10 | 12 | 12 | 12 | 12 | 11 | 12 | 10 | 10 | 12 | 11 | 12 |
| **HR** | **r** | **0,04** | **-0,06** | **0,11** | **0,35** | **0,27** | **0,63** | **.586*** | 1 | **.621*** | **0,46** | **-0,24** | **.751**** | **0,26** | **0,54** | **0,35** | **.907**** | 0,049 |
|  | **p** | 0,911 | 0,862 | 0,725 | 0,271 | 0,453 | 0,052 | 0,045 |  | 0,031 | 0,128 | 0,468 | 0,005 | 0,464 | 0,107 | 0,264 | 0,000 | 0,880 |
|  | **n** | 12 | 12 | 12 | 12 | 10 | 10 | 12 | 12 | 12 | 12 | 11 | 12 | 10 | 10 | 12 | 11 | 12 |
| **CTX-s** | **r** | **0,18** | **0,12** | **0,27** | **-0,12** | **0,11** | **0,31** | **.681*** | **.621*** | 1 | **.749**** | **0,51** | **.769**** | **.679*** | **0,49** | **0,51** | **0,19** | -0,427 |
|  | **p** | 0,584 | 0,699 | 0,394 | 0,720 | 0,765 | 0,384 | 0,015 | 0,031 |  | 0,005 | 0,110 | 0,003 | 0,031 | 0,149 | 0,087 | 0,572 | 0,166 |
|  | **n** | 12 | 12 | 12 | 12 | 10 | 10 | 12 | 12 | 12 | 12 | 11 | 12 | 10 | 10 | 12 | 11 | 12 |
| **STR** | **r** | **0,32** | **0,46** | **0,54** | **0,15** | **0,39** | **0,35** | **.869**** | **0,46** | **.749**** | 1 | **0,32** | **0,36** | **0,33** | **0,28** | **.608*** | **0,45** | **-0,04** |
|  | **p** | 0,318 | 0,129 | 0,069 | 0,635 | 0,261 | 0,328 | 0,000 | 0,128 | 0,005 |  | 0,331 | 0,246 | 0,349 | 0,431 | 0,036 | 0,163 | 0,890 |
|  | **n** | 12 | 12 | 12 | 12 | 10 | 10 | 12 | 12 | 12 | 12 | 11 | 12 | 10 | 10 | 12 | 11 | 12 |
| **sAMY** | **r** | **0,16** | **0,22** | **0,03** | **-0,3** | **-0,19** | **-0,44** | **-0,03** | **-0,24** | **0,51** | **0,32** | 1 | **0,36** | **.654*** | **0,36** | **0,23** | **-0,22** | -.704* |
|  | **p** | 0,636 | 0,515 | 0,933 | 0,376 | 0,628 | 0,199 | 0,941 | 0,468 | 0,110 | 0,331 |  | 0,283 | 0,040 | 0,304 | 0,500 | 0,524 | 0,016 |
|  | **n** | 11 | 11 | 11 | 11 | 9 | 10 | 11 | 11 | 11 | 11 | 11 | 11 | 10 | 10 | 11 | 11 | 11 |
| **PAL** | **r** | **-0,28** | **-0,35** | **-0,17** | **-0,18** | **-0,21** | **0,08** | **0,3** | **.751**** | **.769**** | **0,36** | **0,36** | 1 | **.839**** | **.910**** | **0,24** | **0,02** | **-0,55** |
|  | **p** | 0,381 | 0,264 | 0,599 | 0,570 | 0,561 | 0,818 | 0,341 | 0,005 | 0,003 | 0,246 | 0,283 |  | 0,002 | 0,000 | 0,453 | 0,960 | 0,065 |
|  | **n** | 12 | 12 | 12 | 12 | 10 | 10 | 12 | 12 | 12 | 12 | 11 | 12 | 10 | 10 | 12 | 11 | 12 |
| **Th-S** | **r** | 0,092 | -0,296 | -0,249 | -0,269 | -0,248 | -0,062 | **0,29** | **0,26** | **.679*** | **0,33** | **.654*** | **.839**** | 1 | **.881**** | .733* | 0,191 | -.633* |
|  | **p** | 0,800 | 0,406 | 0,487 | 0,453 | 0,553 | 0,866 | 0,423 | 0,464 | 0,031 | 0,349 | 0,040 | 0,002 |  | 0,001 | 0,016 | 0,597 | 0,049 |
|  | **n** | 10 | 10 | 10 | 10 | 8 | 10 | 10 | 10 | 10 | 10 | 10 | 10 | 10 | 10 | 10 | 10 | 10 |
| **Th-P** | **r** | **-0,22** | **-0,57** | **-0,41** | **0,03** | **-0,4** | **-0** | **0,3** | **0,54** | **0,49** | **0,28** | **0,36** | **.910**** | **.881**** | 1 | **.796**** | **0,52** | -0,421 |
|  | **p** | 0,538 | 0,088 | 0,245 | 0,937 | 0,331 | 0,996 | 0,394 | 0,107 | 0,149 | 0,431 | 0,304 | 0,000 | 0,001 |  | 0,006 | 0,124 | 0,225 |
|  | **n** | 10 | 10 | 10 | 10 | 8 | 10 | 10 | 10 | 10 | 10 | 10 | 10 | 10 | 10 | 10 | 10 | 10 |
| **Hy-o** | **r** | **0,22** | **0,05** | **0,09** | **0,2** | **0,02** | **0,31** | **.650*** | **0,35** | **0,51** | **.608*** | **0,23** | **0,24** | .733* | **.796**** | 1 | **.634*** | **0,03** |
|  | **p** | 0,488 | 0,883 | 0,770 | 0,532 | 0,948 | 0,382 | 0,022 | 0,264 | 0,087 | 0,036 | 0,500 | 0,453 | 0,016 | 0,006 |  | 0,036 | 0,915 |
|  | **n** | 12 | 12 | 12 | 12 | 10 | 10 | 12 | 12 | 12 | 12 | 11 | 12 | 10 | 10 | 12 | 11 | 12 |
| **Hy-m** | **r** | **0,27** | **0,26** | **0,28** | **.695*** | **0,41** | **0,47** | **.609*** | **.907**** | **0,19** | **0,45** | **-0,22** | **0,02** | 0,191 | **0,52** | **.634*** | 1 | **0,58** |
|  | **p** | 0,422 | 0,444 | 0,413 | 0,017 | 0,267 | 0,168 | 0,047 | 0,000 | 0,572 | 0,163 | 0,524 | 0,960 | 0,597 | 0,124 | 0,036 |  | 0,062 |
|  | **n** | 11 | 11 | 11 | 11 | 9 | 10 | 11 | 11 | 11 | 11 | 11 | 11 | 10 | 10 | 11 | 11 | 11 |
| **Hy-l** | **r** | 0,322 | 0,333 | 0,274 | 0,503 | 0,465 | 0,465 | 0,211 | 0,049 | -0,427 | **-0,04** | -.704* | **-0,55** | -.633* | -0,421 | **0,03** | **0,58** | 1 |
|  | **p** | 0,307 | 0,290 | 0,389 | 0,095 | 0,176 | 0,175 | 0,511 | 0,880 | 0,166 | 0,890 | 0,016 | 0,065 | 0,049 | 0,225 | 0,915 | 0,062 |  |
|  | **n** | 12 | 12 | 12 | 12 | 10 | 10 | 12 | 12 | 12 | 12 | 11 | 12 | 10 | 10 | 12 | 11 | 12 |

**Table S6. Results from correlation analysis of cell densities across regions for the D1 P49 group.** Two-tailed Pearson’s correlation (r) tests were used; pearson’s correlation coefficient (r), p value (p) and the number of subjects (n) is shown for each region pair.

|  |  | **MO** | **SS** | **GU/V** | **ACC** | **PFA** | **RSP** | **Olf** | **HR** | **CTX-s** | **STR** | **sAMY** | **PAL** | **Th-S** | **Th-P** | **Hy-o** | **Hy-m** | **Hy-l** |
| --- | --- | --- | --- | --- | --- | --- | --- | --- | --- | --- | --- | --- | --- | --- | --- | --- | --- | --- |
| **MO** | **r** | 1 | **0,36** | **0,48** | **0,3** | **0,4** | **0,3** | **0,42** | **-0,44** | **-0,02** | **0,56** | **-0,3** | **0,1** | -0,002 | **-0,31** | **-0,17** | **-0,36** | -0,285 |
|  | **p** |  | 0,344 | 0,156 | 0,401 | 0,251 | 0,438 | 0,232 | 0,241 | 0,961 | 0,089 | 0,402 | 0,786 | 0,996 | 0,384 | 0,638 | 0,336 | 0,424 |
|  | **n** | 10 | 9 | 10 | 10 | 10 | 9 | 10 | 9 | 10 | 10 | 10 | 10 | 10 | 10 | 10 | 9 | 10 |
| **SS** | **r** | **0,36** | 1 | **0,44** | **.694*** | **.724*** | **.966**** | **.756*** | **0,46** | **0,49** | **.702*** | **-0,06** | **.762*** | .678* | **0,47** | **0,27** | **0,14** | -0,010 |
|  | **p** | 0,344 |  | 0,207 | 0,026 | 0,018 | 0,000 | 0,011 | 0,212 | 0,148 | 0,023 | 0,871 | 0,010 | 0,031 | 0,170 | 0,456 | 0,728 | 0,978 |
|  | **n** | 9 | 10 | 10 | 10 | 10 | 9 | 10 | 9 | 10 | 10 | 10 | 10 | 10 | 10 | 10 | 9 | 10 |
| **GU/V** | **r** | **0,48** | **0,44** | 1 | **.740**** | **.695*** | **0,41** | **0,52** | **0,18** | **0,58** | **.805**** | **-0,06** | **0,36** | 0,386 | **0,36** | **0,08** | **0,14** | 0,132 |
|  | **p** | 0,156 | 0,207 |  | 0,009 | 0,018 | 0,243 | 0,099 | 0,615 | 0,064 | 0,003 | 0,865 | 0,278 | 0,242 | 0,272 | 0,805 | 0,697 | 0,698 |
|  | **n** | 10 | 10 | 11 | 11 | 11 | 10 | 11 | 10 | 11 | 11 | 11 | 11 | 11 | 11 | 11 | 10 | 11 |
| **ACC** | **r** | **0,3** | **.694*** | **.740**** | 1 | **.653*** | **.669*** | **0,45** | **0,55** | **.628*** | **.692*** | **0,01** | **0,59** | 0,483 | **0,45** | **0,45** | **0,43** | -0,093 |
|  | **p** | 0,401 | 0,026 | 0,009 |  | 0,029 | 0,035 | 0,170 | 0,102 | 0,039 | 0,018 | 0,985 | 0,054 | 0,132 | 0,161 | 0,167 | 0,219 | 0,786 |
|  | **n** | 10 | 10 | 11 | 11 | 11 | 10 | 11 | 10 | 11 | 11 | 11 | 11 | 11 | 11 | 11 | 10 | 11 |
| **PFA** | **r** | **0,4** | **.724*** | **.695*** | **.653*** | 1 | **.707*** | **0,56** | **0,4** | **0,35** | **.881**** | **-0,11** | **0,55** | 0,456 | **0,41** | **0,07** | **0,08** | 0,398 |
|  | **p** | 0,251 | 0,018 | 0,018 | 0,029 |  | 0,022 | 0,071 | 0,251 | 0,286 | 0,000 | 0,752 | 0,079 | 0,159 | 0,205 | 0,834 | 0,829 | 0,226 |
|  | **n** | 10 | 10 | 11 | 11 | 11 | 10 | 11 | 10 | 11 | 11 | 11 | 11 | 11 | 11 | 11 | 10 | 11 |
| **RSP** | **r** | **0,3** | **.966**** | **0,41** | **.669*** | **.707*** | 1 | **.748*** | **.714*** | **0,63** | **0,63** | **0,3** | **.911**** | .789** | **.652*** | **0,61** | **0,39** | -0,105 |
|  | **p** | 0,438 | 0,000 | 0,243 | 0,035 | 0,022 |  | 0,013 | 0,031 | 0,050 | 0,052 | 0,399 | 0,000 | 0,007 | 0,041 | 0,059 | 0,300 | 0,773 |
|  | **n** | 9 | 9 | 10 | 10 | 10 | 10 | 10 | 9 | 10 | 10 | 10 | 10 | 10 | 10 | 10 | 9 | 10 |
| **Olf** | **r** | **0,42** | **.756*** | **0,52** | **0,45** | **0,56** | **.748*** | 1 | **0,2** | **.639*** | **.602*** | **0,33** | **.682*** | **0,55** | **0,43** | **0,32** | **0,27** | -0,225 |
|  | **p** | 0,232 | 0,011 | 0,099 | 0,170 | 0,071 | 0,013 |  | 0,581 | 0,034 | 0,050 | 0,328 | 0,021 | 0,078 | 0,184 | 0,343 | 0,446 | 0,507 |
|  | **n** | 10 | 10 | 11 | 11 | 11 | 10 | 11 | 10 | 11 | 11 | 11 | 11 | 11 | 11 | 11 | 10 | 11 |
| **HR** | **r** | **-0,44** | **0,46** | **0,18** | **0,55** | **0,4** | **.714*** | **0,2** | 1 | **.702*** | **0,34** | **0,59** | **.864**** | **.725*** | **0,58** | **.793**** | **.736*** | 0,074 |
|  | **p** | 0,241 | 0,212 | 0,615 | 0,102 | 0,251 | 0,031 | 0,581 |  | 0,023 | 0,340 | 0,073 | 0,001 | 0,018 | 0,080 | 0,006 | 0,015 | 0,840 |
|  | **n** | 9 | 9 | 10 | 10 | 10 | 9 | 10 | 10 | 10 | 10 | 10 | 10 | 10 | 10 | 10 | 10 | 10 |
| **CTX-s** | **r** | **-0,02** | **0,49** | **0,58** | **.628*** | **0,35** | **0,63** | **.639*** | **.702*** | 1 | **0,47** | **0,6** | **.764**** | **.738**** | **.634*** | **.738**** | **.726*** | -0,190 |
|  | **p** | 0,961 | 0,148 | 0,064 | 0,039 | 0,286 | 0,050 | 0,034 | 0,023 |  | 0,140 | 0,053 | 0,006 | 0,010 | 0,036 | 0,009 | 0,018 | 0,576 |
|  | **n** | 10 | 10 | 11 | 11 | 11 | 10 | 11 | 10 | 11 | 11 | 11 | 11 | 11 | 11 | 11 | 10 | 11 |
| **STR** | **r** | **0,56** | **.702*** | **.805**** | **.692*** | **.881**** | **0,63** | **.602*** | **0,34** | **0,47** | 1 | **-0,05** | **.607*** | **0,33** | **0,19** | **0,09** | **0,22** | **0,35** |
|  | **p** | 0,089 | 0,023 | 0,003 | 0,018 | 0,000 | 0,052 | 0,050 | 0,340 | 0,140 |  | 0,888 | 0,048 | 0,327 | 0,581 | 0,783 | 0,550 | 0,285 |
|  | **n** | 10 | 10 | 11 | 11 | 11 | 10 | 11 | 10 | 11 | 11 | 11 | 11 | 11 | 11 | 11 | 10 | 11 |
| **sAMY** | **r** | **-0,3** | **-0,06** | **-0,06** | **0,01** | **-0,11** | **0,3** | **0,33** | **0,59** | **0,6** | **-0,05** | 1 | **0,55** | **0,3** | **0,15** | **.776**** | **.907**** | -0,296 |
|  | **p** | 0,402 | 0,871 | 0,865 | 0,985 | 0,752 | 0,399 | 0,328 | 0,073 | 0,053 | 0,888 |  | 0,078 | 0,369 | 0,667 | 0,005 | 0,000 | 0,377 |
|  | **n** | 10 | 10 | 11 | 11 | 11 | 10 | 11 | 10 | 11 | 11 | 11 | 11 | 11 | 11 | 11 | 10 | 11 |
| **PAL** | **r** | **0,1** | **.762*** | **0,36** | **0,59** | **0,55** | **.911**** | **.682*** | **.864**** | **.764**** | **.607*** | **0,55** | 1 | **.630*** | **0,42** | **.708*** | **.810**** | **-0,04** |
|  | **p** | 0,786 | 0,010 | 0,278 | 0,054 | 0,079 | 0,000 | 0,021 | 0,001 | 0,006 | 0,048 | 0,078 |  | 0,038 | 0,197 | 0,015 | 0,005 | 0,914 |
|  | **n** | 10 | 10 | 11 | 11 | 11 | 10 | 11 | 10 | 11 | 11 | 11 | 11 | 11 | 11 | 11 | 10 | 11 |
| **Th-S** | **r** | -0,002 | .678* | 0,386 | 0,483 | 0,456 | .789** | **0,55** | **.725*** | **.738**** | **0,33** | **0,3** | **.630*** | 1 | **.875**** | 0,567 | 0,302 | -0,154 |
|  | **p** | 0,996 | 0,031 | 0,242 | 0,132 | 0,159 | 0,007 | 0,078 | 0,018 | 0,010 | 0,327 | 0,369 | 0,038 |  | 0,000 | 0,069 | 0,396 | 0,650 |
|  | **n** | 10 | 10 | 11 | 11 | 11 | 10 | 11 | 10 | 11 | 11 | 11 | 11 | 11 | 11 | 11 | 10 | 11 |
| **Th-P** | **r** | **-0,31** | **0,47** | **0,36** | **0,45** | **0,41** | **.652*** | **0,43** | **0,58** | **.634*** | **0,19** | **0,15** | **0,42** | **.875**** | 1 | **0,37** | **0,16** | -0,009 |
|  | **p** | 0,384 | 0,170 | 0,272 | 0,161 | 0,205 | 0,041 | 0,184 | 0,080 | 0,036 | 0,581 | 0,667 | 0,197 | 0,000 |  | 0,265 | 0,649 | 0,978 |
|  | **n** | 10 | 10 | 11 | 11 | 11 | 10 | 11 | 10 | 11 | 11 | 11 | 11 | 11 | 11 | 11 | 10 | 11 |
| **Hy-o** | **r** | **-0,17** | **0,27** | **0,08** | **0,45** | **0,07** | **0,61** | **0,32** | **.793**** | **.738**** | **0,09** | **.776**** | **.708*** | 0,567 | **0,37** | 1 | **.880**** | **-0,48** |
|  | **p** | 0,638 | 0,456 | 0,805 | 0,167 | 0,834 | 0,059 | 0,343 | 0,006 | 0,009 | 0,783 | 0,005 | 0,015 | 0,069 | 0,265 |  | 0,001 | 0,138 |
|  | **n** | 10 | 10 | 11 | 11 | 11 | 10 | 11 | 10 | 11 | 11 | 11 | 11 | 11 | 11 | 11 | 10 | 11 |
| **Hy-m** | **r** | **-0,36** | **0,14** | **0,14** | **0,43** | **0,08** | **0,39** | **0,27** | **.736*** | **.726*** | **0,22** | **.907**** | **.810**** | 0,302 | **0,16** | **.880**** | 1 | **-0,18** |
|  | **p** | 0,336 | 0,728 | 0,697 | 0,219 | 0,829 | 0,300 | 0,446 | 0,015 | 0,018 | 0,550 | 0,000 | 0,005 | 0,396 | 0,649 | 0,001 |  | 0,616 |
|  | **n** | 9 | 9 | 10 | 10 | 10 | 9 | 10 | 10 | 10 | 10 | 10 | 10 | 10 | 10 | 10 | 10 | 10 |
| **Hy-l** | **r** | -0,285 | -0,010 | 0,132 | -0,093 | 0,398 | -0,105 | -0,225 | 0,074 | -0,190 | **0,35** | -0,296 | **-0,04** | -0,154 | -0,009 | **-0,48** | **-0,18** | 1 |
|  | **p** | 0,424 | 0,978 | 0,698 | 0,786 | 0,226 | 0,773 | 0,507 | 0,840 | 0,576 | 0,285 | 0,377 | 0,914 | 0,650 | 0,978 | 0,138 | 0,616 |  |
|  | **n** | 10 | 10 | 11 | 11 | 11 | 10 | 11 | 10 | 11 | 11 | 11 | 11 | 11 | 11 | 11 | 10 | 11 |

**Table S7. Results from correlation analysis of cell densities across regions for the D1 P70 group.** Two-tailed Pearson’s correlation (r) tests were used; pearson’s correlation coefficient (r), p value (p) and the number of subjects (n) is shown for each region pair.

|  |  | **MO** | **SS** | **GU/V** | **ACC** | **PFA** | **RSP** | **Olf** | **HR** | **CTX-s** | **STR** | **sAMY** | **PAL** | **Th-S** | **Th-P** | **Hy-o** | **Hy-m** | **Hy-l** |
| --- | --- | --- | --- | --- | --- | --- | --- | --- | --- | --- | --- | --- | --- | --- | --- | --- | --- | --- |
| **MO** | **r** | 1 | **.873**** | **.899**** | **.932**** | **.935**** | **.788*** | **.984**** | **0,68** | **0,62** | **.740*** | **0,33** | **.797*** | 0,679 | **0,63** | **.745*** | **0,15** | -0,008 |
|  | **p** |  | 0,005 | 0,002 | 0,001 | 0,001 | 0,035 | 0,000 | 0,066 | 0,102 | 0,036 | 0,430 | 0,018 | 0,138 | 0,131 | 0,034 | 0,742 | 0,985 |
|  | **n** | 8 | 8 | 8 | 8 | 8 | 7 | 8 | 8 | 8 | 8 | 8 | 8 | 6 | 7 | 8 | 7 | 8 |
| **SS** | **r** | **.873**** | 1 | **.978**** | **.735*** | **0,67** | **.937**** | **.826*** | **.899**** | **0,31** | **0,45** | **0,21** | **0,48** | .872* | **0,51** | **0,43** | **0,07** | -0,037 |
|  | **p** | 0,005 |  | 0,000 | 0,038 | 0,066 | 0,002 | 0,011 | 0,002 | 0,450 | 0,260 | 0,625 | 0,225 | 0,024 | 0,246 | 0,286 | 0,883 | 0,930 |
|  | **n** | 8 | 8 | 8 | 8 | 8 | 7 | 8 | 8 | 8 | 8 | 8 | 8 | 6 | 7 | 8 | 7 | 8 |
| **GU/V** | **r** | **.899**** | **.978**** | 1 | **.822*** | **.742*** | **.964**** | **.867**** | **.916**** | **0,44** | **0,53** | **0,23** | **0,56** | .895* | **0,57** | **0,53** | **-0,03** | 0,027 |
|  | **p** | 0,002 | 0,000 |  | 0,012 | 0,035 | 0,000 | 0,005 | 0,001 | 0,272 | 0,175 | 0,580 | 0,152 | 0,016 | 0,178 | 0,179 | 0,953 | 0,949 |
|  | **n** | 8 | 8 | 8 | 8 | 8 | 7 | 8 | 8 | 8 | 8 | 8 | 8 | 6 | 7 | 8 | 7 | 8 |
| **ACC** | **r** | **.932**** | **.735*** | **.822*** | 1 | **.973**** | **0,71** | **.966**** | **0,61** | **.795*** | **.875**** | **0,4** | **.905**** | 0,718 | **0,74** | **.867**** | **0,15** | 0,211 |
|  | **p** | 0,001 | 0,038 | 0,012 |  | 0,000 | 0,074 | 0,000 | 0,108 | 0,018 | 0,004 | 0,324 | 0,002 | 0,108 | 0,057 | 0,005 | 0,742 | 0,616 |
|  | **n** | 8 | 8 | 8 | 8 | 8 | 7 | 8 | 8 | 8 | 8 | 8 | 8 | 6 | 7 | 8 | 7 | 8 |
| **PFA** | **r** | **.935**** | **0,67** | **.742*** | **.973**** | 1 | **0,58** | **.970**** | **0,46** | **.774*** | **.912**** | **0,46** | **.932**** | 0,577 | **0,67** | **.905**** | **0,25** | 0,204 |
|  | **p** | 0,001 | 0,066 | 0,035 | 0,000 |  | 0,174 | 0,000 | 0,250 | 0,024 | 0,002 | 0,256 | 0,001 | 0,230 | 0,098 | 0,002 | 0,593 | 0,628 |
|  | **n** | 8 | 8 | 8 | 8 | 8 | 7 | 8 | 8 | 8 | 8 | 8 | 8 | 6 | 7 | 8 | 7 | 8 |
| **RSP** | **r** | **.788*** | **.937**** | **.964**** | **0,71** | **0,58** | 1 | **0,74** | **.968**** | **0,33** | **0,33** | **-0,1** | **0,45** | .882* | **0,64** | **0,31** | **-0,03** | -0,169 |
|  | **p** | 0,035 | 0,002 | 0,000 | 0,074 | 0,174 |  | 0,056 | 0,000 | 0,466 | 0,464 | 0,837 | 0,306 | 0,020 | 0,124 | 0,495 | 0,946 | 0,717 |
|  | **n** | 7 | 7 | 7 | 7 | 7 | 7 | 7 | 7 | 7 | 7 | 7 | 7 | 6 | 7 | 7 | 7 | 7 |
| **Olf** | **r** | **.984**** | **.826*** | **.867**** | **.966**** | **.970**** | **0,74** | 1 | **0,63** | **0,71** | **.843**** | **0,45** | **.870**** | **0,7** | **0,71** | **.838**** | **0,26** | 0,134 |
|  | **p** | 0,000 | 0,011 | 0,005 | 0,000 | 0,000 | 0,056 |  | 0,094 | 0,050 | 0,009 | 0,260 | 0,005 | 0,122 | 0,072 | 0,009 | 0,573 | 0,752 |
|  | **n** | 8 | 8 | 8 | 8 | 8 | 7 | 8 | 8 | 8 | 8 | 8 | 8 | 6 | 7 | 8 | 7 | 8 |
| **HR** | **r** | **0,68** | **.899**** | **.916**** | **0,61** | **0,46** | **.968**** | **0,63** | 1 | **0,29** | **0,26** | **0,06** | **0,31** | **.876*** | **0,51** | **0,26** | **-0,12** | -0,074 |
|  | **p** | 0,066 | 0,002 | 0,001 | 0,108 | 0,250 | 0,000 | 0,094 |  | 0,480 | 0,540 | 0,897 | 0,452 | 0,022 | 0,242 | 0,542 | 0,791 | 0,862 |
|  | **n** | 8 | 8 | 8 | 8 | 8 | 7 | 8 | 8 | 8 | 8 | 8 | 8 | 6 | 7 | 8 | 7 | 8 |
| **CTX-s** | **r** | **0,62** | **0,31** | **0,44** | **.795*** | **.774*** | **0,33** | **0,71** | **0,29** | 1 | **.831*** | **0,59** | **.879**** | **0,3** | **.823*** | **.909**** | **0,34** | 0,197 |
|  | **p** | 0,102 | 0,450 | 0,272 | 0,018 | 0,024 | 0,466 | 0,050 | 0,480 |  | 0,011 | 0,123 | 0,004 | 0,568 | 0,023 | 0,002 | 0,449 | 0,640 |
|  | **n** | 8 | 8 | 8 | 8 | 8 | 7 | 8 | 8 | 8 | 8 | 8 | 8 | 6 | 7 | 8 | 7 | 8 |
| **STR** | **r** | **.740*** | **0,45** | **0,53** | **.875**** | **.912**** | **0,33** | **.843**** | **0,26** | **.831*** | 1 | **0,69** | **.942**** | **0,49** | **0,72** | **.967**** | **0,48** | **0,51** |
|  | **p** | 0,036 | 0,260 | 0,175 | 0,004 | 0,002 | 0,464 | 0,009 | 0,540 | 0,011 |  | 0,057 | 0,000 | 0,324 | 0,070 | 0,000 | 0,270 | 0,201 |
|  | **n** | 8 | 8 | 8 | 8 | 8 | 7 | 8 | 8 | 8 | 8 | 8 | 8 | 6 | 7 | 8 | 7 | 8 |
| **sAMY** | **r** | **0,33** | **0,21** | **0,23** | **0,4** | **0,46** | **-0,1** | **0,45** | **0,06** | **0,59** | **0,69** | 1 | **0,52** | **0,19** | **0,55** | **.731*** | **.831*** | 0,583 |
|  | **p** | 0,430 | 0,625 | 0,580 | 0,324 | 0,256 | 0,837 | 0,260 | 0,897 | 0,123 | 0,057 |  | 0,187 | 0,723 | 0,196 | 0,039 | 0,021 | 0,129 |
|  | **n** | 8 | 8 | 8 | 8 | 8 | 7 | 8 | 8 | 8 | 8 | 8 | 8 | 6 | 7 | 8 | 7 | 8 |
| **PAL** | **r** | **.797*** | **0,48** | **0,56** | **.905**** | **.932**** | **0,45** | **.870**** | **0,31** | **.879**** | **.942**** | **0,52** | 1 | **0,45** | **.811*** | **.929**** | **0,46** | **0,22** |
|  | **p** | 0,018 | 0,225 | 0,152 | 0,002 | 0,001 | 0,306 | 0,005 | 0,452 | 0,004 | 0,000 | 0,187 |  | 0,370 | 0,027 | 0,001 | 0,301 | 0,599 |
|  | **n** | 8 | 8 | 8 | 8 | 8 | 7 | 8 | 8 | 8 | 8 | 8 | 8 | 6 | 7 | 8 | 7 | 8 |
| **Th-S** | **r** | 0,679 | .872* | .895* | 0,718 | 0,577 | .882* | **0,7** | **.876*** | **0,3** | **0,49** | **0,19** | **0,45** | 1 | **0,63** | 0,396 | 0,061 | 0,339 |
|  | **p** | 0,138 | 0,024 | 0,016 | 0,108 | 0,230 | 0,020 | 0,122 | 0,022 | 0,568 | 0,324 | 0,723 | 0,370 |  | 0,179 | 0,437 | 0,909 | 0,511 |
|  | **n** | 6 | 6 | 6 | 6 | 6 | 6 | 6 | 6 | 6 | 6 | 6 | 6 | 6 | 6 | 6 | 6 | 6 |
| **Th-P** | **r** | **0,63** | **0,51** | **0,57** | **0,74** | **0,67** | **0,64** | **0,71** | **0,51** | **.823*** | **0,72** | **0,55** | **.811*** | **0,63** | 1 | **0,75** | **0,5** | 0,120 |
|  | **p** | 0,131 | 0,246 | 0,178 | 0,057 | 0,098 | 0,124 | 0,072 | 0,242 | 0,023 | 0,070 | 0,196 | 0,027 | 0,179 |  | 0,052 | 0,251 | 0,797 |
|  | **n** | 7 | 7 | 7 | 7 | 7 | 7 | 7 | 7 | 7 | 7 | 7 | 7 | 6 | 7 | 7 | 7 | 7 |
| **Hy-o** | **r** | **.745*** | **0,43** | **0,53** | **.867**** | **.905**** | **0,31** | **.838**** | **0,26** | **.909**** | **.967**** | **.731*** | **.929**** | 0,396 | **0,75** | 1 | **0,45** | **0,39** |
|  | **p** | 0,034 | 0,286 | 0,179 | 0,005 | 0,002 | 0,495 | 0,009 | 0,542 | 0,002 | 0,000 | 0,039 | 0,001 | 0,437 | 0,052 |  | 0,312 | 0,344 |
|  | **n** | 8 | 8 | 8 | 8 | 8 | 7 | 8 | 8 | 8 | 8 | 8 | 8 | 6 | 7 | 8 | 7 | 8 |
| **Hy-m** | **r** | **0,15** | **0,07** | **-0,03** | **0,15** | **0,25** | **-0,03** | **0,26** | **-0,12** | **0,34** | **0,48** | **.831*** | **0,46** | 0,061 | **0,5** | **0,45** | 1 | **0,24** |
|  | **p** | 0,742 | 0,883 | 0,953 | 0,742 | 0,593 | 0,946 | 0,573 | 0,791 | 0,449 | 0,270 | 0,021 | 0,301 | 0,909 | 0,251 | 0,312 |  | 0,604 |
|  | **n** | 7 | 7 | 7 | 7 | 7 | 7 | 7 | 7 | 7 | 7 | 7 | 7 | 6 | 7 | 7 | 7 | 7 |
| **Hy-l** | **r** | -0,008 | -0,037 | 0,027 | 0,211 | 0,204 | -0,169 | 0,134 | -0,074 | 0,197 | **0,51** | 0,583 | **0,22** | 0,339 | 0,120 | **0,39** | **0,24** | 1 |
|  | **p** | 0,985 | 0,930 | 0,949 | 0,616 | 0,628 | 0,717 | 0,752 | 0,862 | 0,640 | 0,201 | 0,129 | 0,599 | 0,511 | 0,797 | 0,344 | 0,604 |  |
|  | **n** | 8 | 8 | 8 | 8 | 8 | 7 | 8 | 8 | 8 | 8 | 8 | 8 | 6 | 7 | 8 | 7 | 8 |

**Table S8. Results from correlation analysis of cell densities across regions for the D2 P17 group.** Two-tailed Pearson’s correlation (r) tests were used; pearson’s correlation coefficient (r), p value (p) and the number of subjects (n) is shown for each region pair.

|  |  | **MO** | **SS** | **GU/V** | **ACC** | **PFA** | **RSP** | **Olf** | **HR** | **CTX-s** | **STR** | **sAMY** | **PAL** | **Th-S** | **Th-P** | **Hy-o** | **Hy-m** | **Hy-l** |
| --- | --- | --- | --- | --- | --- | --- | --- | --- | --- | --- | --- | --- | --- | --- | --- | --- | --- | --- |
| **MO** | **r** | 1 | **0,74** | **0,02** | **0,08** | **0,71** | **0,87** | **0,7** | **0,56** | **0,4** | **.843*** | **0,68** | **0,56** | 0,436 | **0,05** | **.765*** | **0,17** | 0,469 |
|  | **p** |  | 0,056 | 0,975 | 0,861 | 0,075 | 0,054 | 0,082 | 0,193 | 0,372 | 0,017 | 0,094 | 0,188 | 0,328 | 0,911 | 0,045 | 0,708 | 0,288 |
|  | **n** | 7 | 7 | 6 | 7 | 7 | 5 | 7 | 7 | 7 | 7 | 7 | 7 | 7 | 7 | 7 | 7 | 7 |
| **SS** | **r** | **0,74** | 1 | **0,46** | **0,69** | **0,74** | **.918*** | **.945**** | **.823*** | **0,75** | **.804*** | **0,59** | **.903**** | 0,448 | **0,07** | **.890**** | **-0,06** | -0,145 |
|  | **p** | 0,056 |  | 0,364 | 0,085 | 0,059 | 0,028 | 0,001 | 0,023 | 0,054 | 0,029 | 0,164 | 0,005 | 0,314 | 0,878 | 0,007 | 0,899 | 0,757 |
|  | **n** | 7 | 7 | 6 | 7 | 7 | 5 | 7 | 7 | 7 | 7 | 7 | 7 | 7 | 7 | 7 | 7 | 7 |
| **GU/V** | **r** | **0,02** | **0,46** | 1 | **0,4** | **.888*** | **-0,53** | **0,45** | **0,49** | **0,81** | **0,51** | **0,6** | **0,29** | -0,669 | **0,37** | **0,25** | **0,64** | 0,179 |
|  | **p** | 0,975 | 0,364 |  | 0,435 | 0,018 | 0,467 | 0,369 | 0,325 | 0,051 | 0,304 | 0,210 | 0,573 | 0,146 | 0,477 | 0,626 | 0,173 | 0,735 |
|  | **n** | 6 | 6 | 6 | 6 | 6 | 4 | 6 | 6 | 6 | 6 | 6 | 6 | 6 | 6 | 6 | 6 | 6 |
| **ACC** | **r** | **0,08** | **0,69** | **0,4** | 1 | **0,41** | **0,65** | **0,64** | **0,7** | **0,61** | **0,25** | **0,15** | **0,61** | 0,015 | **0,13** | **0,42** | **-0,16** | -0,662 |
|  | **p** | 0,861 | 0,085 | 0,435 |  | 0,360 | 0,236 | 0,123 | 0,080 | 0,142 | 0,583 | 0,741 | 0,145 | 0,974 | 0,774 | 0,354 | 0,740 | 0,106 |
|  | **n** | 7 | 7 | 6 | 7 | 7 | 5 | 7 | 7 | 7 | 7 | 7 | 7 | 7 | 7 | 7 | 7 | 7 |
| **PFA** | **r** | **0,71** | **0,74** | **.888*** | **0,41** | 1 | **0,54** | **0,73** | **0,63** | **.835*** | **.886**** | **.829*** | **0,66** | -0,069 | **0,29** | **0,74** | **0,5** | 0,297 |
|  | **p** | 0,075 | 0,059 | 0,018 | 0,360 |  | 0,348 | 0,063 | 0,128 | 0,020 | 0,008 | 0,021 | 0,107 | 0,883 | 0,526 | 0,059 | 0,251 | 0,518 |
|  | **n** | 7 | 7 | 6 | 7 | 7 | 5 | 7 | 7 | 7 | 7 | 7 | 7 | 7 | 7 | 7 | 7 | 7 |
| **RSP** | **r** | **0,87** | **.918*** | **-0,53** | **0,65** | **0,54** | 1 | **0,84** | **0,72** | **0,45** | **0,73** | **0,45** | **0,83** | 0,762 | **-0,17** | **.896*** | **-0,23** | -0,044 |
|  | **p** | 0,054 | 0,028 | 0,467 | 0,236 | 0,348 |  | 0,073 | 0,174 | 0,445 | 0,158 | 0,448 | 0,085 | 0,134 | 0,789 | 0,040 | 0,714 | 0,944 |
|  | **n** | 5 | 5 | 4 | 5 | 5 | 5 | 5 | 5 | 5 | 5 | 5 | 5 | 5 | 5 | 5 | 5 | 5 |
| **Olf** | **r** | **0,7** | **.945**** | **0,45** | **0,64** | **0,73** | **0,84** | 1 | **.852*** | **.791*** | **.816*** | **0,75** | **.864*** | **0,43** | **0,29** | **.828*** | **-0,18** | -0,077 |
|  | **p** | 0,082 | 0,001 | 0,369 | 0,123 | 0,063 | 0,073 |  | 0,015 | 0,034 | 0,025 | 0,053 | 0,012 | 0,341 | 0,524 | 0,022 | 0,691 | 0,869 |
|  | **n** | 7 | 7 | 6 | 7 | 7 | 5 | 7 | 7 | 7 | 7 | 7 | 7 | 7 | 7 | 7 | 7 | 7 |
| **HR** | **r** | **0,56** | **.823*** | **0,49** | **0,7** | **0,63** | **0,72** | **.852*** | 1 | **0,65** | **0,54** | **0,54** | **0,63** | **0,07** | **0,57** | **0,52** | **-0,03** | -0,301 |
|  | **p** | 0,193 | 0,023 | 0,325 | 0,080 | 0,128 | 0,174 | 0,015 |  | 0,111 | 0,209 | 0,210 | 0,130 | 0,877 | 0,185 | 0,233 | 0,949 | 0,512 |
|  | **n** | 7 | 7 | 6 | 7 | 7 | 5 | 7 | 7 | 7 | 7 | 7 | 7 | 7 | 7 | 7 | 7 | 7 |
| **CTX-s** | **r** | **0,4** | **0,75** | **0,81** | **0,61** | **.835*** | **0,45** | **.791*** | **0,65** | 1 | **.778*** | **0,73** | **.842*** | **0,04** | **0,32** | **0,74** | **0,21** | -0,110 |
|  | **p** | 0,372 | 0,054 | 0,051 | 0,142 | 0,020 | 0,445 | 0,034 | 0,111 |  | 0,039 | 0,065 | 0,018 | 0,937 | 0,490 | 0,057 | 0,644 | 0,814 |
|  | **n** | 7 | 7 | 6 | 7 | 7 | 5 | 7 | 7 | 7 | 7 | 7 | 7 | 7 | 7 | 7 | 7 | 7 |
| **STR** | **r** | **.843*** | **.804*** | **0,51** | **0,25** | **.886**** | **0,73** | **.816*** | **0,54** | **.778*** | 1 | **.866*** | **.802*** | **0,37** | **0,09** | **.919**** | **0,22** | **0,39** |
|  | **p** | 0,017 | 0,029 | 0,304 | 0,583 | 0,008 | 0,158 | 0,025 | 0,209 | 0,039 |  | 0,012 | 0,030 | 0,415 | 0,850 | 0,003 | 0,640 | 0,387 |
|  | **n** | 7 | 7 | 6 | 7 | 7 | 5 | 7 | 7 | 7 | 7 | 7 | 7 | 7 | 7 | 7 | 7 | 7 |
| **sAMY** | **r** | **0,68** | **0,59** | **0,6** | **0,15** | **.829*** | **0,45** | **0,75** | **0,54** | **0,73** | **.866*** | 1 | **0,55** | **0,11** | **0,48** | **0,64** | **0,15** | 0,489 |
|  | **p** | 0,094 | 0,164 | 0,210 | 0,741 | 0,021 | 0,448 | 0,053 | 0,210 | 0,065 | 0,012 |  | 0,204 | 0,808 | 0,280 | 0,125 | 0,746 | 0,265 |
|  | **n** | 7 | 7 | 6 | 7 | 7 | 5 | 7 | 7 | 7 | 7 | 7 | 7 | 7 | 7 | 7 | 7 | 7 |
| **PAL** | **r** | **0,56** | **.903**** | **0,29** | **0,61** | **0,66** | **0,83** | **.864*** | **0,63** | **.842*** | **.802*** | **0,55** | 1 | **0,52** | **-0,06** | **.934**** | **-0,08** | **-0,2** |
|  | **p** | 0,188 | 0,005 | 0,573 | 0,145 | 0,107 | 0,085 | 0,012 | 0,130 | 0,018 | 0,030 | 0,204 |  | 0,228 | 0,904 | 0,002 | 0,861 | 0,666 |
|  | **n** | 7 | 7 | 6 | 7 | 7 | 5 | 7 | 7 | 7 | 7 | 7 | 7 | 7 | 7 | 7 | 7 | 7 |
| **Th-S** | **r** | 0,436 | 0,448 | -0,669 | 0,015 | -0,069 | 0,762 | **0,43** | **0,07** | **0,04** | **0,37** | **0,11** | **0,52** | 1 | **-0,48** | 0,587 | -0,634 | 0,021 |
|  | **p** | 0,328 | 0,314 | 0,146 | 0,974 | 0,883 | 0,134 | 0,341 | 0,877 | 0,937 | 0,415 | 0,808 | 0,228 |  | 0,279 | 0,166 | 0,126 | 0,964 |
|  | **n** | 7 | 7 | 6 | 7 | 7 | 5 | 7 | 7 | 7 | 7 | 7 | 7 | 7 | 7 | 7 | 7 | 7 |
| **Th-P** | **r** | **0,05** | **0,07** | **0,37** | **0,13** | **0,29** | **-0,17** | **0,29** | **0,57** | **0,32** | **0,09** | **0,48** | **-0,06** | **-0,48** | 1 | **-0,18** | **0,09** | -0,028 |
|  | **p** | 0,911 | 0,878 | 0,477 | 0,774 | 0,526 | 0,789 | 0,524 | 0,185 | 0,490 | 0,850 | 0,280 | 0,904 | 0,279 |  | 0,694 | 0,843 | 0,952 |
|  | **n** | 7 | 7 | 6 | 7 | 7 | 5 | 7 | 7 | 7 | 7 | 7 | 7 | 7 | 7 | 7 | 7 | 7 |
| **Hy-o** | **r** | **.765*** | **.890**** | **0,25** | **0,42** | **0,74** | **.896*** | **.828*** | **0,52** | **0,74** | **.919**** | **0,64** | **.934**** | 0,587 | **-0,18** | 1 | **0,03** | **0,12** |
|  | **p** | 0,045 | 0,007 | 0,626 | 0,354 | 0,059 | 0,040 | 0,022 | 0,233 | 0,057 | 0,003 | 0,125 | 0,002 | 0,166 | 0,694 |  | 0,949 | 0,798 |
|  | **n** | 7 | 7 | 6 | 7 | 7 | 5 | 7 | 7 | 7 | 7 | 7 | 7 | 7 | 7 | 7 | 7 | 7 |
| **Hy-m** | **r** | **0,17** | **-0,06** | **0,64** | **-0,16** | **0,5** | **-0,23** | **-0,18** | **-0,03** | **0,21** | **0,22** | **0,15** | **-0,08** | -0,634 | **0,09** | **0,03** | 1 | **0,35** |
|  | **p** | 0,708 | 0,899 | 0,173 | 0,740 | 0,251 | 0,714 | 0,691 | 0,949 | 0,644 | 0,640 | 0,746 | 0,861 | 0,126 | 0,843 | 0,949 |  | 0,435 |
|  | **n** | 7 | 7 | 6 | 7 | 7 | 5 | 7 | 7 | 7 | 7 | 7 | 7 | 7 | 7 | 7 | 7 | 7 |
| **Hy-l** | **r** | 0,469 | -0,145 | 0,179 | -0,662 | 0,297 | -0,044 | -0,077 | -0,301 | -0,110 | **0,39** | 0,489 | **-0,2** | 0,021 | -0,028 | **0,12** | **0,35** | 1 |
|  | **p** | 0,288 | 0,757 | 0,735 | 0,106 | 0,518 | 0,944 | 0,869 | 0,512 | 0,814 | 0,387 | 0,265 | 0,666 | 0,964 | 0,952 | 0,798 | 0,435 |  |
|  | **n** | 7 | 7 | 6 | 7 | 7 | 5 | 7 | 7 | 7 | 7 | 7 | 7 | 7 | 7 | 7 | 7 | 7 |

**Table S9. Results from correlation analysis of cell densities across regions for the D2 P25 group.** Two-tailed Pearson’s correlation (r) tests were used; pearson’s correlation coefficient (r), p value (p) and the number of subjects (n) is shown for each region pair.

|  |  | **MO** | **SS** | **GU/V** | **ACC** | **PFA** | **RSP** | **Olf** | **HR** | **CTX-s** | **STR** | **sAMY** | **PAL** | **Th-S** | **Th-P** | **Hy-o** | **Hy-m** | **Hy-l** |
| --- | --- | --- | --- | --- | --- | --- | --- | --- | --- | --- | --- | --- | --- | --- | --- | --- | --- | --- |
| **MO** | **r** | 1 | **.752*** | **0,53** | **.803**** | **.801**** | **0,37** | **.850**** | **.835**** | **0,55** | **.700*** | **0,27** | **.675*** | 0,406 | **0,32** | **.674*** | **0,64** | 0,569 |
|  | **p** |  | 0,012 | 0,119 | 0,005 | 0,005 | 0,410 | 0,002 | 0,003 | 0,102 | 0,024 | 0,444 | 0,032 | 0,278 | 0,400 | 0,033 | 0,063 | 0,110 |
|  | **n** | 10 | 10 | 10 | 10 | 10 | 7 | 10 | 10 | 10 | 10 | 10 | 10 | 9 | 9 | 10 | 9 | 9 |
| **SS** | **r** | **.752*** | 1 | **.693*** | **.718*** | **.689*** | **0,54** | **.867**** | **0,61** | **0,57** | **.801**** | **0,41** | **.805**** | 0,463 | **0,34** | **.662*** | **0,51** | 0,489 |
|  | **p** | 0,012 |  | 0,026 | 0,019 | 0,027 | 0,207 | 0,001 | 0,063 | 0,087 | 0,005 | 0,234 | 0,005 | 0,210 | 0,378 | 0,037 | 0,164 | 0,182 |
|  | **n** | 10 | 10 | 10 | 10 | 10 | 7 | 10 | 10 | 10 | 10 | 10 | 10 | 9 | 9 | 10 | 9 | 9 |
| **GU/V** | **r** | **0,53** | **.693*** | 1 | **0,29** | **.658*** | **.875**** | **.701*** | **0,31** | **.805**** | **.751*** | **.920**** | **0,57** | 0,337 | **0,2** | **.640*** | **0,32** | 0,380 |
|  | **p** | 0,119 | 0,026 |  | 0,411 | 0,039 | 0,010 | 0,024 | 0,384 | 0,005 | 0,012 | 0,000 | 0,085 | 0,375 | 0,598 | 0,046 | 0,409 | 0,312 |
|  | **n** | 10 | 10 | 10 | 10 | 10 | 7 | 10 | 10 | 10 | 10 | 10 | 10 | 9 | 9 | 10 | 9 | 9 |
| **ACC** | **r** | **.803**** | **.718*** | **0,29** | 1 | **.724*** | **0,2** | **.782**** | **.817**** | **0,39** | **0,57** | **0,06** | **.765**** | 0,425 | **0,3** | **0,57** | **0,4** | 0,336 |
|  | **p** | 0,005 | 0,019 | 0,411 |  | 0,018 | 0,661 | 0,008 | 0,004 | 0,270 | 0,083 | 0,870 | 0,010 | 0,255 | 0,437 | 0,086 | 0,281 | 0,376 |
|  | **n** | 10 | 10 | 10 | 10 | 10 | 7 | 10 | 10 | 10 | 10 | 10 | 10 | 9 | 9 | 10 | 9 | 9 |
| **PFA** | **r** | **.801**** | **.689*** | **.658*** | **.724*** | 1 | **0,65** | **.905**** | **0,52** | **.758*** | **.858**** | **0,49** | **.877**** | .667* | **0,53** | **.921**** | **0,61** | .677* |
|  | **p** | 0,005 | 0,027 | 0,039 | 0,018 |  | 0,116 | 0,000 | 0,122 | 0,011 | 0,001 | 0,152 | 0,001 | 0,050 | 0,142 | 0,000 | 0,080 | 0,045 |
|  | **n** | 10 | 10 | 10 | 10 | 10 | 7 | 10 | 10 | 10 | 10 | 10 | 10 | 9 | 9 | 10 | 9 | 9 |
| **RSP** | **r** | **0,37** | **0,54** | **.875**** | **0,2** | **0,65** | 1 | **0,72** | **-0,01** | **.877**** | **0,71** | **.792*** | **0,56** | 0,494 | **0,35** | **0,66** | **0,49** | 0,786 |
|  | **p** | 0,410 | 0,207 | 0,010 | 0,661 | 0,116 |  | 0,068 | 0,976 | 0,010 | 0,072 | 0,034 | 0,189 | 0,260 | 0,436 | 0,109 | 0,327 | 0,064 |
|  | **n** | 7 | 7 | 7 | 7 | 7 | 7 | 7 | 7 | 7 | 7 | 7 | 7 | 7 | 7 | 7 | 6 | 6 |
| **Olf** | **r** | **.850**** | **.867**** | **.701*** | **.782**** | **.905**** | **0,72** | 1 | **0,61** | **.810**** | **.882**** | **0,45** | **.853**** | **0,56** | **0,39** | **.867**** | **0,63** | 0,649 |
|  | **p** | 0,002 | 0,001 | 0,024 | 0,008 | 0,000 | 0,068 |  | 0,060 | 0,005 | 0,001 | 0,187 | 0,002 | 0,116 | 0,297 | 0,001 | 0,070 | 0,058 |
|  | **n** | 10 | 10 | 10 | 10 | 10 | 7 | 10 | 10 | 10 | 10 | 10 | 10 | 9 | 9 | 10 | 9 | 9 |
| **HR** | **r** | **.835**** | **0,61** | **0,31** | **.817**** | **0,52** | **-0,01** | **0,61** | 1 | **0,32** | **0,45** | **0,13** | **0,48** | **0,03** | **0,08** | **0,36** | **0,38** | 0,203 |
|  | **p** | 0,003 | 0,063 | 0,384 | 0,004 | 0,122 | 0,976 | 0,060 |  | 0,369 | 0,193 | 0,728 | 0,158 | 0,944 | 0,841 | 0,308 | 0,312 | 0,601 |
|  | **n** | 10 | 10 | 10 | 10 | 10 | 7 | 10 | 10 | 10 | 10 | 10 | 10 | 9 | 9 | 10 | 9 | 9 |
| **CTX-s** | **r** | **0,55** | **0,57** | **.805**** | **0,39** | **.758*** | **.877**** | **.810**** | **0,32** | 1 | **.793**** | **.699*** | **0,59** | **0,47** | **0,32** | **.833**** | **0,62** | 0,663 |
|  | **p** | 0,102 | 0,087 | 0,005 | 0,270 | 0,011 | 0,010 | 0,005 | 0,369 |  | 0,006 | 0,024 | 0,071 | 0,207 | 0,405 | 0,003 | 0,075 | 0,052 |
|  | **n** | 10 | 10 | 10 | 10 | 10 | 7 | 10 | 10 | 10 | 10 | 10 | 10 | 9 | 9 | 10 | 9 | 9 |
| **STR** | **r** | **.700*** | **.801**** | **.751*** | **0,57** | **.858**** | **0,71** | **.882**** | **0,45** | **.793**** | 1 | **0,61** | **.904**** | **0,62** | **.672*** | **.832**** | **0,57** | **.778*** |
|  | **p** | 0,024 | 0,005 | 0,012 | 0,083 | 0,001 | 0,072 | 0,001 | 0,193 | 0,006 |  | 0,059 | 0,000 | 0,077 | 0,048 | 0,003 | 0,110 | 0,013 |
|  | **n** | 10 | 10 | 10 | 10 | 10 | 7 | 10 | 10 | 10 | 10 | 10 | 10 | 9 | 9 | 10 | 9 | 9 |
| **sAMY** | **r** | **0,27** | **0,41** | **.920**** | **0,06** | **0,49** | **.792*** | **0,45** | **0,13** | **.699*** | **0,61** | 1 | **0,41** | **0,15** | **0,16** | **0,43** | **0** | 0,201 |
|  | **p** | 0,444 | 0,234 | 0,000 | 0,870 | 0,152 | 0,034 | 0,187 | 0,728 | 0,024 | 0,059 |  | 0,238 | 0,696 | 0,674 | 0,216 | 0,993 | 0,605 |
|  | **n** | 10 | 10 | 10 | 10 | 10 | 7 | 10 | 10 | 10 | 10 | 10 | 10 | 9 | 9 | 10 | 9 | 9 |
| **PAL** | **r** | **.675*** | **.805**** | **0,57** | **.765**** | **.877**** | **0,56** | **.853**** | **0,48** | **0,59** | **.904**** | **0,41** | 1 | **.708*** | **.719*** | **.784**** | **0,47** | **0,66** |
|  | **p** | 0,032 | 0,005 | 0,085 | 0,010 | 0,001 | 0,189 | 0,002 | 0,158 | 0,071 | 0,000 | 0,238 |  | 0,033 | 0,029 | 0,007 | 0,204 | 0,053 |
|  | **n** | 10 | 10 | 10 | 10 | 10 | 7 | 10 | 10 | 10 | 10 | 10 | 10 | 9 | 9 | 10 | 9 | 9 |
| **Th-S** | **r** | 0,406 | 0,463 | 0,337 | 0,425 | .667* | 0,494 | **0,56** | **0,03** | **0,47** | **0,62** | **0,15** | **.708*** | 1 | **.803**** | .755* | .802* | .846** |
|  | **p** | 0,278 | 0,210 | 0,375 | 0,255 | 0,050 | 0,260 | 0,116 | 0,944 | 0,207 | 0,077 | 0,696 | 0,033 |  | 0,009 | 0,019 | 0,017 | 0,008 |
|  | **n** | 9 | 9 | 9 | 9 | 9 | 7 | 9 | 9 | 9 | 9 | 9 | 9 | 9 | 9 | 9 | 8 | 8 |
| **Th-P** | **r** | **0,32** | **0,34** | **0,2** | **0,3** | **0,53** | **0,35** | **0,39** | **0,08** | **0,32** | **.672*** | **0,16** | **.719*** | **.803**** | 1 | **0,54** | **0,65** | .936** |
|  | **p** | 0,400 | 0,378 | 0,598 | 0,437 | 0,142 | 0,436 | 0,297 | 0,841 | 0,405 | 0,048 | 0,674 | 0,029 | 0,009 |  | 0,132 | 0,084 | 0,001 |
|  | **n** | 9 | 9 | 9 | 9 | 9 | 7 | 9 | 9 | 9 | 9 | 9 | 9 | 9 | 9 | 9 | 8 | 8 |
| **Hy-o** | **r** | **.674*** | **.662*** | **.640*** | **0,57** | **.921**** | **0,66** | **.867**** | **0,36** | **.833**** | **.832**** | **0,43** | **.784**** | .755* | **0,54** | 1 | **.798**** | **.771*** |
|  | **p** | 0,033 | 0,037 | 0,046 | 0,086 | 0,000 | 0,109 | 0,001 | 0,308 | 0,003 | 0,003 | 0,216 | 0,007 | 0,019 | 0,132 |  | 0,010 | 0,015 |
|  | **n** | 10 | 10 | 10 | 10 | 10 | 7 | 10 | 10 | 10 | 10 | 10 | 10 | 9 | 9 | 10 | 9 | 9 |
| **Hy-m** | **r** | **0,64** | **0,51** | **0,32** | **0,4** | **0,61** | **0,49** | **0,63** | **0,38** | **0,62** | **0,57** | **0** | **0,47** | .802* | **0,65** | **.798**** | 1 | **.820**** |
|  | **p** | 0,063 | 0,164 | 0,409 | 0,281 | 0,080 | 0,327 | 0,070 | 0,312 | 0,075 | 0,110 | 0,993 | 0,204 | 0,017 | 0,084 | 0,010 |  | 0,007 |
|  | **n** | 9 | 9 | 9 | 9 | 9 | 6 | 9 | 9 | 9 | 9 | 9 | 9 | 8 | 8 | 9 | 9 | 9 |
| **Hy-l** | **r** | 0,569 | 0,489 | 0,380 | 0,336 | .677* | 0,786 | 0,649 | 0,203 | 0,663 | **.778*** | 0,201 | **0,66** | .846** | .936** | **.771*** | **.820**** | 1 |
|  | **p** | 0,110 | 0,182 | 0,312 | 0,376 | 0,045 | 0,064 | 0,058 | 0,601 | 0,052 | 0,013 | 0,605 | 0,053 | 0,008 | 0,001 | 0,015 | 0,007 |  |
|  | **n** | 9 | 9 | 9 | 9 | 9 | 6 | 9 | 9 | 9 | 9 | 9 | 9 | 8 | 8 | 9 | 9 | 9 |

**Table S10. Results from correlation analysis of cell densities across regions for the D2 P35 group.** Two-tailed Pearson’s correlation (r) tests were used; pearson’s correlation coefficient (r), p value (p) and the number of subjects (n) is shown for each region pair.

|  |  | **MO** | **SS** | **GU/V** | **ACC** | **PFA** | **RSP** | **Olf** | **HR** | **CTX-s** | **STR** | **sAMY** | **PAL** | **Th-S** | **Th-P** | **Hy-o** | **Hy-m** | **Hy-l** |
| --- | --- | --- | --- | --- | --- | --- | --- | --- | --- | --- | --- | --- | --- | --- | --- | --- | --- | --- |
| **MO** | **r** | 1 | **.753*** | **.724*** | **.781*** | **0,65** | **-0,96** | **.744*** | **-0,39** | **.767*** | **0,45** | **.749*** | **0,04** | -0,039 | **0,57** | **0,37** | **.682*** | 0,566 |
|  | **p** |  | 0,019 | 0,028 | 0,013 | 0,056 | 0,179 | 0,021 | 0,304 | 0,016 | 0,224 | 0,020 | 0,934 | 0,921 | 0,111 | 0,321 | 0,043 | 0,112 |
|  | **n** | 9 | 9 | 9 | 9 | 9 | 3 | 9 | 9 | 9 | 9 | 9 | 8 | 9 | 9 | 9 | 9 | 9 |
| **SS** | **r** | **.753*** | 1 | **.831**** | **.955**** | **.864**** | **-0,39** | **0,58** | **0,15** | **.766**** | **.772**** | **.680*** | **0,56** | 0,010 | **.773**** | **0,54** | **.827**** | 0,632 |
|  | **p** | 0,019 |  | 0,003 | 0,000 | 0,001 | 0,744 | 0,078 | 0,683 | 0,010 | 0,009 | 0,030 | 0,118 | 0,979 | 0,009 | 0,110 | 0,003 | 0,050 |
|  | **n** | 9 | 10 | 10 | 10 | 10 | 3 | 10 | 10 | 10 | 10 | 10 | 9 | 10 | 10 | 10 | 10 | 10 |
| **GU/V** | **r** | **.724*** | **.831**** | 1 | **.799**** | **.732*** | **-0,04** | **0,61** | **-0,12** | **.973**** | **.883**** | **.930**** | **0,66** | 0,025 | **.653*** | **.779**** | **.805**** | .869** |
|  | **p** | 0,028 | 0,003 |  | 0,006 | 0,016 | 0,977 | 0,064 | 0,737 | 0,000 | 0,001 | 0,000 | 0,055 | 0,946 | 0,041 | 0,008 | 0,005 | 0,001 |
|  | **n** | 9 | 10 | 10 | 10 | 10 | 3 | 10 | 10 | 10 | 10 | 10 | 9 | 10 | 10 | 10 | 10 | 10 |
| **ACC** | **r** | **.781*** | **.955**** | **.799**** | 1 | **.944**** | **-0,78** | **0,63** | **0,25** | **.741*** | **.716*** | **0,63** | **0,53** | 0,110 | **.792**** | **0,49** | **.685*** | 0,615 |
|  | **p** | 0,013 | 0,000 | 0,006 |  | 0,000 | 0,432 | 0,052 | 0,484 | 0,014 | 0,020 | 0,051 | 0,140 | 0,763 | 0,006 | 0,155 | 0,029 | 0,059 |
|  | **n** | 9 | 10 | 10 | 10 | 10 | 3 | 10 | 10 | 10 | 10 | 10 | 9 | 10 | 10 | 10 | 10 | 10 |
| **PFA** | **r** | **0,65** | **.864**** | **.732*** | **.944**** | 1 | **-0,37** | **0,55** | **0,34** | **.636*** | **.690*** | **0,54** | **0,55** | 0,280 | **.725*** | **0,5** | **0,54** | 0,622 |
|  | **p** | 0,056 | 0,001 | 0,016 | 0,000 |  | 0,756 | 0,098 | 0,333 | 0,048 | 0,027 | 0,109 | 0,125 | 0,434 | 0,018 | 0,144 | 0,104 | 0,055 |
|  | **n** | 9 | 10 | 10 | 10 | 10 | 3 | 10 | 10 | 10 | 10 | 10 | 9 | 10 | 10 | 10 | 10 | 10 |
| **RSP** | **r** | **-0,96** | **-0,39** | **-0,04** | **-0,78** | **-0,37** | 1 | **-0,91** | **0,73** | **0,06** | **0** | **-0,65** | **0,09** | 0,629 | **-0,75** | **0,25** | **0,62** | 0,152 |
|  | **p** | 0,179 | 0,744 | 0,977 | 0,432 | 0,756 |  | 0,279 | 0,476 | 0,963 | 0,998 | 0,549 | 0,943 | 0,567 | 0,461 | 0,837 | 0,571 | 0,903 |
|  | **n** | 3 | 3 | 3 | 3 | 3 | 3 | 3 | 3 | 3 | 3 | 3 | 3 | 3 | 3 | 3 | 3 | 3 |
| **Olf** | **r** | **.744*** | **0,58** | **0,61** | **0,63** | **0,55** | **-0,91** | 1 | **0,04** | **0,55** | **.639*** | **.714*** | **.717*** | **0,22** | **.665*** | **0,47** | **0,43** | 0,298 |
|  | **p** | 0,021 | 0,078 | 0,064 | 0,052 | 0,098 | 0,279 |  | 0,910 | 0,096 | 0,047 | 0,020 | 0,030 | 0,541 | 0,036 | 0,170 | 0,215 | 0,403 |
|  | **n** | 9 | 10 | 10 | 10 | 10 | 3 | 10 | 10 | 10 | 10 | 10 | 9 | 10 | 10 | 10 | 10 | 10 |
| **HR** | **r** | **-0,39** | **0,15** | **-0,12** | **0,25** | **0,34** | **0,73** | **0,04** | 1 | **-0,19** | **0,08** | **-0,32** | **0,56** | **0,6** | **0,21** | **-0,04** | **-0,17** | -0,248 |
|  | **p** | 0,304 | 0,683 | 0,737 | 0,484 | 0,333 | 0,476 | 0,910 |  | 0,602 | 0,827 | 0,369 | 0,118 | 0,067 | 0,565 | 0,908 | 0,640 | 0,489 |
|  | **n** | 9 | 10 | 10 | 10 | 10 | 3 | 10 | 10 | 10 | 10 | 10 | 9 | 10 | 10 | 10 | 10 | 10 |
| **CTX-s** | **r** | **.767*** | **.766**** | **.973**** | **.741*** | **.636*** | **0,06** | **0,55** | **-0,19** | 1 | **.788**** | **.908**** | **0,47** | **-0,05** | **0,55** | **.769**** | **.800**** | .820** |
|  | **p** | 0,016 | 0,010 | 0,000 | 0,014 | 0,048 | 0,963 | 0,096 | 0,602 |  | 0,007 | 0,000 | 0,200 | 0,893 | 0,101 | 0,009 | 0,005 | 0,004 |
|  | **n** | 9 | 10 | 10 | 10 | 10 | 3 | 10 | 10 | 10 | 10 | 10 | 9 | 10 | 10 | 10 | 10 | 10 |
| **STR** | **r** | **0,45** | **.772**** | **.883**** | **.716*** | **.690*** | **0** | **.639*** | **0,08** | **.788**** | 1 | **.837**** | **.832**** | **0,18** | **.772**** | **.832**** | **.677*** | **.682*** |
|  | **p** | 0,224 | 0,009 | 0,001 | 0,020 | 0,027 | 0,998 | 0,047 | 0,827 | 0,007 |  | 0,003 | 0,005 | 0,625 | 0,009 | 0,003 | 0,031 | 0,030 |
|  | **n** | 9 | 10 | 10 | 10 | 10 | 3 | 10 | 10 | 10 | 10 | 10 | 9 | 10 | 10 | 10 | 10 | 10 |
| **sAMY** | **r** | **.749*** | **.680*** | **.930**** | **0,63** | **0,54** | **-0,65** | **.714*** | **-0,32** | **.908**** | **.837**** | 1 | **0,53** | **-0,11** | **0,55** | **.754*** | **.692*** | .793** |
|  | **p** | 0,020 | 0,030 | 0,000 | 0,051 | 0,109 | 0,549 | 0,020 | 0,369 | 0,000 | 0,003 |  | 0,140 | 0,764 | 0,103 | 0,012 | 0,027 | 0,006 |
|  | **n** | 9 | 10 | 10 | 10 | 10 | 3 | 10 | 10 | 10 | 10 | 10 | 9 | 10 | 10 | 10 | 10 | 10 |
| **PAL** | **r** | **0,04** | **0,56** | **0,66** | **0,53** | **0,55** | **0,09** | **.717*** | **0,56** | **0,47** | **.832**** | **0,53** | 1 | **0,63** | **.719*** | **0,56** | **0,42** | **0,11** |
|  | **p** | 0,934 | 0,118 | 0,055 | 0,140 | 0,125 | 0,943 | 0,030 | 0,118 | 0,200 | 0,005 | 0,140 |  | 0,066 | 0,029 | 0,115 | 0,266 | 0,777 |
|  | **n** | 8 | 9 | 9 | 9 | 9 | 3 | 9 | 9 | 9 | 9 | 9 | 9 | 9 | 9 | 9 | 9 | 9 |
| **Th-S** | **r** | -0,039 | 0,010 | 0,025 | 0,110 | 0,280 | 0,629 | **0,22** | **0,6** | **-0,05** | **0,18** | **-0,11** | **0,63** | 1 | **0,31** | 0,102 | 0,013 | -0,073 |
|  | **p** | 0,921 | 0,979 | 0,946 | 0,763 | 0,434 | 0,567 | 0,541 | 0,067 | 0,893 | 0,625 | 0,764 | 0,066 |  | 0,387 | 0,779 | 0,971 | 0,841 |
|  | **n** | 9 | 10 | 10 | 10 | 10 | 3 | 10 | 10 | 10 | 10 | 10 | 9 | 10 | 10 | 10 | 10 | 10 |
| **Th-P** | **r** | **0,57** | **.773**** | **.653*** | **.792**** | **.725*** | **-0,75** | **.665*** | **0,21** | **0,55** | **.772**** | **0,55** | **.719*** | **0,31** | 1 | **0,41** | **0,61** | 0,414 |
|  | **p** | 0,111 | 0,009 | 0,041 | 0,006 | 0,018 | 0,461 | 0,036 | 0,565 | 0,101 | 0,009 | 0,103 | 0,029 | 0,387 |  | 0,238 | 0,060 | 0,234 |
|  | **n** | 9 | 10 | 10 | 10 | 10 | 3 | 10 | 10 | 10 | 10 | 10 | 9 | 10 | 10 | 10 | 10 | 10 |
| **Hy-o** | **r** | **0,37** | **0,54** | **.779**** | **0,49** | **0,5** | **0,25** | **0,47** | **-0,04** | **.769**** | **.832**** | **.754*** | **0,56** | 0,102 | **0,41** | 1 | **0,54** | **0,54** |
|  | **p** | 0,321 | 0,110 | 0,008 | 0,155 | 0,144 | 0,837 | 0,170 | 0,908 | 0,009 | 0,003 | 0,012 | 0,115 | 0,779 | 0,238 |  | 0,107 | 0,105 |
|  | **n** | 9 | 10 | 10 | 10 | 10 | 3 | 10 | 10 | 10 | 10 | 10 | 9 | 10 | 10 | 10 | 10 | 10 |
| **Hy-m** | **r** | **.682*** | **.827**** | **.805**** | **.685*** | **0,54** | **0,62** | **0,43** | **-0,17** | **.800**** | **.677*** | **.692*** | **0,42** | 0,013 | **0,61** | **0,54** | 1 | **0,61** |
|  | **p** | 0,043 | 0,003 | 0,005 | 0,029 | 0,104 | 0,571 | 0,215 | 0,640 | 0,005 | 0,031 | 0,027 | 0,266 | 0,971 | 0,060 | 0,107 |  | 0,060 |
|  | **n** | 9 | 10 | 10 | 10 | 10 | 3 | 10 | 10 | 10 | 10 | 10 | 9 | 10 | 10 | 10 | 10 | 10 |
| **Hy-l** | **r** | 0,566 | 0,632 | .869** | 0,615 | 0,622 | 0,152 | 0,298 | -0,248 | .820** | **.682*** | .793** | **0,11** | -0,073 | 0,414 | **0,54** | **0,61** | 1 |
|  | **p** | 0,112 | 0,050 | 0,001 | 0,059 | 0,055 | 0,903 | 0,403 | 0,489 | 0,004 | 0,030 | 0,006 | 0,777 | 0,841 | 0,234 | 0,105 | 0,060 |  |
|  | **n** | 9 | 10 | 10 | 10 | 10 | 3 | 10 | 10 | 10 | 10 | 10 | 9 | 10 | 10 | 10 | 10 | 10 |

**Table S11. Results from correlation analysis of cell densities across regions for the D2 P49 group.** Two-tailed Pearson’s correlation (r) tests were used; pearson’s correlation coefficient (r), p value (p) and the number of subjects (n) is shown for each region pair.

|  |  | **MO** | **SS** | **GU/V** | **ACC** | **PFA** | **RSP** | **Olf** | **HR** | **CTX-s** | **STR** | **sAMY** | **PAL** | **Th-S** | **Th-P** | **Hy-o** | **Hy-m** | **Hy-l** |
| --- | --- | --- | --- | --- | --- | --- | --- | --- | --- | --- | --- | --- | --- | --- | --- | --- | --- | --- |
| **MO** | **r** | 1 | **0,35** | **0,41** | **.766*** | **0,51** | **0,57** | **-0,12** | **0,44** | **0,65** | **0,04** | **0,05** | **0,49** | 0,350 | **0,4** | **0,33** | **0,5** | 0,510 |
|  | **p** |  | 0,349 | 0,276 | 0,027 | 0,161 | 0,315 | 0,783 | 0,234 | 0,057 | 0,914 | 0,901 | 0,179 | 0,395 | 0,292 | 0,389 | 0,207 | 0,243 |
|  | **n** | 9 | 9 | 9 | 8 | 9 | 5 | 8 | 9 | 9 | 9 | 8 | 9 | 8 | 9 | 9 | 8 | 7 |
| **SS** | **r** | **0,35** | 1 | **0,65** | **0,67** | **0,62** | **0,85** | **0,19** | **0,64** | **0,56** | **0,09** | **0,22** | **0,43** | .830* | **.739*** | **0,48** | **0,67** | -0,064 |
|  | **p** | 0,349 |  | 0,057 | 0,068 | 0,074 | 0,071 | 0,647 | 0,065 | 0,114 | 0,816 | 0,599 | 0,243 | 0,011 | 0,023 | 0,187 | 0,069 | 0,891 |
|  | **n** | 9 | 9 | 9 | 8 | 9 | 5 | 8 | 9 | 9 | 9 | 8 | 9 | 8 | 9 | 9 | 8 | 7 |
| **GU/V** | **r** | **0,41** | **0,65** | 1 | **0,65** | **.886**** | **.882*** | **0,49** | **0,6** | **.714*** | **.667*** | **0,62** | **.772*** | 0,530 | **.899**** | **.744*** | **0,34** | 0,703 |
|  | **p** | 0,276 | 0,057 |  | 0,079 | 0,001 | 0,048 | 0,218 | 0,087 | 0,031 | 0,050 | 0,099 | 0,015 | 0,177 | 0,001 | 0,021 | 0,412 | 0,078 |
|  | **n** | 9 | 9 | 9 | 8 | 9 | 5 | 8 | 9 | 9 | 9 | 8 | 9 | 8 | 9 | 9 | 8 | 7 |
| **ACC** | **r** | **.766*** | **0,67** | **0,65** | 1 | **0,68** | **0,76** | **0,32** | **.833*** | **0,7** | **0,22** | **0,21** | **.707*** | 0,521 | **0,66** | **0,54** | **0,62** | 0,534 |
|  | **p** | 0,027 | 0,068 | 0,079 |  | 0,064 | 0,140 | 0,480 | 0,010 | 0,055 | 0,596 | 0,648 | 0,050 | 0,230 | 0,077 | 0,167 | 0,139 | 0,275 |
|  | **n** | 8 | 8 | 8 | 8 | 8 | 5 | 7 | 8 | 8 | 8 | 7 | 8 | 7 | 8 | 8 | 7 | 6 |
| **PFA** | **r** | **0,51** | **0,62** | **.886**** | **0,68** | 1 | **0,65** | **0,63** | **0,6** | **.940**** | **.686*** | **0,31** | **.871**** | 0,582 | **.800**** | **.874**** | **0,45** | 0,678 |
|  | **p** | 0,161 | 0,074 | 0,001 | 0,064 |  | 0,231 | 0,093 | 0,088 | 0,000 | 0,041 | 0,454 | 0,002 | 0,130 | 0,010 | 0,002 | 0,268 | 0,094 |
|  | **n** | 9 | 9 | 9 | 8 | 9 | 5 | 8 | 9 | 9 | 9 | 8 | 9 | 8 | 9 | 9 | 8 | 7 |
| **RSP** | **r** | **0,57** | **0,85** | **.882*** | **0,76** | **0,65** | 1 | **0,44** | **0,49** | **0,48** | **0,4** | **0,94** | **0,47** | 0,620 | **0,83** | **0,37** | **-0,49** | 0,022 |
|  | **p** | 0,315 | 0,071 | 0,048 | 0,140 | 0,231 |  | 0,557 | 0,402 | 0,411 | 0,502 | 0,061 | 0,429 | 0,264 | 0,082 | 0,542 | 0,513 | 0,978 |
|  | **n** | 5 | 5 | 5 | 5 | 5 | 5 | 4 | 5 | 5 | 5 | 4 | 5 | 5 | 5 | 5 | 4 | 4 |
| **Olf** | **r** | **-0,12** | **0,19** | **0,49** | **0,32** | **0,63** | **0,44** | 1 | **0,56** | **0,42** | **0,52** | **0,42** | **0,68** | **-0,03** | **0,6** | **.780*** | **-0,47** | 0,612 |
|  | **p** | 0,783 | 0,647 | 0,218 | 0,480 | 0,093 | 0,557 |  | 0,146 | 0,294 | 0,186 | 0,354 | 0,062 | 0,955 | 0,112 | 0,022 | 0,289 | 0,196 |
|  | **n** | 8 | 8 | 8 | 7 | 8 | 4 | 8 | 8 | 8 | 8 | 7 | 8 | 7 | 8 | 8 | 7 | 6 |
| **HR** | **r** | **0,44** | **0,64** | **0,6** | **.833*** | **0,6** | **0,49** | **0,56** | 1 | **0,56** | **0,37** | **0,01** | **.687*** | **0,63** | **.686*** | **.728*** | **0,43** | 0,547 |
|  | **p** | 0,234 | 0,065 | 0,087 | 0,010 | 0,088 | 0,402 | 0,146 |  | 0,113 | 0,327 | 0,986 | 0,041 | 0,094 | 0,041 | 0,026 | 0,291 | 0,204 |
|  | **n** | 9 | 9 | 9 | 8 | 9 | 5 | 8 | 9 | 9 | 9 | 8 | 9 | 8 | 9 | 9 | 8 | 7 |
| **CTX-s** | **r** | **0,65** | **0,56** | **.714*** | **0,7** | **.940**** | **0,48** | **0,42** | **0,56** | 1 | **0,52** | **0** | **.808**** | **0,64** | **0,65** | **.809**** | **0,5** | 0,594 |
|  | **p** | 0,057 | 0,114 | 0,031 | 0,055 | 0,000 | 0,411 | 0,294 | 0,113 |  | 0,149 | 0,995 | 0,008 | 0,085 | 0,059 | 0,008 | 0,205 | 0,160 |
|  | **n** | 9 | 9 | 9 | 8 | 9 | 5 | 8 | 9 | 9 | 9 | 8 | 9 | 8 | 9 | 9 | 8 | 7 |
| **STR** | **r** | **0,04** | **0,09** | **.667*** | **0,22** | **.686*** | **0,4** | **0,52** | **0,37** | **0,52** | 1 | **0,36** | **.816**** | **0,1** | **0,49** | **.812**** | **0,1** | **.979**** |
|  | **p** | 0,914 | 0,816 | 0,050 | 0,596 | 0,041 | 0,502 | 0,186 | 0,327 | 0,149 |  | 0,385 | 0,007 | 0,821 | 0,183 | 0,008 | 0,815 | 0,000 |
|  | **n** | 9 | 9 | 9 | 8 | 9 | 5 | 8 | 9 | 9 | 9 | 8 | 9 | 8 | 9 | 9 | 8 | 7 |
| **sAMY** | **r** | **0,05** | **0,22** | **0,62** | **0,21** | **0,31** | **0,94** | **0,42** | **0,01** | **0** | **0,36** | 1 | **0,21** | **-0,14** | **0,58** | **0,12** | **-0,34** | -0,158 |
|  | **p** | 0,901 | 0,599 | 0,099 | 0,648 | 0,454 | 0,061 | 0,354 | 0,986 | 0,995 | 0,385 |  | 0,625 | 0,763 | 0,133 | 0,783 | 0,451 | 0,765 |
|  | **n** | 8 | 8 | 8 | 7 | 8 | 4 | 7 | 8 | 8 | 8 | 8 | 8 | 7 | 8 | 8 | 7 | 6 |
| **PAL** | **r** | **0,49** | **0,43** | **.772*** | **.707*** | **.871**** | **0,47** | **0,68** | **.687*** | **.808**** | **.816**** | **0,21** | 1 | **0,39** | **0,6** | **.882**** | **0,55** | **.922**** |
|  | **p** | 0,179 | 0,243 | 0,015 | 0,050 | 0,002 | 0,429 | 0,062 | 0,041 | 0,008 | 0,007 | 0,625 |  | 0,336 | 0,085 | 0,002 | 0,154 | 0,003 |
|  | **n** | 9 | 9 | 9 | 8 | 9 | 5 | 8 | 9 | 9 | 9 | 8 | 9 | 8 | 9 | 9 | 8 | 7 |
| **Th-S** | **r** | 0,350 | .830* | 0,530 | 0,521 | 0,582 | 0,620 | **-0,03** | **0,63** | **0,64** | **0,1** | **-0,14** | **0,39** | 1 | **0,57** | 0,515 | 0,549 | -0,077 |
|  | **p** | 0,395 | 0,011 | 0,177 | 0,230 | 0,130 | 0,264 | 0,955 | 0,094 | 0,085 | 0,821 | 0,763 | 0,336 |  | 0,144 | 0,192 | 0,202 | 0,885 |
|  | **n** | 8 | 8 | 8 | 7 | 8 | 5 | 7 | 8 | 8 | 8 | 7 | 8 | 8 | 8 | 8 | 7 | 6 |
| **Th-P** | **r** | **0,4** | **.739*** | **.899**** | **0,66** | **.800**** | **0,83** | **0,6** | **.686*** | **0,65** | **0,49** | **0,58** | **0,6** | **0,57** | 1 | **.741*** | **0,22** | 0,346 |
|  | **p** | 0,292 | 0,023 | 0,001 | 0,077 | 0,010 | 0,082 | 0,112 | 0,041 | 0,059 | 0,183 | 0,133 | 0,085 | 0,144 |  | 0,022 | 0,593 | 0,448 |
|  | **n** | 9 | 9 | 9 | 8 | 9 | 5 | 8 | 9 | 9 | 9 | 8 | 9 | 8 | 9 | 9 | 8 | 7 |
| **Hy-o** | **r** | **0,33** | **0,48** | **.744*** | **0,54** | **.874**** | **0,37** | **.780*** | **.728*** | **.809**** | **.812**** | **0,12** | **.882**** | 0,515 | **.741*** | 1 | **0,37** | **.802*** |
|  | **p** | 0,389 | 0,187 | 0,021 | 0,167 | 0,002 | 0,542 | 0,022 | 0,026 | 0,008 | 0,008 | 0,783 | 0,002 | 0,192 | 0,022 |  | 0,368 | 0,030 |
|  | **n** | 9 | 9 | 9 | 8 | 9 | 5 | 8 | 9 | 9 | 9 | 8 | 9 | 8 | 9 | 9 | 8 | 7 |
| **Hy-m** | **r** | **0,5** | **0,67** | **0,34** | **0,62** | **0,45** | **-0,49** | **-0,47** | **0,43** | **0,5** | **0,1** | **-0,34** | **0,55** | 0,549 | **0,22** | **0,37** | 1 | **0,22** |
|  | **p** | 0,207 | 0,069 | 0,412 | 0,139 | 0,268 | 0,513 | 0,289 | 0,291 | 0,205 | 0,815 | 0,451 | 0,154 | 0,202 | 0,593 | 0,368 |  | 0,628 |
|  | **n** | 8 | 8 | 8 | 7 | 8 | 4 | 7 | 8 | 8 | 8 | 7 | 8 | 7 | 8 | 8 | 8 | 7 |
| **Hy-l** | **r** | 0,510 | -0,064 | 0,703 | 0,534 | 0,678 | 0,022 | 0,612 | 0,547 | 0,594 | **.979**** | -0,158 | **.922**** | -0,077 | 0,346 | **.802*** | **0,22** | 1 |
|  | **p** | 0,243 | 0,891 | 0,078 | 0,275 | 0,094 | 0,978 | 0,196 | 0,204 | 0,160 | 0,000 | 0,765 | 0,003 | 0,885 | 0,448 | 0,030 | 0,628 |  |
|  | **n** | 7 | 7 | 7 | 6 | 7 | 4 | 6 | 7 | 7 | 7 | 6 | 7 | 6 | 7 | 7 | 7 | 7 |

**Table S12. Results from correlation analysis of cell densities across regions for the D2 P70 group.** Two-tailed Pearson’s correlation (r) tests were used; pearson’s correlation coefficient (r), p value (p) and the number of subjects (n) is shown for each region pair.

| **Genotype** | **Age** | **Sex** | **Raw dataset DOI** | **# of subjects analyzed** |
| --- | --- | --- | --- | --- |
| D1 | P70 | M | 10.25493/AVRZ-4JB | 5 |
|  |  | F | 10.25493/5MXR-AW7 | 6 |
|  | P49 | M | 10.25493/GVFP-10X | 7 |
|  |  | F | 10.25493/31D4-SKG | 5 |
|  | P35 | M | 10.25493/37DN-V7S | 9 |
|  |  | F | 10.25493/F3QR-M3S | 5 |
|  | P25 | M | 10.25493/AMBF-6CV | 6 |
|  |  | F | 10.25493/C3A9-VVM | 8 |
|  | P17 | M | 10.25493/TJK8-D7W | 7 |
|  |  | F | 10.25493/V09E-521 | 9 |
| D2 | P70 | M | 10.25493/4DEB-5AJ | 5 |
|  |  | F | 10.25493/VTD0-D15 | 4 |
|  | P49 | M | 10.25493/ANPQ-05J | 5 |
|  |  | F | 10.25493/JJYX-T5R | 5 |
|  | P35 | M | 10.25493/MFF5-KY5 | 4 |
|  |  | F | 10.25493/1J85-QSK | 6 |
|  | P25 | M | 10.25493/4G63-GDE | 4 |
|  |  | F | 10.25493/C3A9-VVM | 3 |
|  | P17 | M | 10.25493/9DHD-DXZ | 3 |
|  |  | F | 10.25493/G5VR-63E | 5 |

**Table S13. Data used in the analyses.** We quantified D1 and D2 receptor positive cells in five age groups across males and females. The DOIs of the datasets containing the raw data (high-resolution tiff images) are given, these datasets are publicly available through the EBRAINS Knowledge Graph.

| **Region name (custom regions used in QUINT analysis)** | **Major brain region** |
| --- | --- |
| Frontal pole, cerebral cortex | Motor areas |
| Primary motor area | Motor areas |
| Secondary motor area | Motor areas |
| Primary somatosensory area, nose | Somatosensory areas |
| Primary somatosensory area, barrel field | Somatosensory areas |
| Primary somatosensory area, lower limb | Somatosensory areas |
| Primary somatosensory area, mouth | Somatosensory areas |
| Primary somatosensory area, upper limb | Somatosensory areas |
| Primary somatosensory area, trunk | Somatosensory areas |
| Primary somatosensory area, unassigned | Somatosensory areas |
| Supplemental somatosensory area | Somatosensory areas |
| Gustatory areas | Gustatory and visceral areas |
| Visceral area | Gustatory and visceral areas |
| Dorsal auditory area | Auditory areas |
| Primary auditory area | Auditory areas |
| Posterior auditory area | Auditory areas |
| Ventral auditory area | Auditory areas |
| Anterolateral visual area | Visual areas |
| Anteromedial visual area | Visual areas |
| Lateral visual area | Visual areas |
| Primary visual area | Visual areas |
| Posterolateral visual area | Visual areas |
| Posteromedial visual area | Visual areas |
| Laterointermediate area | Visual areas |
| Postrhinal area | Visual areas |
| Anterior cingulate area, dorsal part | Anterior cingulate areas |
| Anterior cingulate area, ventral part | Anterior cingulate areas |
| Prelimbic area | Prefrontal areas |
| Infralimbic area | Prefrontal areas |
| Orbital area | Prefrontal areas |
| Orbital area, lateral part | Prefrontal areas |
| Orbital area, medial part | Prefrontal areas |
| Orbital area, ventrolateral part | Prefrontal areas |
| Agranular insular area, dorsal part | Prefrontal areas |
| Agranular insular area, posterior part | Prefrontal areas |
| Agranular insular area, ventral part | Prefrontal areas |
| Retrosplenial area, lateral agranular part | Retrosplenial areas |
| Retrosplenial area, dorsal part | Retrosplenial areas |
| Retrosplenial area, ventral part | Retrosplenial areas |
| Posterior parietal association areas | Posterior parietal areas |
| Anterior area | Posterior parietal areas |
| Rostrolateral visual area | Visual areas |
| Temporal association areas | Other isocortical areas |
| Perirhinal area | Other isocortical areas |
| Ectorhinal area | Other isocortical areas |
| Main olfactory bulb | Olfactory areas |
| Accessory olfactory bulb | Olfactory areas |
| Anterior olfactory nucleus | Olfactory areas |
| Taenia tecta, dorsal part | Olfactory areas |
| Taenia tecta, ventral part | Olfactory areas |
| Dorsal peduncular area | Olfactory areas |
| Piriform area | Olfactory areas |
| Nucleus of the lateral olfactory tract | Olfactory areas |
| Cortical amygdalar area, anterior part | Olfactory areas |
| Cortical amygdalar area, posterior part, lateral zone | Olfactory areas |
| Cortical amygdalar area, posterior part, medial zone | Olfactory areas |
| Piriform-amygdalar area | Olfactory areas |
| Postpiriform transition area | Olfactory areas |
| Field CA1 | Hippocampal region |
| Field CA2 | Hippocampal region |
| Field CA3 | Hippocampal region |
| Dentate gyrus | Hippocampal region |
| Fasciola cinerea | Hippocampal region |
| Induseum griseum | Hippocampal region |
| Entorhinal area, lateral part | Retrohippocampal region |
| Entorhinal area, medial part | Retrohippocampal region |
| Parasubiculum | Retrohippocampal region |
| Postsubiculum | Retrohippocampal region |
| Presubiculum | Retrohippocampal region |
| Subiculum | Retrohippocampal region |
| Prosubiculum | Retrohippocampal region |
| Hippocampo-amygdalar transition area | Retrohippocampal region |
| Area prostriata | Retrohippocampal region |
| Claustrum | Cortical subplate |
| Endopiriform nucleus, dorsal part | Cortical subplate |
| Endopiriform nucleus, ventral part | Cortical subplate |
| Lateral amygdalar nucleus | Cortical subplate |
| Basolateral amygdalar nucleus | Cortical subplate |
| Basomedial amygdalar nucleus | Cortical subplate |
| Posterior amygdalar nucleus | Cortical subplate |
| Striatum, unspecified | Striatum |
| Caudoputamen | Striatum |
| Nucleus accumbens | Striatum |
| Fundus of striatum | Striatum |
| Olfactory tubercle | Striatum |
| Lateral septal nucleus | Striatum |
| Septofimbrial nucleus | Striatum |
| Septohippocampal nucleus | Striatum |
| Anterior amygdalar area | Striatum-like amygdalar areas |
| Bed nucleus of the accessory olfactory tract | Striatum-like amygdalar areas |
| Central amygdalar nucleus, capsular part | Striatum-like amygdalar areas |
| Central amygdalar nucleus, lateral part | Striatum-like amygdalar areas |
| Central amygdalar nucleus, medial part | Striatum-like amygdalar areas |
| Intercalated amygdalar nucleus | Striatum-like amygdalar areas |
| Medial amygdalar nucleus | Striatum-like amygdalar areas |
| Striatum-like amygdalar nuclei | Striatum-like amygdalar areas |
| Pallidum, unspecified | Pallidum |
| Globus pallidus, external segment | Pallidum |
| Globus pallidus, internal segment | Pallidum |
| Substantia innominata | Pallidum |
| Magnocellular nucleus | Pallidum |
| Medial septal nucleus | Pallidum |
| Diagonal band nucleus | Pallidum |
| Triangular nucleus of septum | Pallidum |
| Bed nuclei of the stria terminalis | Pallidum |
| Thalamus, unspecified | Thalamus, sensory-motor cortex related |
| Ventral anterior-lateral complex of the thalamus | Thalamus, sensory-motor cortex related |
| Ventral medial nucleus of the thalamus | Thalamus, sensory-motor cortex related |
| Ventral posterior complex of the thalamus | Thalamus, sensory-motor cortex related |
| Posterior triangular thalamic nucleus | Thalamus, sensory-motor cortex related |
| Subparafascicular nucleus | Thalamus, sensory-motor cortex related |
| Subparafascicular area | Thalamus, sensory-motor cortex related |
| Peripeduncular area | Thalamus, sensory-motor cortex related |
| Medial geniculate complex | Thalamus, sensory-motor cortex related |
| Dorsal part of the lateral geniculate complex | Thalamus, sensory-motor cortex related |
| Lateral posterior nucleus of the thalamus | Thalamus, polymodal association cortex related |
| Posterior complex of the thalamus | Thalamus, polymodal association cortex related |
| Posterior limiting nucleus of the thalamus | Thalamus, polymodal association cortex related |
| Suprageniculate nucleus | Thalamus, polymodal association cortex related |
| Ethmoid and retroethmoid nucleus | Thalamus, polymodal association cortex related |
| Anteroventral nucleus of thalamus | Thalamus, polymodal association cortex related |
| Anteromedial nucleus (including the interanteromedial and interanterodorsal nucleus) | Thalamus, polymodal association cortex related |
| Anterodorsal nucleus | Thalamus, polymodal association cortex related |
| Lateral dorsal nucleus of thalamus | Thalamus, polymodal association cortex related |
| Intermediodorsal nucleus of the thalamus | Thalamus, polymodal association cortex related |
| Mediodorsal nucleus of thalamus | Thalamus, polymodal association cortex related |
| Perireuniens nucleus | Thalamus, polymodal association cortex related |
| Paraventricular nucleus of the thalamus | Thalamus, polymodal association cortex related |
| Parataenial nucleus | Thalamus, polymodal association cortex related |
| Nucleus of reuniens | Thalamus, polymodal association cortex related |
| Xiphoid thalamic nucleus | Thalamus, polymodal association cortex related |
| Central nuclei | Thalamus, polymodal association cortex related |
| Rhomboid nucleus | Thalamus, polymodal association cortex related |
| Parafascicular nucleus | Thalamus, polymodal association cortex related |
| Posterior intralaminar thalamic nucleus | Thalamus, polymodal association cortex related |
| Reticular nucleus of the thalamus | Thalamus, polymodal association cortex related |
| Intergeniculate leaflet of the lateral geniculate complex | Thalamus, polymodal association cortex related |
| Intermediate geniculate nucleus | Thalamus, polymodal association cortex related |
| Ventral part of the lateral geniculate complex | Thalamus, polymodal association cortex related |
| Subgeniculate nucleus | Thalamus, polymodal association cortex related |
| Medial habenula | Thalamus, polymodal association cortex related |
| Lateral habenula | Thalamus, polymodal association cortex related |
| Supraoptic group | Hypothalamus, other |
| Paraventricular hypothalamic nucleus | Hypothalamus, other |
| Periventricular zone | Hypothalamus, other |
| Arcuate hypothalamic nucleus | Hypothalamus, other |
| Dorsomedial nucleus of the hypothalamus | Hypothalamus, other |
| Preoptic area | Hypothalamus, other |
| Hypothalamus, other | Hypothalamus, other |
| Parastrial nucleus | Hypothalamus, other |
| Subparaventricular zone | Hypothalamus, other |
| Suprachiasmatic nucleus | Hypothalamus, other |
| Anterior hypothalamic nucleus | Hypothalamic medial zone |
| Lateral mammillary nucleus | Hypothalamic medial zone |
| Medial mammillary nucleus | Hypothalamic medial zone |
| Supramammillary nucleus | Hypothalamic medial zone |
| Tuberomammillary nucleus | Hypothalamic medial zone |
| Medial preoptic nucleus | Hypothalamic medial zone |
| Dorsal premammillary nucleus | Hypothalamic medial zone |
| Ventral premammillary nucleus | Hypothalamic medial zone |
| Paraventricular hypothalamic nucleus, descending division | Hypothalamic medial zone |
| Ventromedial hypothalamic nucleus | Hypothalamic medial zone |
| Posterior hypothalamic nucleus | Hypothalamic medial zone |
| Lateral hypothalamic area | Hypothalamic lateral zone |
| Lateral preoptic area | Hypothalamic lateral zone |
| Parasubthalamic and preparasubthalamic nucleus | Hypothalamic lateral zone |
| Perifornical nucleus | Hypothalamic lateral zone |
| Retrochiasmatic area | Hypothalamic lateral zone |
| Subthalamic nucleus | Hypothalamic lateral zone |
| Tuberal nucleus | Hypothalamic lateral zone |
| Zona incerta | Hypothalamic lateral zone |
| Fields of Forel | Hypothalamic lateral zone |
| Superior colliculus, sensory related | Midbrain |
| Inferior colliculus | Midbrain |
| Nucleus of the brachium of the inferior colliculus | Midbrain |
| Nucleus sagulum | Midbrain |
| Parabigeminal nucleus | Midbrain |
| Midbrain trigeminal nucleus | Midbrain |
| Subcommissural organ | Midbrain |
| Substantia nigra, reticular part | Midbrain |
| Substantia nigra, compact part | Midbrain |
| Ventral tegmental area | Midbrain |
| Paranigral area | Midbrain |
| Midbrain reticular nucleus | Midbrain |
| Midbrain reticular nucleus, retrorubral area | Midbrain |
| Superior colliculus, motor related | Midbrain |
| Periaqueductal grey | Midbrain |
| Olivary pretectal nucleus | Midbrain |
| Pretectal region | Midbrain |
| Cuneiform nucleus | Midbrain |
| Red nucleus | Midbrain |
| Midbrain, motor related, other | Midbrain |
| Pedunculopontine nucleus | Midbrain |
| Interpeduncular nucleus | Midbrain |
| Midbrain raphe nuclei, other | Midbrain |
| Pons, sensory related | Pons |
| Pons, motor related | Pons |
| Pons, behavioral state related | Pons |
| Medulla, unassigned | Medulla |
| Medulla, sensory related | Medulla |
| Medulla, motor related | Medulla |
| Medulla, behavioral state related | Medulla |
| Cerebellar cortex | Cerebellum |
| Cerebellar nuclei | Cerebellum |
| Fiber tracts | Fiber tracts |
| Ventricular systems | Ventricular system |

**Table S14. Hierarchical groups used for analyses.** Custom regions terms used to group regions in nutil are shown in the right column; the major hierarchical region assigned for each custom region is shown in the left column.

| **Region** | **Age** | **Sex** | **Receptor** | **Skewness** | **Kurtosis** | **Animal ID** |
| --- | --- | --- | --- | --- | --- | --- |
| Motor areas | P49 | Female | D2 | 2.127 | 4.611 | D166 |
|  | P70 | Male | D1 | -1.960 | 4.054 | C19 |
| Somatosensory areas | P70 | Female | D1 | 2.120 | 4.710 | E15 |
| Gustatory and visceral areas | P25 | Male | D2 | 1.972 | 3.901 | D235 |
| Anterior cingulate areas | P70 | Male | D2 | 2.179 | 4.809 | D123 |
| Prefrontal areas | P49 | Female | D1 | -1.835 | 3.547 | C68 |
|  | P49 | Male | D1 | 2.238 | 5.394 | C53 |
| Retrosplenial areas | P35 | Male | D1 | -1.959 | 4.398 | C108 |
| Olfactory areas | P70 | Female | D2 | -1.941 | 3.799 | D143 |
| Hippocampal region | P70 | Male | D1 | 1.951 | 3.914 | C17 |
| Striatum-like amygdalar areas | P49 | Male | D1 | 2.235 | 5.108 | C45 |
| Pallidum | P49 | Male | D2 | -1.843 | 3.515 | D157 |
| Thalamus, sensory-motor cortex related | P17 | Female | D1 | 1.845 | 4.147 | D13 |
|  | P25 | Female | D1 | 2.333 | 5.799 | E62 |
| Thalamus, polymodal association cortex related | P25 | Female | D1 | 2.267 | 5.482 | E62 |
| Hypothalamic medial zone | P25 | Female | D1 | 2.293 | 5.539 | E62 |
|  | P35 | Male | D2 | 1.996 | 3.985 | D197 |
|  | P49 | Male | D1 | 1.682 | 3.854 | C45 |
|  | P70 | Male | D1 | 2.041 | 4.404 | C17 |
|  | P70 | Male | D2 | 1.919 | 3.744 | D146 |
| Hypothalamic lateral zone | P35 | Male | D2 | 1.976 | 3.915 | D197 |
|  | P70 | Male | D2 | 2.035 | 4.254 | D146 |

**Table S15.** Skewness and Kurtosis values for any group with a statistical outlier. These values were excluded from statistical analyses. P = postnatal day.
